# Advancing Nano-Flow Cytometry: High-Precision Sorting and Analysis of Extracellular Vesicles

**DOI:** 10.64898/2026.07.27.741001

**Authors:** Irene Castrosín, Vanessa Costa, Brandy Pickney, Ionita Ghiran, Kieran Brennan, Flor Delgado, César Reyes-Pérez, K Jensen, Alfonso Blanco, John Tigges, Margaret Mc Gee

**Affiliations:** School of Biomolecular & Biomedical Science, University College Dublin (UCD), Belfield, Dublin, 4, Ireland; Flow Cytometry Core and Center for Nanoparticle Research (RRID:SCR_012305), Beth Israel Deaconess Medical Center, Harvard Medical School, Boston, MA, United States of America; Department of Anesthesia, Beth Israel Deaconess Medical Center, Harvard Medical School, Boston, MA, United States of America; UCD Conway Flow Cytometry Core, Conway Institute, University College Dublin (UCD), Dublin, Ireland; Harvard Medical School, Boston, MA, United States of America; Conway Institute of Biomolecular & Biomedical Research, University College Dublin (UCD), Dublin, Ireland

**Keywords:** Flow Cytometry, Extracellular Vesicles, Nanoflow Cytometry, nanoFACS, Particle sorting

## Abstract

Extracellular Vesicles (EVs) are small membrane-bound particles secreted by cells that play key roles in intercellular communication, gene regulation and modulation of cell function. They are involved in both physiological and pathological processes and, due to their ability to transport biomolecules across biological barriers, have emerged as promising tools for use as drug delivery vehicles and biomarkers with diagnostic and prognostic applications. Various methodologies are currently employed for the isolation, characterization, and analysis of EVs, including Ultracentrifugation (UC), Transmission Electron Microscopy (TEM), Nanoparticle Tracking Analysis (NTA), and Flow Cytometry. Flow Cytometry has emerged as a powerful technique capable of providing a multiparametric analysis of individual EVs. Recent advancements have led to the development of cytometers with higher sensitivity and increased limit of detection, enabling the detection and sorting of nanoscale particles—a technique known as Nano-Flow Cytometry. In this study, we show the optimization of small particle sorting, termed nanoFACS, via the CytoFLEX SRT. This method enables sorting based on size or fluorescence, enhancing reproducibility and broadening the potential for application in biological and clinical assays. Furthermore, we demonstrate the utility of nanoFACS in isolating nanoparticles from complex biofluids and in detecting miRNA using molecular beacons (MBs) highlighting its potential in both basic research and translational applications.

## 1. Introduction

Extracellular Vesicles (EVs) have gained significant attention due to their presence in all biological fluids and their ability to transport molecular cargo, regulate gene expression, and consequently alter the function of various cell types <u>(Simeoli R. et al, 2017; Cecchin R. et al, 2023)</u>. These characteristics make EVs an important focus of study, particularly with regard to their functional roles, molecular content, and cellular origin. Despite this growing interest, the characterization of EVs remains challenging due to limitations in current technologies and analytical techniques.

A range of traditional and well-established methodologies have been used for the analysis of these nanoparticles, including Ultracentrifugation, Size Exclusion Chromatography, Dark-field Microscopy, Western Blotting, Enzyme-Linked Immunosorbent Assay (ELISA), Real-Time Polymerase Chain Reaction (RT-PCR), Mass Spectrometry and Flow Cytometry (FCM) (Figure 1).

**Figure 1.**
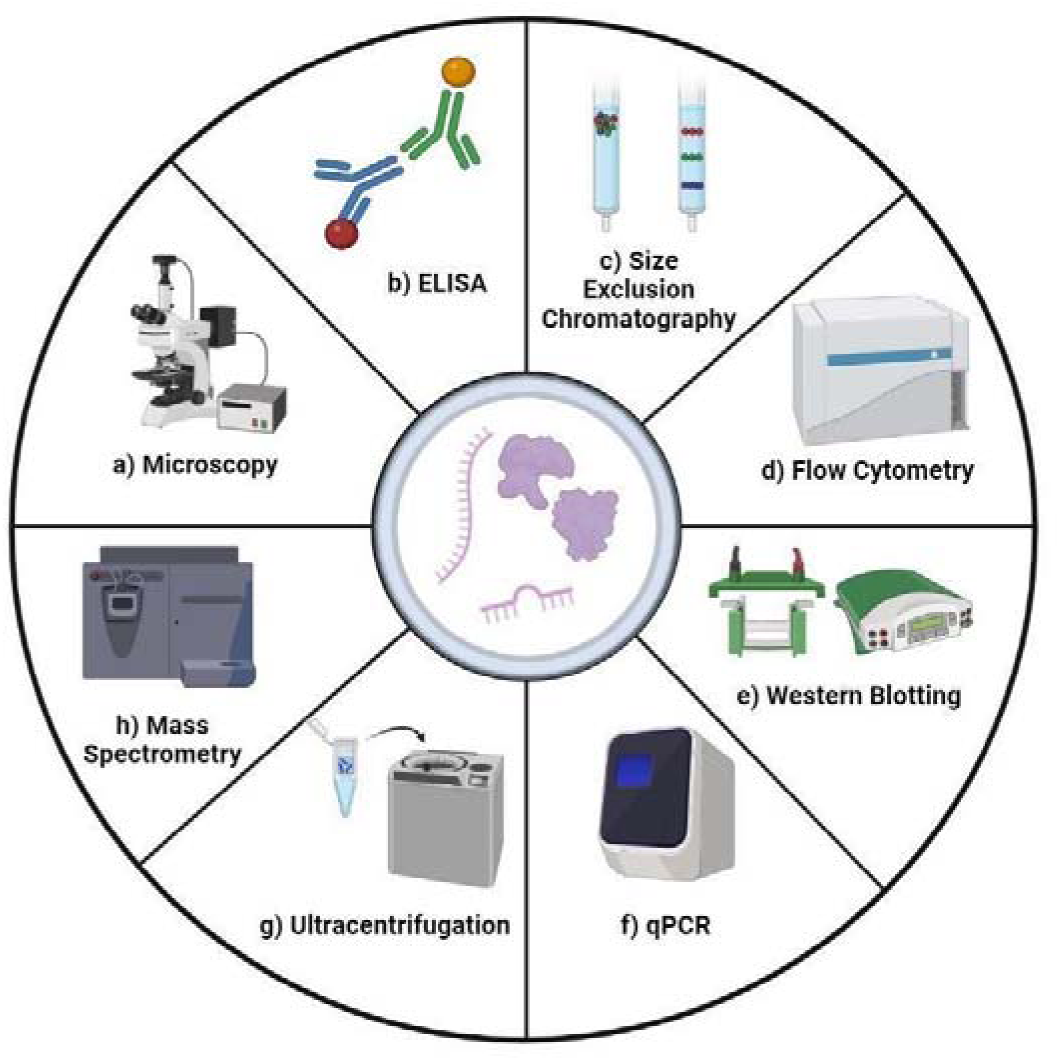
Schematic overview of techniques used for EV analysis. Adapted from <u>Zhang J et al., 2023</u>.

Nanoscale Flow Cytometry has emerged as a powerful tool for the characterization of Extracellular Vesicles (EVs), offering high-throughput, single-particle, and multiparametric analysis. The CytoFLEX S (Beckman Coulter, Indianapolis, IN) has previously demonstrated the ability to detect 70 nm polystyrene nanoparticles using the Violet Side Scatter parameter (VSSCH) (<u>Burnie et al., 2026</u>). Its enhanced sensitivity is attributed to advanced optical components, including Wavelength Division Multiplexing (WDM), Avalanche Photodiodes (APDs), and diode lasers. These engineering advancements significantly improve both light scatter and fluorescence sensitivity which has driven efforts to standardize nanoscale flow cytometry protocols, with the goal of improving rigor, reproducibility, and the accessibility of high throughput single particle characterization for broader applications, particularly in translational and clinical research.

Targeting and phenotyping EVs is achieved through Violet laser (405 nm) thresholding, fluorescent antibody labelling, and the use of calibration particles enabling precise identification and characterization based on size and surface markers. Traditionally, flow cytometric analysis results in sample loss, as particles are discarded into the waste stream after acquisition. However, the integration of cell sorting technology into nanoscale Flow Cytometry (nanoFACS), as originally described by <u>Morales-Kastresana et al. (2017</u>), addresses this limitation. By leveraging both size and fluorescence-based discrimination, nanoFACS enables the isolation of specific EV populations for downstream analyses. This capability is particularly valuable for studying EVs released under pathological conditions, such as those derived from cancer cells (<u>Wang et al., 2022</u>; <u>Maqsood et al., 2024</u>). Consequently, nanoFACS provides a promising platform not only for improved EV detection and isolation but also for advancing cancer diagnosis, prognosis, and the development of EV-based biomarker libraries and therapeutic delivery systems.

While Nanoscale Flow Cytometry continues to advance, the field of Nanoscale Fluorescence-Activated Cell Sorting (nanoFACS) remains relatively underexplored due to its inherent technical complexity. It has been previously demonstrated by <u>Morales-Kastresana et al., 2019</u> that the Beckman Coulter MoFlo Astrios EQ (Indianapolis, IN) is capable of sorting nanoparticles with an efficiency of 25-45%. However, the MoFlo Astrios EQ system has a large footprint and requires complex setup protocols limiting its use to highly skilled technicians. In contrast, the CytoFLEX SRT offers a compact design and automated sort setup, minimizing instrument alignment requirements and enabling more standardized and reproducible nanoFACS experiments. Building on the user-friendly software (CytExpert) interface and complimentary optical configuration of the CytoFLEX S, the CytoFLEX SRT presents a practical and accessible platform for nanoparticle sorting. To evaluate the feasibility and performance of the CytoFLEX SRT in nanoFACS applications, we conducted a series of experiments using standard reference materials and biologically relevant samples that closely resemble EVs (recombinant EVs, Mouse Leukemia Virus, Liposomes, Virus-like particles). These experiments assessed various detection strategies, including intrinsic fluorescence, fluorogenic dyes, and fluorescent antibody-based surface labeling comparing fluorescent versus scatter thresholding approaches.

To further validate this approach in biologically relevant samples, we first established the protein-based EV sorting strategy using EVs derived from cultured cells. Specifically, we analyzed EVs from SKBR3 cells, a well-established HER2-positive Human Adenocarcinoma cell line. SKBR3 cells are characterized by overexpression of the Human Epidermal Growth Factor Receptor 2 (HER2), and absence of Estrogen (ER) and Progesterone (PR) receptors, making them representative of the HER2-enriched Breast Cancer subtype. SKBR3 derived EVs also contain the EV marker CD9. Therefore, SKBR3 EVs were selectively targeted via the presence of HER2 and CD9, combined with fluorogenic lipid membrane dyes (Aco-490/Aco-600) for effective analysis and sorting.

After establishing the method in cell culture-derived EVs, we next evaluated its applicability in complex biological biofluids such as plasma. To do so, we first used previously characterized reference biological materials (Liposomes and Virus-Like Particles) and red blood cell EVs (RBC.EVs) (<u>Oliveira et al., 2023</u>) spiked into plasma, and demonstrated their successful re-isolation by sorting. Then, we combined both approaches by assessing the sorting of tumor-derived EVs directly in plasma. EVs derived from HeLa cells were spiked into plasma at a low concentration and subsequently sorted based on the expression of Epidermal Growth Factor

Receptor (EGFR) and CD147, two markers associated with cervical cancer progression and tumor invasiveness. This final experiment demonstrates the applicability of protein-targeted EV sorting for the selective isolation of tumor-derived EVs in complex biological environments.

Finally, we extended the methodology to RNA-based detection. Specifically, we used molecular beacons (MB) targeting miR-451a to isolate MB-positive EVs derived from red blood cells (RBC.EVs), thereby demonstrating the applicability of nanoFACS for both protein and RNA biomarker isolation. Collectively, these experiments highlight the capabilities of the CytoFLEX SRT to efficiently sort nanoparticles based on surface protein expression and nucleic acid content. Our findings validate the platform’s utility for high-precision nanoparticle isolation and reinforce its potential for biomarker discovery, diagnostic development, and therapeutic applications.

## 2. Materials and Methods

### 2.1. Flow Cytometry Equipment and Instrument Setup

The Beckman Coulter CytoFLEX SRT (Indianapolis, IN) was used to analyze and sort, respectively, all biological and non-biological materials. The instrument is equipped with 405 nm, 488 nm, 561 nm and 638 nm lasers and detects up to 13 fluorescence parameters. Instrument configurations listed in Supplemental (Table S2).

Daily startup and QC were performed following manufacturer’s description within the instructions for use (IFU). In brief, the automated system start-up program was performed. After start-up was completed, instrument QC was performed using CytoFLEX Daily QC fluorospheres (REF# C65719; Beckman Coulter, Indianapolis IN). Upon completion of the instrument QC, the instrument was configured for VSSC detection as described in (<u>Brittain et al., 2019</u>). The 405/10 filter and 450 filter positions were swapped, thus eliminating the Pacific Blue channel and creating the Violet Side Scatter Height (VSSC) channel. To add an additional Red Side Scatter (RSSC) for some experiments, the 710/50 BP was replaced with the 638/10 RSSC. Multi-side scatter (Violet, Blue and Red SSC) acquisition was used to help to identify subpopulations of sub-micron particles, as it leverages wavelength-dependent scattering to better discriminate particle populations based on their unique optical scatter signatures and not only on size or fluorescence. Both automated drop delay and sort calibration were performed via the automated processes within the SRT CytExpert software (Beckman Coulter, Indianapolis IN). A general gating strategy set up for EV detection can be found on Figure S1, Supplemental.

### 2.2. Microscopy

Microscopy images were taken with an Olympus BX62 microscope (Olympus Corporation, Waltham, MA, USA). EV images were acquired using double immersion dark field and fluorescence microscopy as described previously. Images were captured using an Andor Sona EMCCD camera at 40×1.00 NA iris diaphragm UPlan Apo objective. Both the FITC single-channel and Texas Red single channel were used with the exposure set to 100 ms.

### 2.3. Preparation and Use of NIST Beads

NIST Beads (ThermoFisher, Fremont, CA) are polystyrene spheres with sizes ranging from 70 to 495 nm. The spheres have a refractive index of 1.59 at 589 nm (25°C) and a density of 1.05 g/cm^3^ (further details are provided in Table S1, Supplementary). Beads were prepared as described by Welsh and Jones in FCM_PASS_ - Acquisition and gating of light scatter reference materials V.2 (<u>Welsh and Jones., 2020</u>). Briefly, each stock bottle was mixed by rapid inversion, vortexed for 2 mins, and sonicated for 1 min to maximize separation of bead particles and one drop was aliquoted. 10 µL of the droplet was serially diluted using clean, filtered DPBS to a concentration of ∼1×10^7^. Beads were then acquired at the lowest flow rate and 10,000 events were recorded in the designated gate using Violet Side Scatter configuration on the CytoFLEX S and SRT. The gating strategy and analysis of these beads on each instrument are detailed in the Supplementary Materials (Fig S2, S3 and S4). Subsequently, the Median VSSC-H values recorded for each bead size were used to perform a scatter cross calibration using FCM_PASS_ software. This scatter calibration was applied to convert the median VSSC-H values of the particles of interest to estimated particle diameters (The nominal diameter in nm) based on Mie theory, while accounting for the refractive index of the target population.

### 2.4. MESF Calibration Beads Preparation and Acquisition

Fluorescence calibration was performed using FITC-5 MESF beads (Cat# 823B; Bangs Laboratories, Inc., Fishers, IN), BD Quantibrite™ Beads PE Fluorescence Quantitation Kit (Cat# 340495; Becton, Dickinson and Company, BD Biosciences, CA), and 8-peak rainbow beads (Cat# 559123; Spherotech Inc., Lake Forest, IL). The 8-peak rainbow beads contain eight different fluorescence intensities across all channels and were primarily used for voltage optimization (voltration) to adjust the instrument settings for fluorescence measurements. The FITC-5 MESF (7-9 µm) beads consist of five populations with varying FITC intensities, and the PE MESF beads (3.6 µm) contain four populations with different PE levels. Both MESF bead sets, along with the 8-peak rainbow beads, were used to perform cross calibration via FCM_PASS_.

FITC-5 MESF, PE MESF and 8-peak beads were prepared and acquired as described in FCM_PASS_ - Acquisition and gating of fluorescence reference materials. Briefly, one drop of the FITC-5 MESF and 8-peak beads, and the reconstituted PE MESF beads, were added to 250 µl of DPBS into separate tubes respectively. Beads were then acquired at the lowest flow rate and 10,000 events were recorded in the designated bead gate using small particle analysis settings. The analysis of these calibration beads on each instrument are detailed in the Supplementary Materials (Fig S5). The median fluorescence intensity signals for FITC and PE from the FITC-5 MESF and PE MESF beads were cross-calibrated with the 8-peak rainbow beads via FCM_PASS_ software.

### 2.5. Preparation and Use of ApogeeMix Sizing Beads

For the sorting of HeLa-derived EVs another set of beads was used for the set-up of the CytoFLEX SRT. ApogeeMix #1527 Beads (Apogee, Catalogne, Spain) are a mix of fluorescent polystyrene (refractive index IZ=1.59) and non-fluorescent silica beads (refractive index IZ=1.43) with sizes ranging from 80 nm to 1300 nm diameter. Beads were mixed by rapid inversion and mixing 100 µl of the ApogeeMix with 100 µl of filtered fresh PBS. Beads were then acquired at the lowest flow rate and events were recorded for 2 min. Then, in a bivariate density plot of VSSC versus RSSC, an “Apogee” gate was generated to select only those events ranging between 80 and 500 nm beads.

### 2.6. Fluorescent Microspheres

100 nm and 200 nm Fluoresbrite YG Microspheres (Polysciences, Inc. Warrington, PA) are internally dyed microspheres with narrow FL CVs. YG microspheres (λEx 441 nm; λEm 486 nm) have an Ex/Em similar to that of FITC (λEx 488 nm; λEm 513-530 nm) and can be observed with a 525BP. For sorting experiments, 100 nm and 200 nm Fluoresbrite YG microspheres (4.55 x 10^13^ particles/mL, RI 1.59) were serially diluted in filtered DPBS to a concentration of ∼1×10^7^particles/mL. Subsequently, the two bead populations were mixed at a 1:1 ratio.

### 2.7. Virus Like Particles (VLPs)

Virus Like Particles (VLPs) are non-infectious particles that mimic the structure of a virus but lack its genetic material. HIV VLPs tagged with two fluorescent proteins: eGFP (λEx 488 nm, λEm 513 nm) and tdTomato (λEx 554 nm, λEm 581 nm), have a size of 100-120 nm and a refractive index (RI) of 1.369. Due to their well-defined characteristics, the VLPs serve as reliable biological reference material for standardization. eGFP and tdTomato tagged VLPs were prepared as described by <u>Gummuluru, 2012; 2013</u>. The expression plasmid HIV-1 pGag-eGFP and/or pGag-tdTomato which expresses a Gag-enhanced green fluorescent protein (eGFP) and/or tdTomato fluorescent fusion protein, has been described previously (Cat # ARP-11468 NIH HIV Reagent Program, Division of AIDS, NIAID, NIH; contributed by Marilyn D. Resh and George Pavlakis). In brief, HIV Gag-eGF/tdTomato VLPs were generated via calcium phosphate mediated transfections of HEK293T cells. Upon harvest, VLP-containing supernatant was cleared of cell debris by 10,000 x g for 15 minutes, passed through a 0.45-μM filter, and pelleted through a 20% sucrose cushion (20 mM HEPES [pH 7.4], 100 mM NaCl) and centrifuged at 200,000 × g for 2 h in a SW55 Ti rotor (Beckman). The p24gag content of VLPs was determined by ELISA.

### 2.8. Recombinant EVs (rEVs)

Recombinant EVs (rEVs) (Sigma-Aldrich, SAE0193-1VL) are small particles (50-180 nm, RI 1.37) expressing the green fluorescent protein (gag-EGFP) on their membrane. Details of size and structure are included in <u>Geeurickx et al., 2019</u>. These fluorescent standard particles were prepared according to the manufacturer’s instructions and were treated as biological reference material for sorting experiments.

### 2.9. Mouse Leukemia Virus (MLVs)

MLV-ViroFlow VLPs are non-infectious virus-like particles that mimic the murine leukemia virus structure without incorporating viral genetic material. Produced using the ViroFlow system (<u>Renner et al., 2020</u>), these particles are engineered to express a superfolded variant of GFP (sfGFP, ∼637 FITC equivalents) enabling robust fluorescence detection. MLVs constant fluorescence and well-defined size (Average Diameter of 120 nm) (<u>Welsh et al., 2020</u>) ensures their reliability as biological reference material for nanoFACS standardization and calibration.

### 2.10. Plasma preparation

Blood at BIDMC was obtained from healthy adult volunteers in accordance with the Declaration of Helsinki and approved by the Institutional Review Board of Beth Israel Deaconess Medical Center. Plasma preparation was performed following the protocol described by <u>Danielson et al., 2016</u>. Whole blood (WB) was first centrifuged at 300 × g for 20 min at room temperature, and the supernatant was collected and centrifuged again at 5000 × g for 10 min to obtain plasma. To generate platelet-poor plasma (PPP), plasma was further centrifuged at 2500 × g for 5 min at room temperature, and the supernatant was collected and centrifuged again at 12,500 × g for 10 min at room temperature. The resulting plasma was immediately stored at −80 °C until Extracellular Vesicle isolation and downstream analyses.

Blood donor plasma samples at UCD were obtained from the Irish Blood Transfusion Service (IBTS). Irish blood donors were asymptomatic and provided consent at donation for the use of their blood samples in anonymized research. Platelet free plasma (PFP) was obtained by centrifuging peripheral blood samples two times at 2500 × g at 4 °C, for 15 min. Samples were stored at −80 °C until EV isolation.

### 2.11. Red Blood Cell EVs (RBC.EVs)

#### 2.11.1. RBC purification

RBC.EVs were obtained as previously reported by <u>Ghiran et al., 2023</u>. The current study was approved by the Beth Israel Deaconess Medical Center Institutional Review Board (IRB #2001P000591). Fresh whole blood was obtained via venipuncture using Vacutainer EDTA tubes (BD, Franklin Lakes, NJ) from self-declared, healthy volunteers. Platelet-rich plasma was removed by centrifugation at 500 × g for 10 min. Blood was diluted in PBS, and leukocytes were removed by passing the blood through an Acrodisc WBC syringe filter (Pall Corporation, NY). RBCs were washed 3 times in PBS and centrifuged at 500 × g for 10 min.

#### 2.11.2. RBC EV generation and purification by SEC

Cells were incubated at 37°C for 15 min in a rotating block. Then, Ionomycin 10 μM (Streptomyces conglobatus, I9657, Sigma-Aldrich) or A23187 10 μM (C7522, Sigma-Aldrich) were added to the cells and incubated at 37°C for 1 h on a rotating block. Cells were centrifuged twice at 2500 × g for 15 min (brakes off), and the supernatant was concentrated to 500 μL using a 10 kDa Amicon® (Millipore Sigma). RBC.EVs were purified by loading the concentrate into a qEV 35 nm size-exclusion chromatography (SEC) column (Izon Science, Christchurch, New Zealand). Twenty 500 μL fractions were collected. For EV analysis, fractions 6–9 were pooled and concentrated to 100 μL using 100 kDa Amicon filter units.

### 2.12. Acoerela Dyes and Liposome standards

Acoerela fluorogenic conjugated oligoelectrolyte dyes (Aco490 and Aco600) were obtained from Acoerela (National University Singapore, Singapore). Aco490 (λEx 405 nm; λEm 458-508 nm) and Aco600 (λEx 488 or 561 nm; λEm 586-635 nm) were prepared via manufacturer’s specifications. Briefly, lyophilized dye was reconstituted in 100 µL filtered DPBS to obtain a 25 µM stock solution. Stock solution was vortexed for 2 mins followed by 1 min microfuge spin to pellet the dye. Next, 100 µL stock solution was added to 150 µL of filtered DPBS to obtain a 10 µM working solution.

Acoerela AcoRL Reference Liposomes (National University Singapore, Singapore) were obtained from Acoerela. AcoRL-490 (λEx 405 nm; λEm 458-508 nm) and AcoRL-600 (λEx 488 or 561 nm; λEm 586-635 nm) were chosen to correspond with the Aco dyes and verify instrument performance in the BV525 and YG610 channels. The lyophilized AcoRL standards have an average nominal diameter of 100 nm and mimic biological EV samples in RI. Lyophilized AcoRL 490 and 600 were reconstituted in 100 µL of filtered DPBS, vortexed for 5 mins, and centrifuged in a microfuge for 2 mins. A serial dilution was performed using filtered DPBS to obtain ∼1×106 particles/ml from 1×1012 particles/mL stock.

### 2.13. Labeling of RBC.EVs, VLPs, rEVs, and plasma with Aco dyes

Samples were labeled with either Aco490 or Aco600 according to manufacturer’s specifications. 5 µL of 10 µM Aco dye was added to 45 µL of ∼1×10^10^ biological samples. Samples were incubated at RT for 1h. After incubation, 10 µL of stained samples were added to 990 µL of filtered DPBS. Samples were then serially diluted to assess optimal concentration to acquire on the CytoFLEX SRT (∼1×10^7^).

### 2.14. Preparation of SKBR3 and HeLa EVs

#### 2.14.1. Cells culture

SKBR3 human breast adenocarcinoma cells (ATTC, HTB-30) display epithelial morphology, have a size of 17-22 μm and grow as adherent layers with a doubling time of 70 hours. SKBR3 cells were grown in McCoy’s 5A medium, with sodium bicarbonate, without L-glutamine (SIGMA ALDRICH) and supplemented with 1% Pen/Strep, 1% L-glutamine and 10% fetal bovine serum (FBS). HeLa human cervical adenocarcinoma cells (ATTC, CCL-2) display epithelial morphology, have a size of 10-20 μm and grow as adherent layers with a doubling time of 24 hours. HeLa cells were grown in Eagle’s Minimum Essential Medium, with sodium bicarbonate, without L-glutamine (SIGMA ALDRICH) and supplemented with 1% Pen/Strep, 1% L-glutamine and 10% FBS.

#### 2.14.2. Preparation of conditioned media for HeLa and SKBR3 EV isolation

HeLa and SKBR3 cells at passages 15-20 were grown to confluence in complete media in T175 cell culture flasks at 37 °C under standard conditions. Cells were collected by centrifugation at 450 × g for 3 min. 2.5 × 10^6^ cells were cultured in 145/20 mm cell culture dishes with complete media. After 48 h, cells were attached to the culture dishes and the complete media was replaced by serum-free media. After 48 h, the conditioned media from the cultures was collected for EV isolation. Cells were removed by centrifugation at 450 × g for 3 min, and cell debris was removed by centrifugation at 2500 × g for 15 min. EV isolation was carried out immediately following conditioned media collection.

#### 2.14.3. HeLa and SKBR3 EV isolation by Iodixanol Density Gradient Ultracentrifugation

EVs were separated from soluble protein by Iodixanol density gradient ultracentrifugation, using a modified protocol from <u>Brennan et al. (2022)</u>. HeLa and SKBR3-derived cell culture conditioned media was centrifuged at 120,000 × g Avg (SW32Ti, Beckman Coulter (Miami, FL, USA), (brake = 9) for 2 h 10 min at 20 °C (centrifugation durations determined using a “50 nm cut-off size” as described in <u>Livshits et al. (2015)</u>. The pellet was washed in 1 mL PBS and centrifuged at 120,000 × g Avg for 35 min (MLA-130, Beckman Coulter, brake = 9). The EV-enriched pellet underwent a floatation density gradient for a high-purity EV isolation. A 54% iodixanol-PBS working solution (1.295 g/ml IPBS) was prepared by diluting a stock solution of OptiPrepTM (60% (w/v) aqueous iodixanol from Axis-Shield PoC, Norway) with 10X particle-free PBS (Gibco, Waltham, MA, USA), which was further diluted with 1x PBS to make a 1.2g/ml IPBS solution. The EV-enriched pellet was resuspended in 100 µL PBS and mixed with 336 µL of 54% iodixanol-PBS (IPBS). 500 µL 1.2 g/mL IPBS was overlaid on top of the sample layer followed by an overlay of 50 µL PBS, and the gradients were centrifuged at 120,000 ×g Avg for 15 h at 4 °C. The EV layer that formed on top of the 1.2 g/mL fraction was transferred to a new tube and diluted to less than 1.03 g/mL in PBS and centrifuged for 35 min at 20 °C (MLA-130, Beckman Coulter, brake = 9). The resulting EV-enriched pellet was resuspended in 100 μL PBS and stored at −80 °C.

#### 2.14.4. Antibody staining of SKBR3 and HeLa EVs

EVs were labelled with either anti-HER2-BV421 (5 µL/test, 756616 BD Biosciences), anti-CD9-PE (2 µL/test, 555372 BD Biosciences), anti-CD147-APC (0.12 µl/test, MEM-M6/1, Thermo Scientific) or EGFR-BV421 (0.2 µl/test, clone EMab-134, 749755 BD Bioscience) in 10 µl PBS for 30 min in the dark. To avoid false positive events, all antibodies were centrifuged before use at 20,000 g for 30 min to remove antibody aggregates. To avoid carry-over effects between each sample measurement, a washing step was performed with filtered double distilled water for 1 min at an increased flow rate of 60 μL/min. HeLa and SKBR3 EVs and plasma particles were quantified using the CytoFLEX S, as described in <u>Brennan et al. (2022</u>). SKBR3 EVs (1 × 10^8^ EVs/µL) were labelled with antibodies in 10 µL PBS for 30 min in the dark before being diluted to 1 mL with PBS to a EV concentration of 1 × 10^6^ EVs/µL for Flow Cytometry analysis and sorting. Staining controls for SKBR3 EVs are detailed in Supplementary Materials (Fig. S12).

HeLa EVs and plasma particles were diluted to achieve working stocks of HeLa EVs and plasma with a concentration of 6 × 10^6^ particles/µL. The HeLa EVs and plasma working stocks were combined in a ratio of 1:1, 1:3, 1:9, and 1:19 to make spiked plasma solutions. 1 µl Hela EVs, plasma, or spiked plasma solutions (6 × 10^6^ particles) were incubated with antibodies in 10 µL PBS for 30 min in the dark before being diluted to 200 μL with PBS prior to recording. For longer sortings (>15 mins) multiple samples were prepared at the same concentration, which were combined into one tube to provide enough sample for sorting.

### 2.15. Bead Based Molecular Beacons Assay

#### 2.15.1. Molecular Beacons and Target Sequences

The molecular beacons (MBs) and synthetic miRNA target analogs were obtained from Integrated DNA Technologies (IDT, Coralville, IA). The MBs were conjugated with a 5’ end 6-carboxyfluorescein (λEx 493 nm; λEm 517 nm) (FAM), and at the 3’ end an internal ZEN quencher (an internal dark quencher developed by Integrated DNA Technologies), followed by an 18-atom hexa-ethylene glycol spacer (ISp18), and biotin. The MBs were synthesized with the optimized stem sequence CGCGATC, as previously reported (<u>Ryazantsev et al., 2014</u>, <u>Oliveira GP Jr et al., 2020</u>). All the MBs and sequences used for the experiments are shown in Table 1.

**Table 1.**
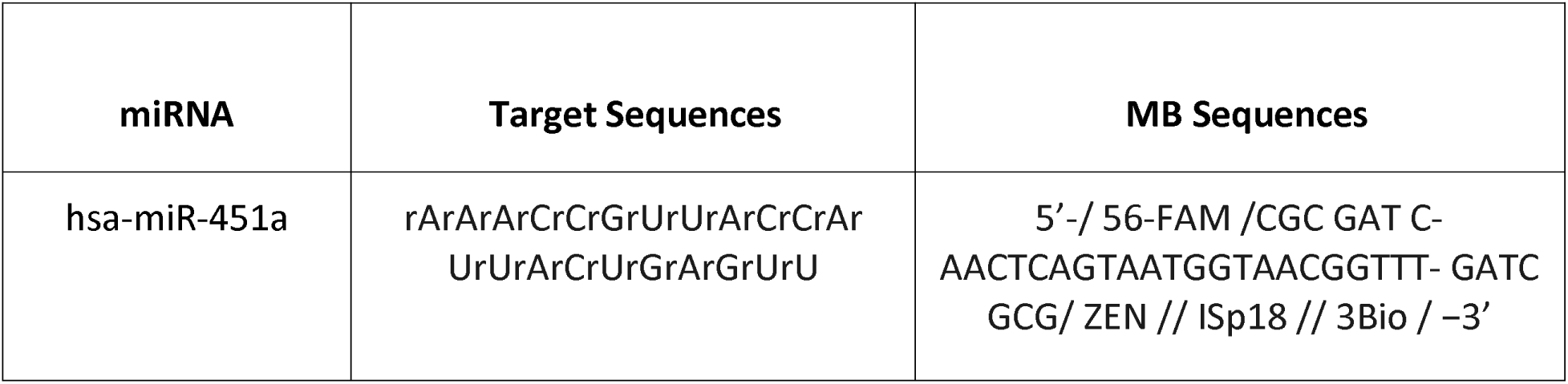
Molecular Beacons and corresponding target sequences.

#### 2.15.2. Streptavidin Beads

Streptavidin Coated Polystyrene Particles were obtained from Spherotech, Inc (REF#SVP-03-10). These particles are 192 nm beads with streptavidin protein that have similar binding properties of avidin, but with less non-specific binding. The particles were used to bind the MB and perform analysis and nano sorting of miRNA.

#### 2.15.3. Attachment of Molecular Beacons to Streptavidin Beads

The streptavidin beads were mixed with the biotinylated molecular beacons (MBBeads) in the 1:1 ratio and vortexed briefly. MBBeads were diluted in 500 µL of Dulbecco’s Phosphate Buffered Saline (DPBS) and incubated for 20 mins at room temperature. After incubation, a centrifugation of 20,000 x g was performed for 5 mins. The supernatant was removed, and another wash was used to remove aggregated beads. Once the wash was finalized, the pellet was resuspended in 500 µL of DPBS and was mixed with samples (RBC.EVs and Plasma).

#### 2.15.4. Staining of Plasma / RBC.EVs with MBBeads

An aliquot of plasma (500 µL) and RBC.EVs (50 µL) was lysed using MagIC lysis Buffer (ElementZero Biolabs, Berlin, Germany), in a 1:1 ratio for 10 mins at room temperature. After lysing, the biological samples were mixed with MBBeads and incubated for 40 mins at room temperature. The supernatant was removed, and the pellet was resuspended in 300 µL of DPBS and centrifugated at 20,000 x g for 5 mins. Then, the pellet was diluted in 500 µL of PBS and acquired on the flow cytometer.

## 3. Results

In this study, we describe a standardized technique for sorting small particles on the CytoFLEX SRT. To support this process, we incorporated FCM_PASS_, a specialized calibration software designed to enhance data consistency and instrument performance. This strategy allows for precise calibration of both light scattering and fluorescence signals, addressing a significant challenge in small particle analysis. The subsequent sections outline our key findings: first, we examine the impact of calibration on signal stability across various detection parameters; next, we assess the effectiveness of fluorescence-based nanoFACS under different threshold conditions; and finally, we demonstrate the use of this method for sorting biologically relevant samples. Specifically, we sorted HER2+ SKBR3 EVs from cell culture supernatant, and Liposomes, Virus-Like Particles (VLPs), Red Blood Cell EVs (RBC.EVs) and HeLa EVs spiked in plasma. These two sections exemplified membrane and protein-based sorting. In addition, we extend the methodology to RNA-based detection, with the sorting of miR-451a+ RBC.EVs in plasma.

### 3.1. Standardization using reference particles

#### 3.1.1 Light Scattering and Fluorescence Calibration

To ensure consistency in fluorescence and light scattering detection, we implemented a calibration strategy using FCM_PASS_ software. This approach allowed us to refine both scatter-and fluorescence-based detection, improving the comparability of results across different experiments and instruments. A summary of calibration reports for each instrument is available in the Supplementary Materials (Fig S6-S9).

For scatter calibration, NIST-traceable polystyrene beads were used to establish a reference for particle size estimation. This calibration provides insights into instrument performance and enables standardized data comparison across different flow cytometers. By importing the calibrated .fcs files into FlowJo software (v. 10.10), we calculated the nominal diameter of EVs and other small particles, improving the accuracy of size estimations and providing a reliable framework for nano flow analysis.

To standardize fluorescence intensity measurements, MESF standards were applied for PE and FITC, which enabled the conversion of fluorescence signals from arbitrary units to standard units. This normalization allows for direct comparison of fluorescence intensities across different instruments, minimizing variability due to differences in detector gain settings. FCM_PASS_ facilitated the calibration process by generating calibration curves and incorporating standardized values into the dataset, thereby ensuring reliable and comparable fluorescence measurements across all instruments.

In addition, to ensure consistent fluorescence measurements across experimental runs over time, we employed 8-peak Rainbow beads to assess detector performance. Through the cross-calibration process, FCM_PASS_ aligned the intensity distribution of the Rainbow beads with that of MESF beads, enabling direct fluorescence comparison across instruments while reducing the need for repeated MESF calibration.

#### 3.1.2 Sorting of beads

Initial sorting experiments were performed using NIST Traceable beads of 100, 125, and 300 nm diameter. By using well defined reference standards of equal properties, including RI, but differing only in size, one can assess the SRTs ability to sort based on VSSC. NIST beads are highly homogenous and monodisperse resulting in distinct populations which facilitates ease of gating of the three populations of interest.

By plotting VSSC-W vs VSSC-H, doublets, which are bead aggregates, are excluded from the analysis (Figure 2A). Sort logic was created for the three distinct singlet bead populations and sorted based on the VSSC threshold. Upon post sort analysis, increased background noise was observed as a result of droplet-mediated dilution of the sample. This occurs because the droplets generated by the CytoFLEX SRT are substantially larger than the nanoparticles being sorted in essence causing a dilution of the purified population resulting in a change in the signal to noise ratio. Consequently, the percentage of positive events and population statistics become skewed. Despite this, the sorting efficiency remained at 100% for all three populations (data not shown). Subsequent tests showed 300 nm beads were successfully separated from the 100 nm and 125 nm bead populations with similar efficiency. However, the percent purity does not reflect the sort efficiency. Moreover, although the populations were isolated from one another efficiently, the post sort statistics do not reflect the isolation efficiency due to the increased noise from the sheath fluid. This results in a lower bead signal to sheath noise ratio. These findings demonstrate the necessity of removing the fluid by concentrating the sorted populations prior to evaluating the sort purity (Figure 2B).

**Figure 2.**
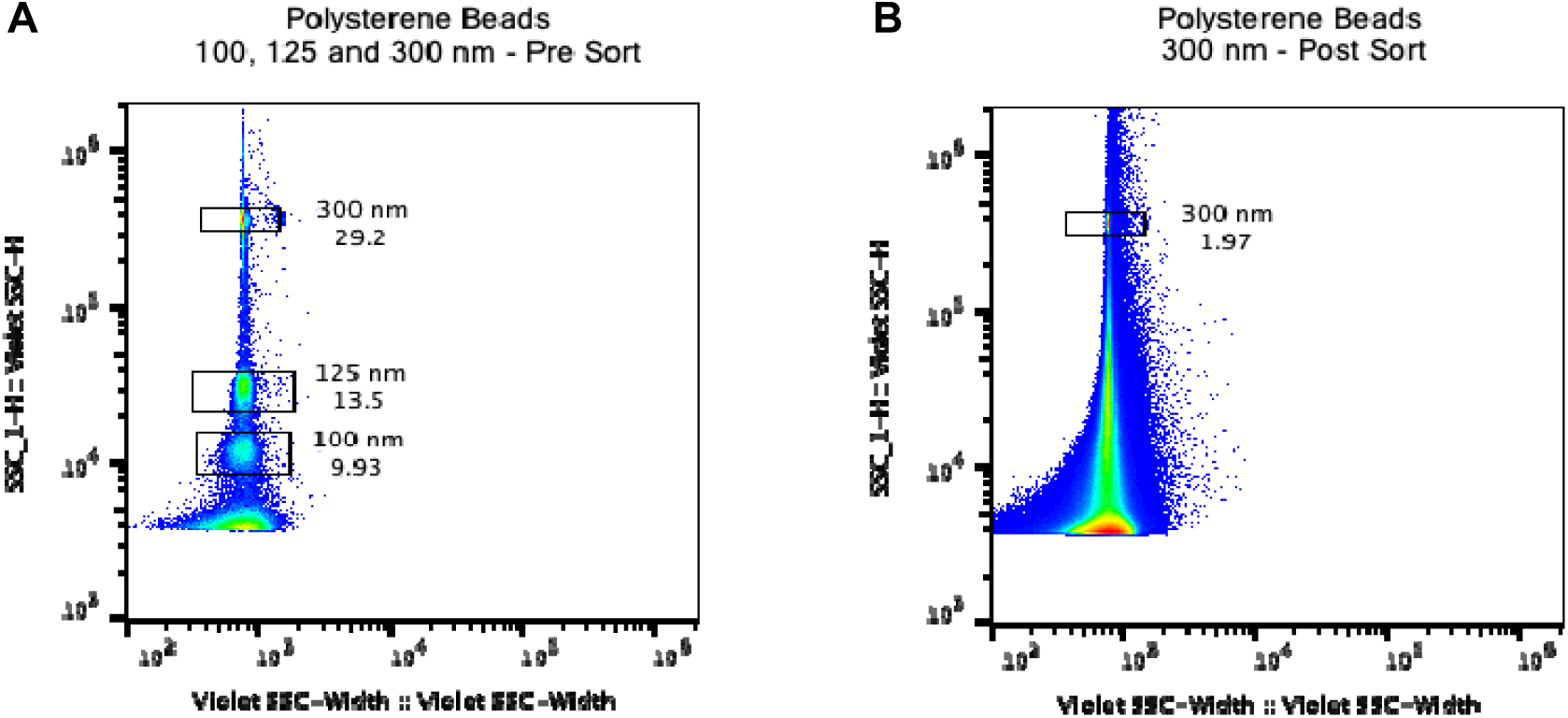
Separation based on size: 100, 125 and 300 nm mixed beads. NIST beads of varying sizes (100, 125 and 300 nm) were mixed and acquired using the CytoFLEX SRT. A) Separation of beads based on size and RI allowed for the simultaneous sorting of multiple bead populations and improved workflow efficiency. B) Despite droplet-mediated dilution of the sample, the particle size distribution of the beads remains consistent after sorting, although visualization of the population became more challenging due to the increased noise level.

To assess fluorescence (FL) sensitivity and the feasibility of sorting based on differences in Mean Fluorescence Intensity (MFI), 100 nm and 200 nm YG Fluoresbrite microspheres were employed. The threshold was established by evaluating the “noise” contribution from DPBS alone and adjusted to minimize noise contribution in post sort analysis. Furthermore, an increased number of events were sorted to alleviate the signal to noise ratio issue affecting sort purity statistics seen previously. Moreover, plotting VSSC vs B525A enabled clear delineation of the two bead populations based on size and FL intensity (Figure 3A) allowing accurate gating and sort logic. Figures 3B and 3C, outlines the post-sort analysis and demonstrates efficient separation of 100 nm and 200 nm YG beads, with purity of 91% and 85% respectively.

**Figure 3.**
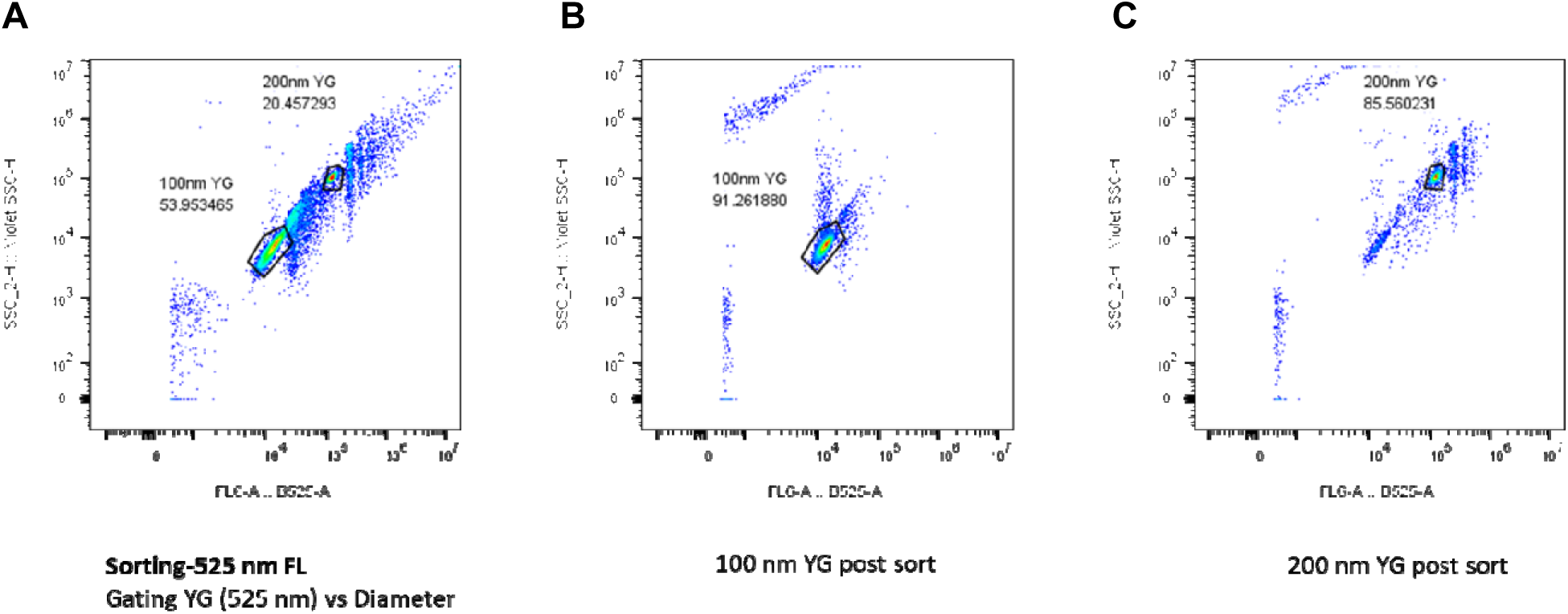
Fluorescence-based separation of 100 nm and 200 nm Yellow-Green beads using the CytoFLEX SRT. Yellow-green (YG) FL beads of two sizes (100 nm and 200 nm), exhibiting distinct fluorescence intensities (Ex 488 nm/Em 525 nm), were analyzed and sorted using the CytoFLEX SRT. Separation was performed based on differences in particle fluorescence intensity (A). Following sorting, each bead population was reanalyzed to evaluate the capability and effectiveness of fluorescence-based discrimination and sorting (B and C). The post-sort analysis confirms successful separation of the two fluorescent bead populations.

In addition, recombinant Extracellular Vesicles (rEVs) (nominal diameter 50-180 nm, RI 1.4, <u>Geeurickx et al., 2019</u>) were mixed with the 100 nm and 200 nm YG beads in a 1:1:1 ratio. Based on size distribution alone, 100 nm YG beads and rEVs occupy the same general gating area due to effects of RI (data not shown). Moreover, rEVs have characteristics that mimic EV materials and unknown samples need calibration to understand “true” sizing and particle characterization. Therefore, using the inherent differences in FL MFI, the three populations can be separated efficiently and sorted accordingly (Figure S10, Supplemental). Calibration of post sort samples show rEVs are larger in size compared to the 100nm YG beads, due to RI differences. Calibrated data shows rEVs are intermediate in size between 100 nm and 200 nm PS. These initial findings lead to the usage of FL parameters for sorting and threshold as FL is agnostic of RI changes. Furthermore, FL post sort analysis eliminates background noise through gating.

#### 3.1.3 Sorting of Fluorescent Reference Material and Comparison of Different Thresholds

Fluorescence-based sorting of small particles was evaluated using Virus-like particles (VLPs), Mouse Leukemia Virus (MLVs), and Recombinant EVs (rEVs) as model systems. VLPs labeled with eGFP or tdTomato served as well-characterized fluorescence controls, while MLVs (ViroFlow) and GFP-labeled rEVs, commercially available fluorescent reference material, were selected for their defined structural properties and predictable behavior in Flow Cytometry. Sorting strategies were designed based on fluorescence intensity, with MLV-derived VLPs sorted into two subpopulations, “GFP Low” and “GFP High”, without applying predefined MFI thresholds to assess fluorescence resolution (Figure 4A), whereas HIV gag-eGFP VLPs were arbitrarily classified into three subpopulations (“GFP Low,” “GFP Mid,” and “GFP High”) to model fluorescence variation akin to EVs with different epitope densities (Figure 4B).

**Figure 4.**
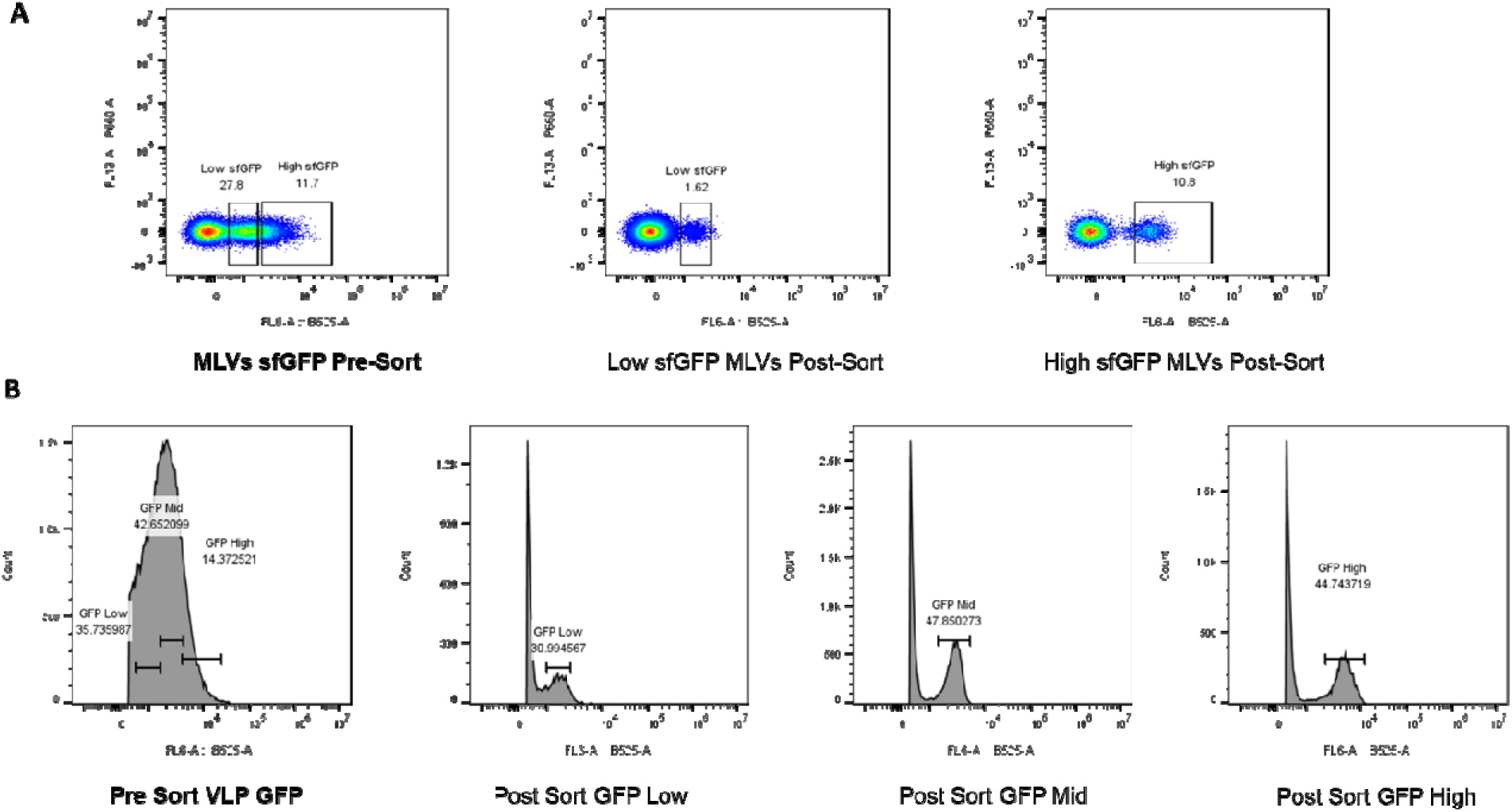
Particle sorting based on 525 nm fluorescence intensities. A) Sorting of Mouse Leukemia Virus (MLVs;-ViroFlow) particles by low and high sfGFP intensity. MLV-derived VLPs expressing sfGFP, with an average size of 124 ± 14 nm, were separated based on differential fluorescence intensity. A VSSC threshold was applied to eliminate background noise and singlet gating was used to ensure analysis of individual particles. The distinct separation of populations based on fluorescence intensity demonstrates the feasibility of sorting VLPs based on epitope abundance rather than size alone B) Sorting of HIV gag-eGFP VLPs into three fluorescence-defined subpopulations (“GFP Low,” “GFP Mid,” and “GFP High”) by three different fluorescent intensities. Particles were classified based on fluorescence intensity to mimic EV populations with varying epitope densities. The analysis demonstrates the feasibility of using fluorescence intensity as a robust parameter for subpopulation separation.

To further refine subpopulation discrimination, Aco490 labeling was introduced as an additional fluorescence parameter (Figure 5). As a fluorogenic lipophilic dye, Aco490 fluoresces upon interaction with lipid membranes, allowing for the differentiation of rEVs subpopulations based on both protein-associated and membrane-specific fluorescence signals This dual-parameter approach identified a GFP-Aco490 double-positive population as well as an Aco490-positive, GFP-negative population. Together, these results demonstrate the enhanced capability of Single Vesicle nanoFACS to resolve EV heterogeneity.

**Figure 5.**
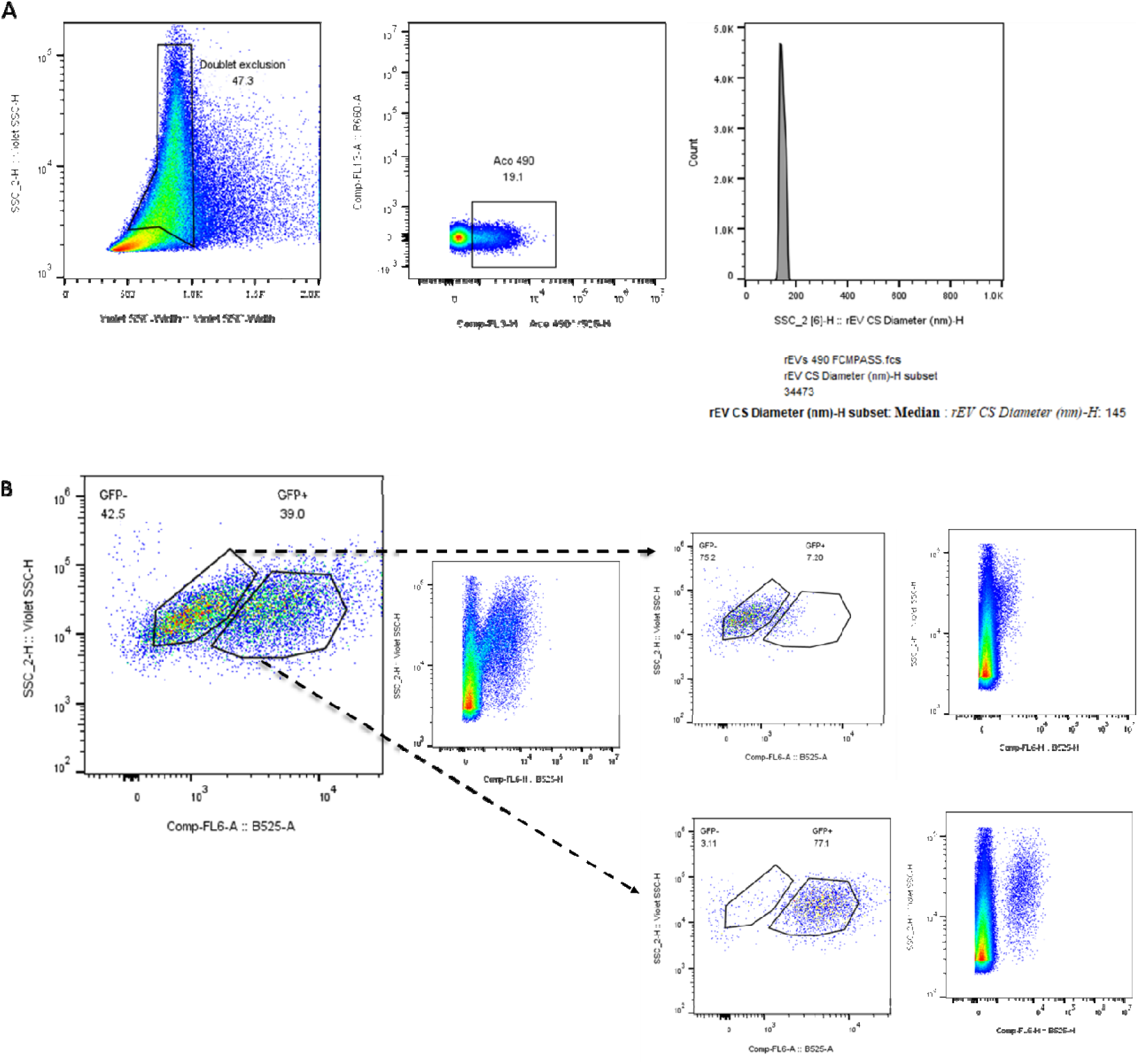
Sorting of labeled rEVs and HEK293T EVs. A) Standard rEVs were labeled with Aco490. After applying a doublet exclusion gate, Aco490+ events were selected (Aco490). B) Aco490+ rEVs were subsequently sorted based on their B525-GFP fluorescence (GFP- and GFP+). Post-sorting analysis confirmed successful isolation of GFP+ Aco490+ and GFP- Aco490+ rEVs using Single Vesicle nanoFACS, demonstrating the feasibility of sorting rEVs based on membrane labelling and the presence of intrinsic fluorescent proteins.

Due to hardware and software enhancements in the CytoFLEX platform, dual thresholding using VSSC and FL channel of interest can be implemented. This enables improved characterization of nanoparticles based on size and epitope density. By combining these parameters, distinct subpopulations can be identified without compromising the signal to noise (S/N) ratio; resulting in greater data accuracy and minimal contamination of noise related "false" negatives. To demonstrate this capability, the previously characterized rEVs were re-analyzed and sorted using combined VSSC and fluorescence-based thresholds (Figure 6). This strategy was subsequently extended to another well-characterized biological reference material: HIV gag-eGFP/tdTomato tagged VLPs (Figure 7). These samples were sorted with both VSSC and fluorescence thresholds, allowing for a direct comparison between size-based and fluorescence-based sorting approaches. Overall, this dual-threshold approach highlights how fluorescence-based strategies can refine nanoparticle isolation and characterization by maintaining signal-to-noise ratio accuracy and minimizing contamination from noise-related artifacts.

**Figure 6.**
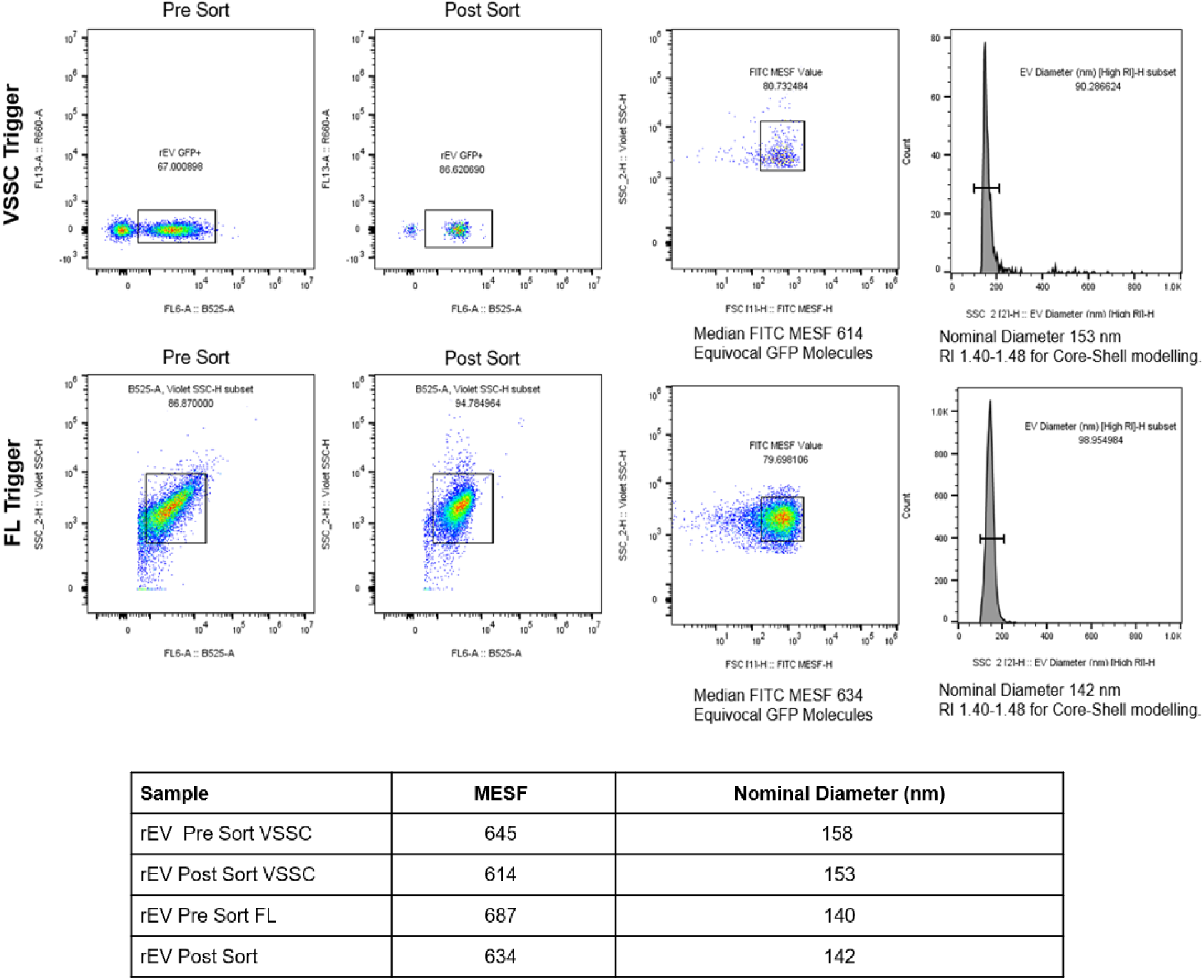
Comparison of VSSC and FL threshold for sorting of rEVs. rEVs-Sigma Aldrich are between 50-180 nm in diameter with a RI of 1.4 (<u>Geeurickx et al., 2019</u>), and a consistent GFP MESF signal for QA and assay reproducibility. The analysis also demonstrates that sorting does not affect EV statistics. The Nominal Diameter for Core-Shell modelling can be compared before and after sorting. By using VSSC, rEVs Pre-Sort is 158 nm and Post Sort 153 nm. By using FL threshold for sorting, rEVs Pre-Sort is 140 nm and it results in 142 nm Post Sort. Similarly, the Median FITC MESF corresponds to 614 Equivocal GFP Molecule for the post sorted rEVs by using VSSC threshold and to 634 Equivocal GFP Molecules for the post sorted rEVs by using FL threshold for sorting.

**Figure 7.**
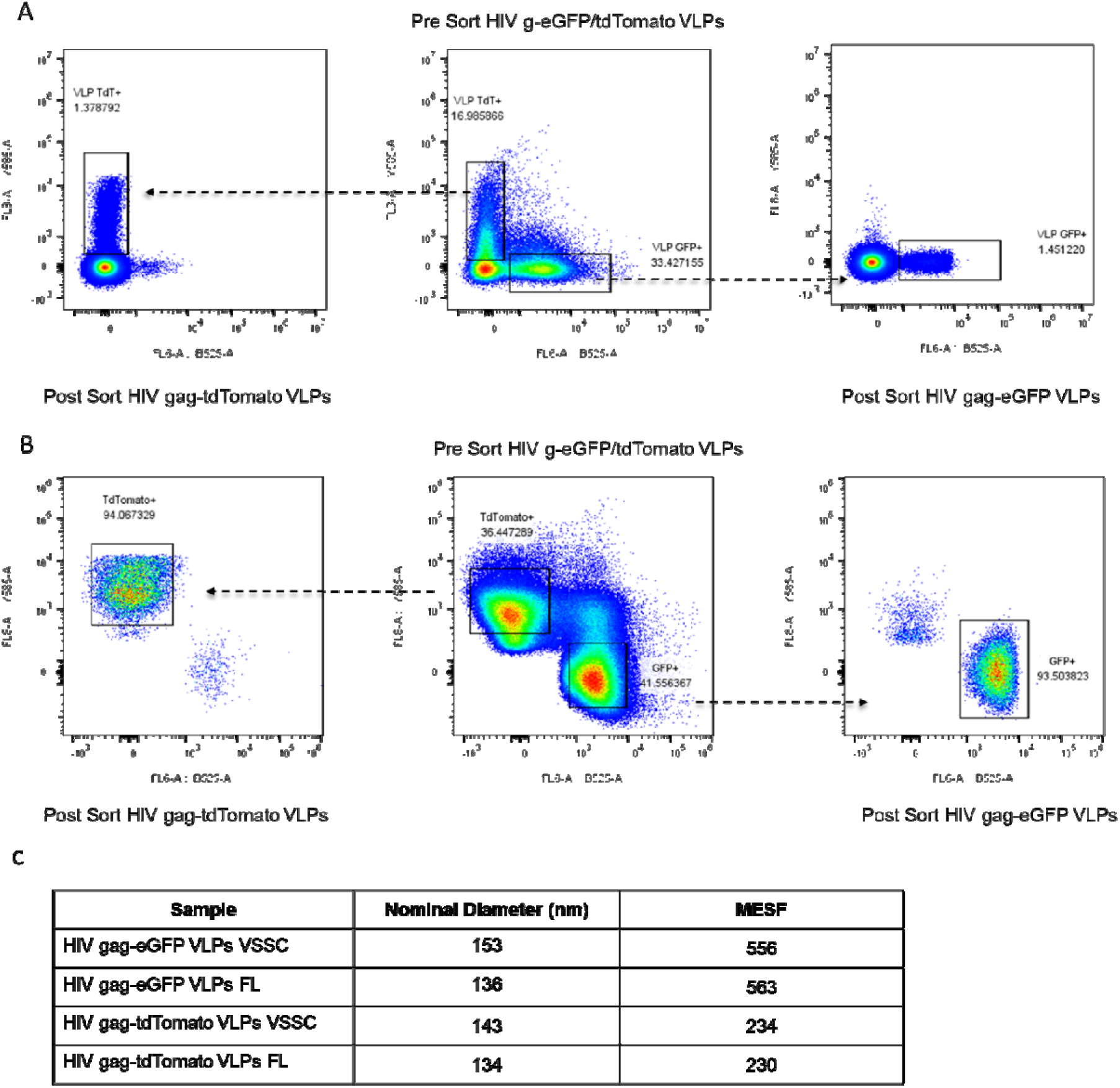
Sorting HIV gag-eGFP/tdTomato VLPs using VSSC or dual thresholding. A) Sort of HIV gag-eGFP/tdTomato VLPs using VSSC. B) Sort of HIV gag-eGFP/tdTomato VLPs using dual thresholding. CytoFLEX SRT enables thresholds to be applied on two parameters simultaneously. Applying a fluorescent threshold circumvents issues associated with "dark noise" as only FL+ events are registered. This enhances the signal to noise ratio (S/N) increasing the percentages of detected events, allowing for sort purity checks to be based on separation of populations, while minimizing false negatives caused by noise. C) The post-sort analysis of the HIV gag-GFP VLPs calibrated via FCM_PASS_, demonstrating consistent results between samples with nominal sizes of 153 nm and 136 nm, and MESF values of 556 and 563 for the sorting with VSSC and for the sorting with FL thresholds, respectively. These results confirm that the FL threshold enables the detection of smaller particles while epitope density remains unchanged, as the RI does not influence MFI/MESF calculations. Thus, the use of two thresholds improves detection accuracy and overall sorting reliability.

Furthermore, the advanced capabilities of the CytoFLEX platform, along with FCM_PASS_ standardization, ensure that EV statistics remain consistent before and after sorting independent of the threshold selected (Figures 6 and 7). This robust framework not only enhances the reliability and precision of measurements but also allows for cross-instrument comparisons to be performed. For instance, the sorting of HIV gag-eGFP/TdTomato tagged VLPs and calibration analysis via FCM_PASS_ across two institutions (UCD and BIDMC) demonstrates that this standardization facilitates direct comparisons between instruments (Figure S11, Supplemental), thereby highlighting the improved reliability and comparability of EV data in multi-institutional studies.

### 3.2. Sorting of SKBR3 HER2 positive EVs from cell culture supernatant

SKBR3 EVs were characterized by FCM_PASS_ Core-Shell modelling, which estimated their diameter at ∼145 nm (RI 1.40–1.48). Membrane detection was performed with Acoerela fluorogenic dyes, while surface proteins were assessed using antibodies against HER2 and several tetraspanins (CD9, CD63, CD81, and a combined panel of the three). Based on tetraspanin analyses, CD9 was selected as the reference tetraspanin. This combined evaluation of size, membrane, and protein markers established the foundation for accurate identification and subsequent sorting of HER2-positive SKBR3 EVs. Further details are provided in the supplementary section (Figure S12, Supplemental). The next step was to develop a strategy for sorting SKBR3 EVs with the CytoFLEX SRT. The first approach involved sorting HER2-positive SKBR3 EVs followed by post sort staining and analysis of the collected particles (Figure 8A). Single HER2-positive EVs were successfully isolated and post-sort staining with Aco600 confirmed their membranous nature while discriminating against antibody aggregates. The second approach consisted of the sorting of triple positive SKBR3 EVs (Aco490-positive, HER2-positive, and CD9-positive) using a gating strategy set up to identify single positive Aco490+, HER2+, and CD9+ SKBR3 EVs individually with the three gates being combined to identify triple positive SKBR3 EVs. Additionally, a FL threshold was employed during the analysis of the triple positive sorted population to ensure precise detection and characterization (Figure 8B).

**Figure 8.**
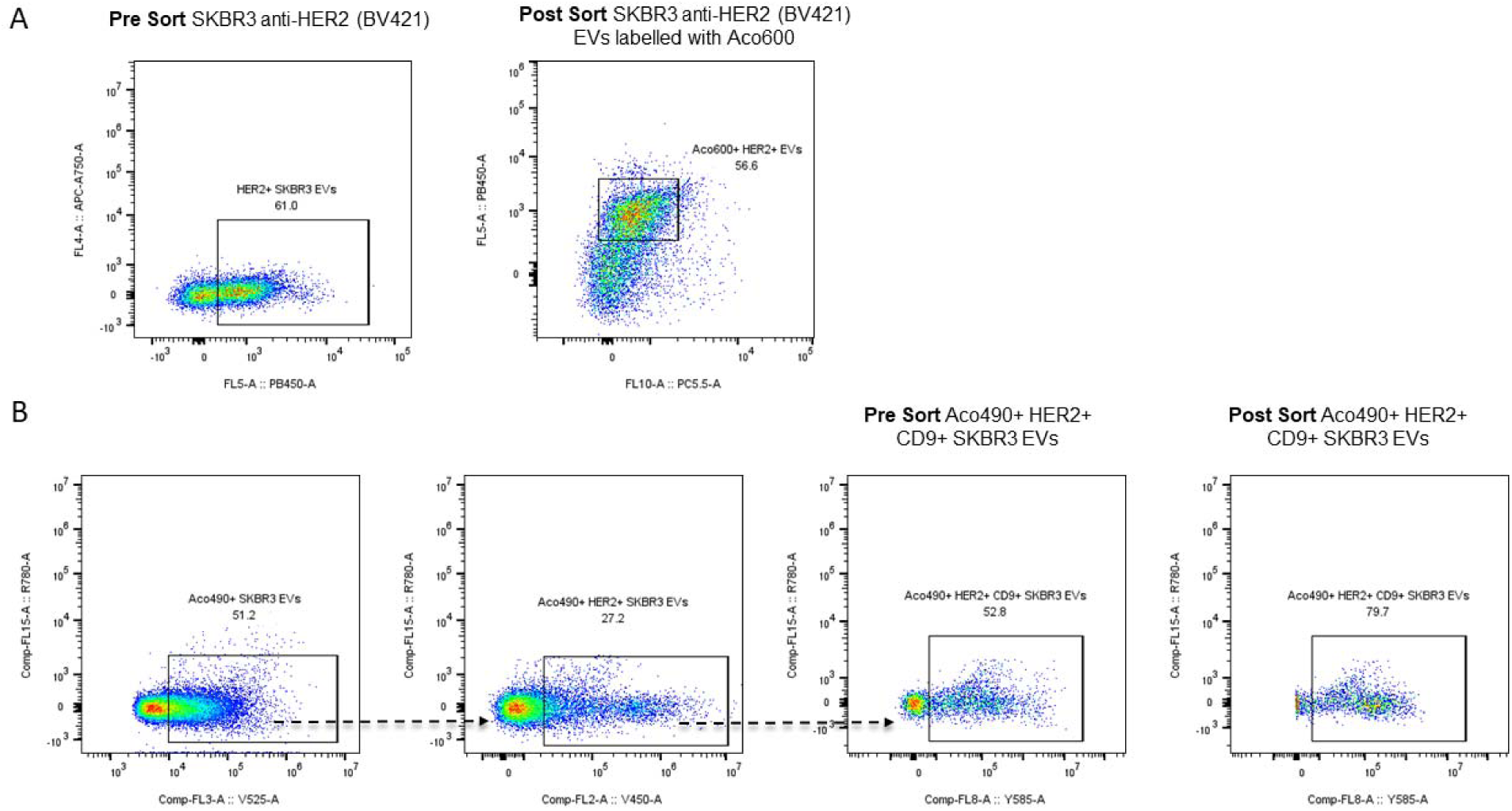
Sorting of HER2+ SKBR3 EVs. A) Single HER2+ (BV421) SKBR3EVs were gated and sorted as “HER2+ SKBR3 EVs”. The collected HER2+ SKBR3 EVs were stained with Aco600 (PC5.5) to confirm their membranous nature. B) Sorting of Aco490+ HER2+ CD9+ SKBR3 EVs, showing the gating strategy progressing from left to right to isolate the “Aco490+ HER2+ CD9+” population. Triple-positive EVs are shown at pre-sort (top) and their post sort re-analysis using a FL trigger (bottom) demonstrating successful enrichment and precise detection of the targeted population.

### 3.3. Sorting nanoparticles and microRNA within biofluids

#### 3.3.1 Sorting of Liposomes, Virus-Like Particles (VLPs) and RBC.EVs from Plasma

To evaluate the developed sorting methodology under more physiologically relevant conditions, we tested standard materials in plasma which contain proteins and biological components that create a more complex and challenging environment for EV detection and sorting. First, standard Liposomes from Acoerela, 490RL and 600RL, were spiked into plasma at a 1:10 ratio and sorted using either VSSC (Figure 9A) or a FL threshold (Figure 9B), with reanalysis performed with both thresholds. Following the same principle, 490RL liposomes together with HIV gag-eGFP VLPs, were spiked into plasma and sorted based on either a double (V525/B525) or VSSC threshold (Figures S13 and S14, Supplementary). The nominal diameter for the 490RL (Fig S13 and S14) and HIV gag-eGFP VLPs (Figure S14) was calculated pre- and post-sorting based on the Core-Shell modelling with FCM_PASS_ confirming that the sorting procedure does not significantly impact EV size statistics.

**Figure 9.**
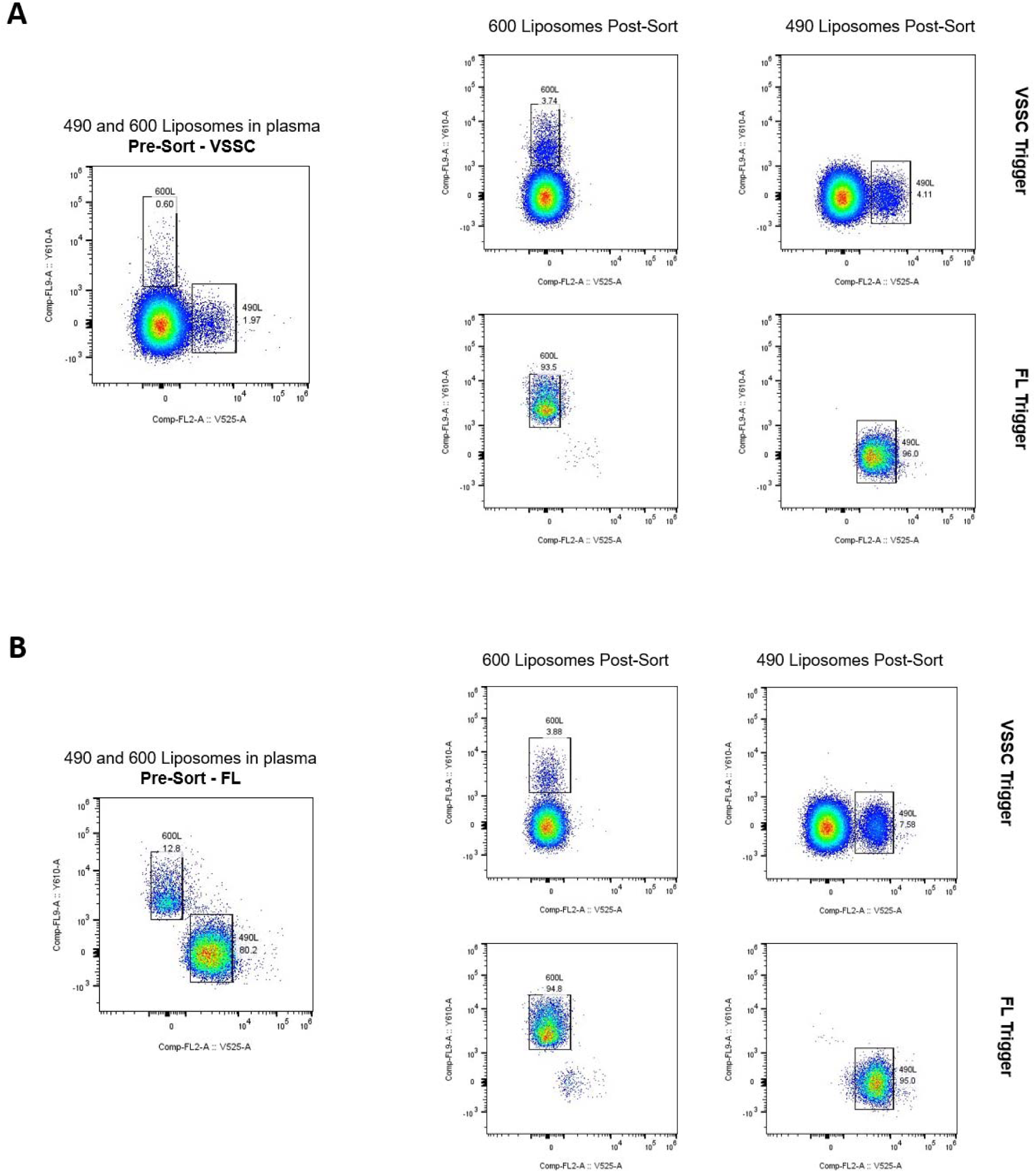
Sorting of 490 and 600 Liposomes using VSSC and FL Thresholds. A) Sorting of 490 and 600 Liposomes in PBS using a VSSC threshold. The controls are shown on top. B) Sorting of 490 and 600 Liposomes in plasma using VSSC threshold and reanalysis using both VSSC and FL thresholds. C) Sorting of 490 and 600 Liposomes in plasma using FL threshold and reanalysis using both VSSC and FL thresholds.

Furthermore, we assessed the feasibility of sorting Aco490/GFP-positive rEVs from plasma. rEVs were spiked into plasma at a 1:10 ratio. Subsequently, the mixture was labeled with Aco490 (5 μL, 10 μM). Aco490-positive rEVs were sorted and confirmed as GFP/490 double-positives by microscopy (Figure 10A). Additionally, Aco490 labelling was performed on Red blood cell-derived EVs (RBC.EVs, ∼1×10^5^) spiked into diluted plasma (1:1000) (Figure 10B), and directly on plasma (Figure 10C). In these sortings, we show that an apparently heterogeneous sample such as plasma can be resolved using Aco490 staining, which helps delineate EV subpopulations even within a complex environment. In figures 10B and 10C, the events initially appear more dispersed in the singlet plot (VSSC-H vs VSSC-Width). However, after sorting based on Aco490 fluorescence, the population is clearly delineated, both for RBC.EVs and plasma, in the singlet plot.

**Figure 10.**
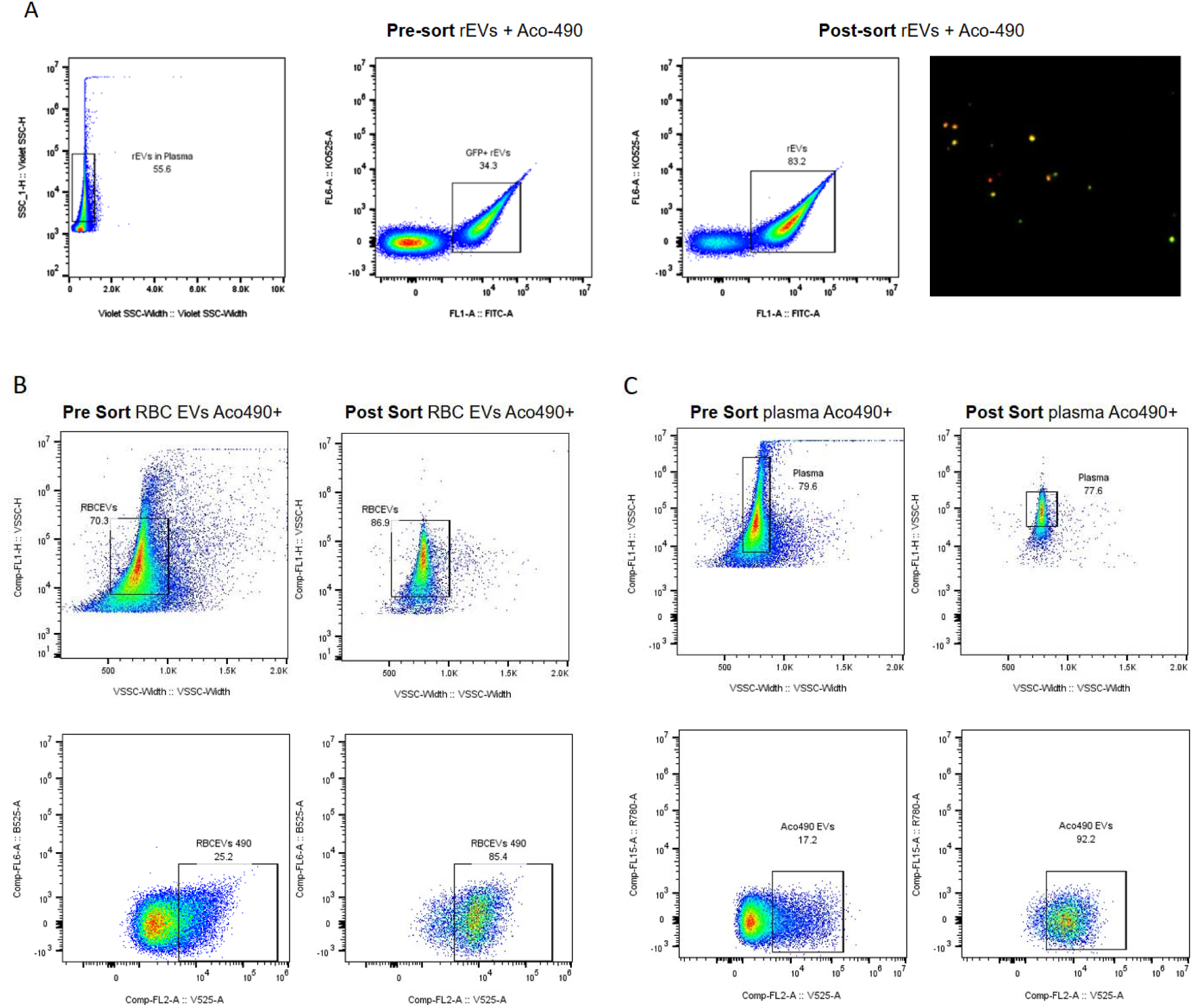
Sorting of rEVs, RBC.EVs and EVs in plasma using Aco490 fluorescence. A) Sorting of rEVs spiked into plasma and subsequently labeled with Aco490. Double-positive EVs (GFP and Aco490) were sorted by nanoFACS and their identity confirmed by microscopy (Darkfield image overlaid with FL). B) Red blood cell-derived EVs were spiked into plasma and then labelled with Aco490 for sorting of Aco490+ EVs. C) Sorting of plasma EVs labeled with Aco490.

#### 3.3.2. Sorting of spiked HeLa CD147+ and/or EGFR+ EVs on plasma

So far, this study demonstrates the successful sorting of reference liposomes, Virus-Like particles, and EVs from plasma based on pre-existing fluorescence (prior to spiking into plasma), or via a lipophilic dye (after spiking into plasma), as well as the sorting of cancer-derived EVs in PBS using antibody labeling of abundant proteins. The clinical translation of EV-based approaches for personalized medicine and oncology requires the detection and sorting of low-abundance, disease-specific EVs directly within complex biofluids. To address this challenge, we used the previously established methodology to investigate the sorting of HeLa cell EVs spiked into plasma, with antibody labeling performed post-spiking. This sequence allowed the plasma protein corona to form on the EVs, which has been suggested to interfere with antibody accessibility and labeling (<u>Försönits et al., 2025</u>), thereby more accurately mimicking the sorting of patient-derived plasma EVs. HeLa cells express Epidermal Growth Factor Receptor (EGFR) and CD147, both of which are associated with cervical cancer progression (<u>Tien et al., 2025</u>, <u>Xin et al., 2016</u>). Following EV isolation, single HeLa EVs were identified as CD147⁺ or EGFR⁺, with CD147 serving as a high-abundance marker (22.5%) and EGFR as a low-abundance marker (0.27%) (Figure S15). Due to the low abundance of EGFR EVs and substantial antibody background (Figure S16C), we refined the population definition: double-positive CD147⁺ EGFR⁺ particles were considered to be EGFR^+^ EVs, whereas CD147^-^EGFR⁺ particles were attributed to antibody background (Figure 11).

**Figure 11.**
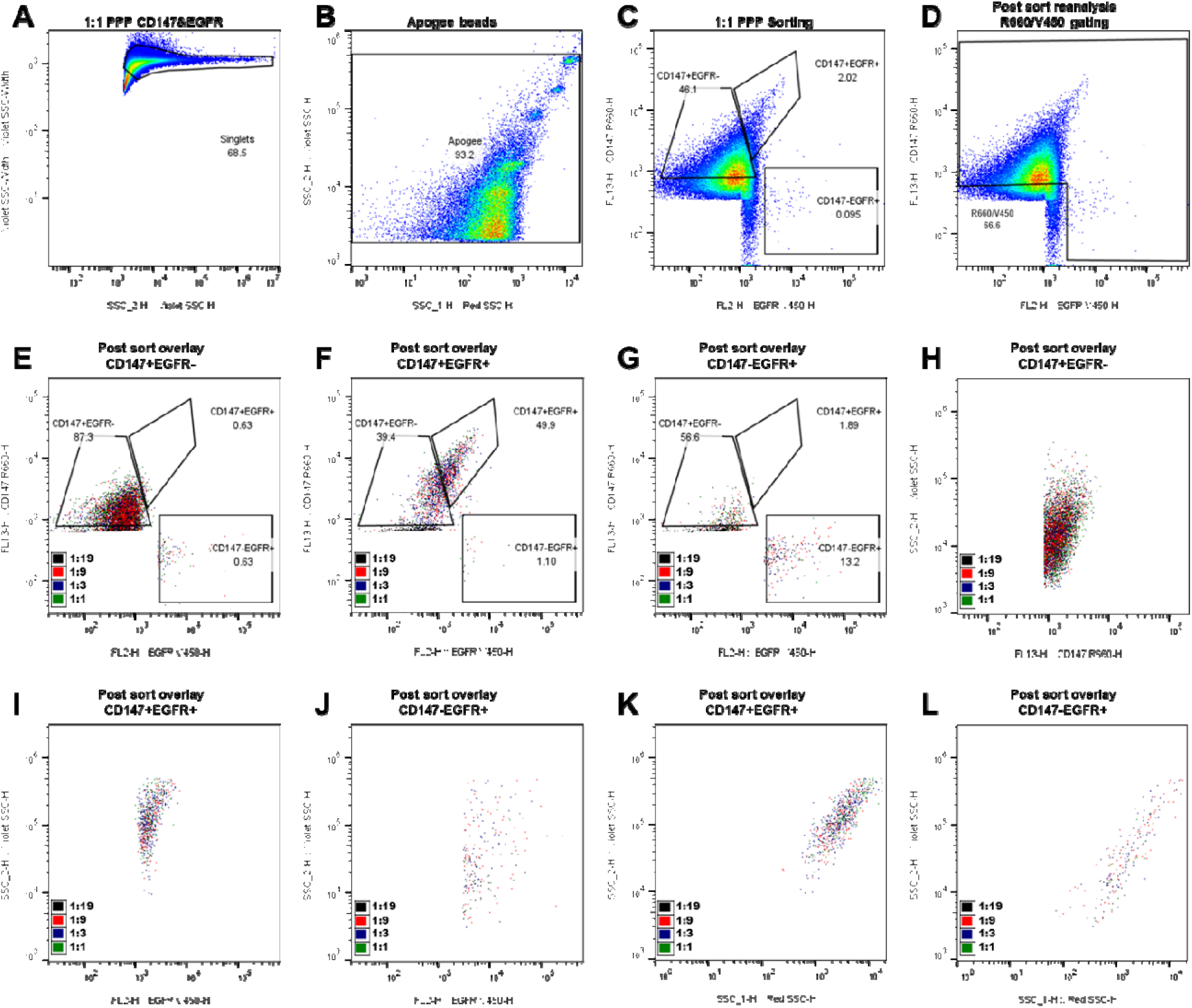
CD147+ and/or EGFR+ HELA EVs sorting from plasma using the CytoFLEX SRT. An EV sorting gating strategy was established by first defining a singlet gate on a VSSC-width/ VSSC-H dot-plot (A) to remove coincident events. An Apogee gate was defined on a VSSC-H/ RSSC-H dot-plot using Apogee beads to eliminate events larger than 500 nm polystyrene bead (B). A R660-H vs V450-H dot plot illustrates the double stained spiked sample under the dual fluorescence threshold and the gating strategy used to sort the stained EVs into CD147+EGFR-, CD147-EGFR+ and CD147+EGFR+ populations (C). For post-sort reanalysis, a R660/V450 gate was applied to remove events near the threshold boundaries or below the specific population gates (D). Overlays of the three sorted populations on R660-H vs V450-H dot plots (E–G) revealed that the CD147+EGFR-EVs were homogeneous but dimmer than the double-positive population (E), while CD147+EGFR+ double-positive EVs (F) correspond to a homogeneous group of EVs with a greater CD147-APC MFI. In contrast, the CD147-EGFR+ events exhibited a highly heterogeneous size distribution (G). Further analysis of the sorted populations on VSSC-H vs R660-H or V450-Hdot plots (H, I & J) showed that CD147+EGFR-EVs (H) are smaller possessing a lower VSSC-H intensity compared to CD147+EGFR+ EVs (I), whereas CD147-EGFR+ events appear as a diffuse, heterogeneous mixture of events with variable VSSC and EGFR-BV450 intensities (J). EGFR+ sorted populations were plotted on VSSC-H vs RSSC-H dot plots (K & L) showing that CD147+EGFR+ EVs (K) and CD147-EGFR+ EVs (L) have unique optical scatter signatures, with CD147-EGFR+ EVs having lower VSSC-H and RSSC-H intensities.

HeLa EVs and plasma particles were quantified as previously described (<u>Brennan et al., 2022</u>) and diluted to working stocks with a concentration of 6 × 10^6^ particles/µL. These stocks were combined to create spiked plasma solutions at ratios of 1:1, 1:3, 1:9, and 1:19. For EV sorting, the following gating strategy was employed: first, a singlet gate was defined on a VSSC-Width vs. VSSC-H dot-plot to remove coincident events (Figure 11A). Events within the singlet gate were plotted on a VSSC vs. RSSC dot-plot via VSSC thresholding, where an "Apogee gate" was used to exclude particles with VSSC intensities greater than those of 500 nm polystyrene beads (Figure 11B). This gate was applied to establish the HeLa EV population. This gate was then applied to samples sorted under a dual fluorescence threshold to include particles that would have been excluded by a standard VSSC threshold; the established gate was verified by backgating of the FL+ population. The remaining events were plotted on an R660-H vs. V450-H dot-plot to identify three distinct EV populations: CD147^+^ EGFR^-^, CD147^-^ EGFR^+^ and CD147^+^ EGFR^+^ (Figure S16E). Gate limits were optimized for purity and yield through iterative sorting and reanalysis using control samples (antibodies in PBS, single- and double-stained samples) as references (Figure S16B-E). Due to the low percentage of events captured under a VSSC threshold, sorting was performed using a dual EGFR-BV450/CD147-APC fluorescence threshold. This approach significantly improved detection; for example, it increased the percentage of CD147^+^ EGFR^-^ events from 14% (Figure S16E) to 46.1% (Figure 11C).

Post-sort analysis of the three sorted EV populations (Figure S16F–H) revealed that the double-positive population (CD147+ EGFR+) corresponds to a homogeneous group of larger EVs. In contrast, CD147+ EGFR-EVs were homogeneous but smaller than the double-positive population (Figure S16F), while CD147-EGFR+ EVs displayed a relative heterogeneous size distribution (Figure S16G). Following the successful sorting of CD147+ and/or EGFR+ HeLa EVs from plasma at a 1:1 ratio, the process was repeated at decreasing EV:plasma ratios (1:3, 1:9, and 1:19) to assess feasibility at concentrations more reflective of clinical samples. An "R660/V450" gate was applied during reanalysis to exclude events near the dual-threshold border and those below the specific population gates (Figure 11D); this simplified the detection of low-abundance events in post-sort overlays. Across the 1:1 to 1:19 dilutions, the fluorescence profiles for the three sorted populations remained consistent (Figure 11E–L). Post sort analysis revealed that the majority of CD147+ EGFR-sorted events were recovered within the original gate (Figure 11E). However, a decrease in EGFR-BV450 and CD147-APC staining intensity of CD147+ EGFR+ sorted EVs was observed, with 39.4% of CD147+ EGFR+ events shifting into the CD147+ EGFR- gate (Figure 11F). Given that the population remained homogeneous, we propose this signal shift results from antibody dissociation from the EV surface following the sorting process. While the fluorescence profiles of CD147- EGFR+ sorted events were consistent across dilutions (Figure 11G), only 13.2% of events remained within the target gate. This is significantly lower than the recovery rates for CD147+ EGFR- EVs (87.3%) or CD147+ EVs from the double-positive sort (89.3%). We propose that these CD147- EGFR+ particles are primarily composed of EGFR-BV450 antibody background. Furthermore, the droplet-mediated dilution inherent to the sorting process likely causes these antibody aggregates to dissociate, preventing them from falling within the original gate during reanalysis.

To further characterize these samples, the VSSC (indicative of size) was compared with the marker fluorescence profiles across the four dilutions, revealing that all samples maintained a consistent size and fluorescence distribution regardless of the spike ratio. Specifically, CD147+ EGFR- EVs (Figure 11H) exhibited lower VSSC intensity compared to the CD147+ EGFR+ EV population (Figure 11I). The CD147+ EGFR+ EVs (Figure 11I) presented as a defined, homogeneous population with high VSSC, while the CD147- EGFR+ events (Figure 11J) appeared as a diffuse, heterogeneous mixture spanning a wide range of VSSC and EGFR-BV450 intensities, which supports our hypothesis that these events largely represent antibody background rather than distinct EV populations. Further analysis of the EGFR+ sorted populations on VSSC-H vs RSSC-H dot plots showed that CD147+EGFR+ EVs (Figure 11K) and CD147-EGFR+ EVs (Figure 11L) have different optical scatter signatures, which suggests these two particle populations have different membrane and/or cargo composition. The overlays of the sorted populations demonstrate that even when target EVs are rare, they can be reliably detected and isolated; however, the clinical utility of this method is inherently tied to event frequency, as lower EV concentrations will result in significantly extended sort times. This practical constraint is highlighted by the sort rate of the CD147+ EGFR- population decreasing from 84 events/s in the 1:1 spiked plasma to 3 events/s in the 1:19 dilution.

#### 3.3.3. Sorting of miRNA using beads

Molecular beacons are single-stranded oligonucleotide hybridization probes that adopt stem-and-loop (hairpin) conformation. The loop contains a probe sequence complementary to a specific target nucleic acid, while the stem is formed through intramolecular base pairing between complementary arm sequences flanking the probe region. A FAM fluorophore is covalently attached to the end of one arm and a ZEN quencher to the other. In their native folded state, molecular beacons do not fluoresce because the close proximity of the fluorophore and quencher suppresses emission. When the beacon hybridizes to a complementary target sequence, however, it undergoes a conformational rearrangement that separates the fluorophore from the quencher, resulting in strong fluorescence upon excitation.

A wide range of extracellular and intracellular RNAs are normally complexed with RNA-binding proteins, which in principle could hinder detection strategies that rely on molecular beacons. Nevertheless, oligonucleotide probes exhibit substantially stronger affinity for their complementary sequences than RNA-binding proteins do for their RNA partners, often by more than an order of magnitude. As a result, molecular beacons can still reliably hybridize to their target mRNAs inside living cells, effectively displacing the proteins that were previously bound to the RNA (<u>Oliveira GP Jr, 2020, Vargas DY, 2005, Bratu DP, 2003</u>, <u>Chen M, 2017</u>).

miR-451 was initially identified in human tissues in 2005 and has since been recognized as a key regulator of diverse physiological and pathological processes. This microRNA plays an important role in cellular differentiation and development across multiple systems and is frequently dysregulated in cancer. Altered miR-451a expression has been associated with tumor initiation and progression, influencing critical oncogenic pathways related to cell proliferation, apoptosis, angiogenesis, epithelial–mesenchymal transition, drug resistance, and metastasis. In many malignancies, miR-451a functions predominantly as a tumor suppressor by modulating gene expression through the targeting of multiple downstream mRNAs, underscoring its relevance as both a biomarker and a potential therapeutic target (<u>Xu et al., 2019</u>).

However, miRNA expression levels typically differ among donors and health status and are often difficult to detect even with qPCR due to pre-analytical variability resulting from sample preparation. Challenges in miRNA detection also apply to nanoparticle Flow Cytometry, specifically due to limitations with size-based detection and increased background noise from other plasma constituents or from the cytometer itself. In fact, miRNAs have been found to have a contour length of only approximately 6-7 nm, which is significantly smaller than the widely-accepted detectable nanoparticle size – 100 nm. To combat such limitations, and to increase detectable signal and specificity of detection, custom-designed molecular beacon (MBs) were employed with a miRNA-specific reverse complementary sequence and an additional biotinylated region to allow for conjugation to streptavidin (SA) beads. With this technology, fluorescence signal is only present with binding of MB with its specific target and is concentrated within a specific bead population on a side-scatter plot. In this study, biotinylated MB – specific to miR-451a – was validated on the Beckman Coulter CytoFLEX SRT cell sorter (Figure 12) for bead-based sorting.

**Figure 12.**
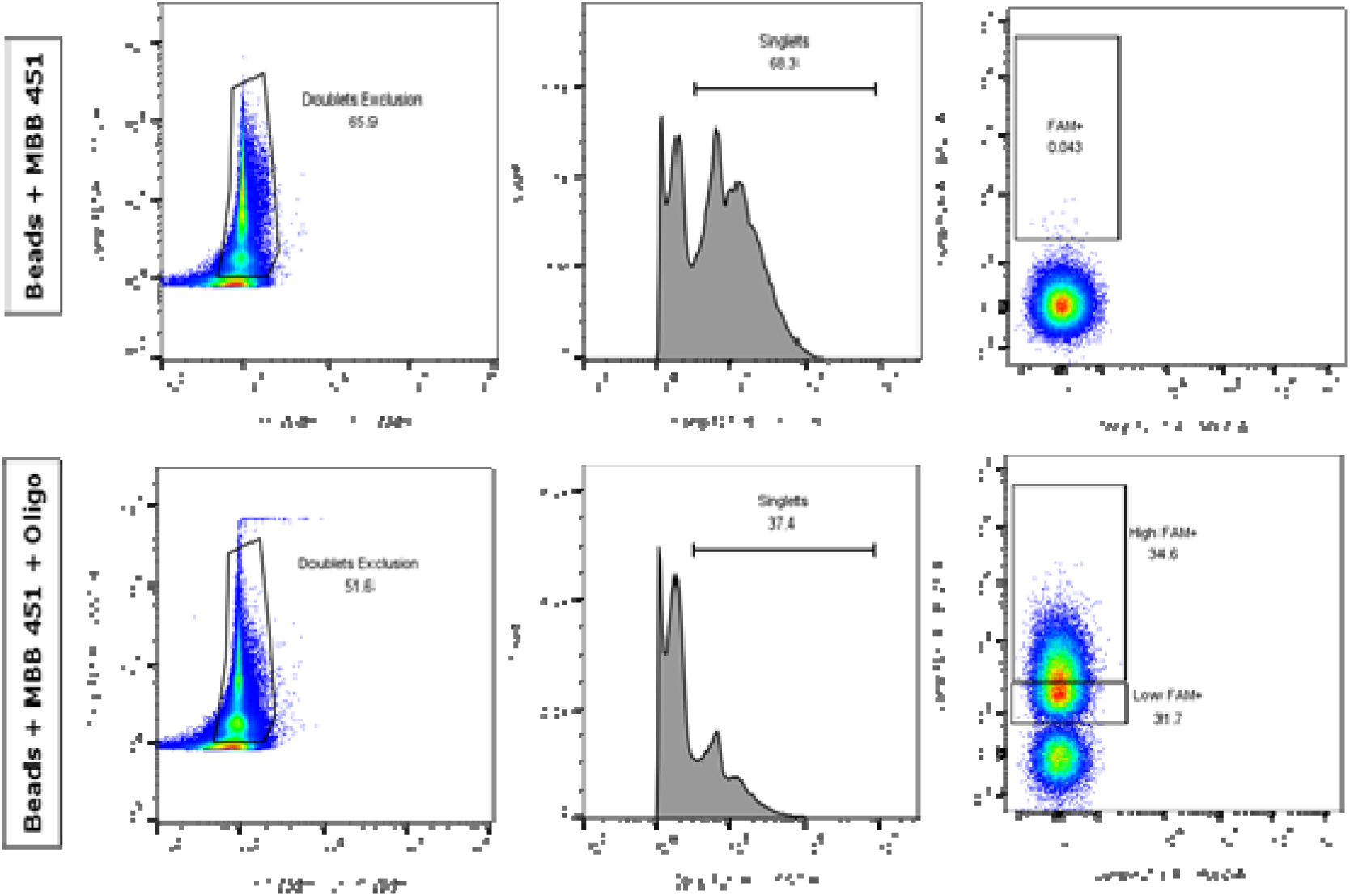
Validation of MB-451a specificity using streptavidin beads by nano-Flow Cytometry. Streptavidin bead–based nano-flow cytometry assay to validate the specificity of MB-451a detection of miR-451a. (Top row) Beads incubated with MB-451a alone show minimal FAM signal, confirming low background fluorescence. (Bottom row) Addition of the complementary miR-451a oligonucleotide results in a marked increase in FAM fluorescence, with clear separation between Low and High FAM⁺ populations. Doublet exclusion, singlet gating, and fluorescence intensity analysis were performed using a CytoFLEX SRT. The analysis strategy follows the doublet exclusion gate, and a selection of singlets events, not associated with “noise”. On singlets events, FAM/B525+ events were selected (FAM+) based on the presence of the target. These results confirm target-dependent activation of MB-451a.

To validate the findings obtained by nano-Flow Cytometry, the same experiment was further analyzed using dark-field microscopy (Figure 13). MB Beads lacking the target were first isolated and examined on a microscope slide. To confirm target specificity, the target was gradually added directly to the slide, resulting in a progressive emergence of fluorescence from the MBBs upon binding to the specific target.

**Figure 13.**
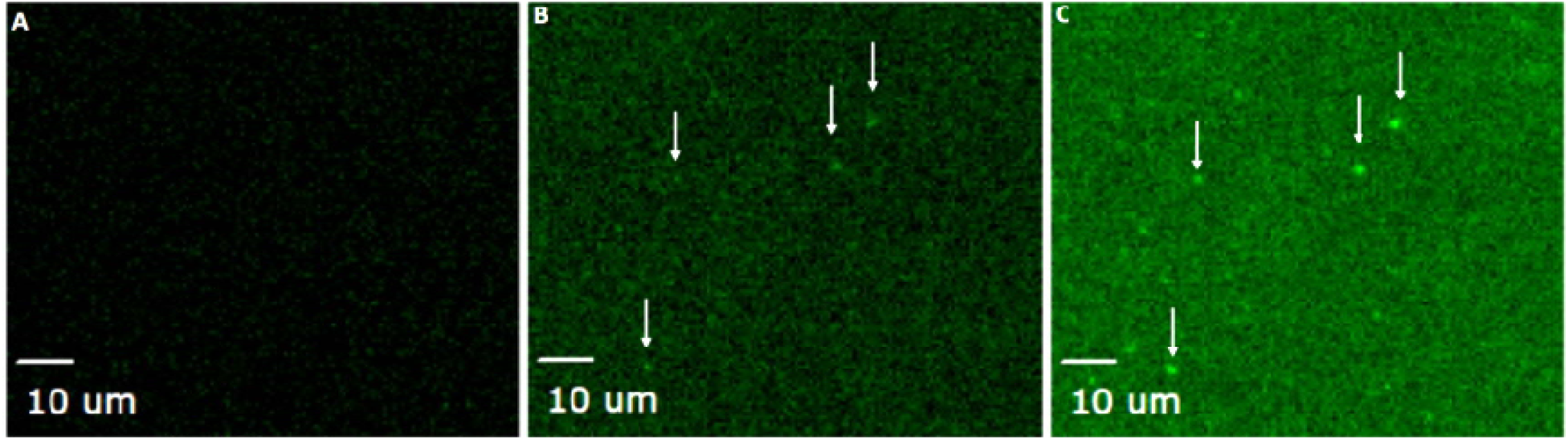
Validation of MB-451a specificity by dark-field and fluorescence microscopy. Dark-field and fluorescence microscopy analysis of MBBeads in the absence and presence of the miR-451a target. (A) MBBeads without target show minimal background fluorescence. (B) Incremental addition of miR-451a to the slide results in the appearance of discrete fluorescent signals (arrows), indicating target-dependent activation of the molecular beacon. (C) Increased number and intensity of fluorescent MBBeads following further target addition, confirming the specificity of MB-451a for miR-451a. Scale bars: 10 µm.

As previously mentioned, miR-451a is widely recognized as a biomarker in multiple pathological conditions, including autoimmune diseases and cancer, where its differential expression has diagnostic and prognostic relevance. Accurate detection and quantification of miR-451a are thus critical for both biomarker validation and translational applications. Sorting miR-451a–positive RBC.EVs enables validation of the detection strategy and facilitates downstream molecular and functional analyses. RBC.EVs have been shown to carry miRNA-451a; therefore, a complementary molecular beacon (MB-451a) was designed to enable the specific detection of miR-451a within these vesicles (Figure 14). Prior to molecular beacon hybridization, RBC.EVs were lysed using MagIC Lysis Buffer to release intravesicular miR-451a, after which the lysates were incubated with the MBBeads as described above. The RBC.EV MB451-AF488 sample was analyzed under FITC channel fluorescent triggering. Gating of populations was performed according to established methodology and High expressors were sorted.

**Figure 14.**
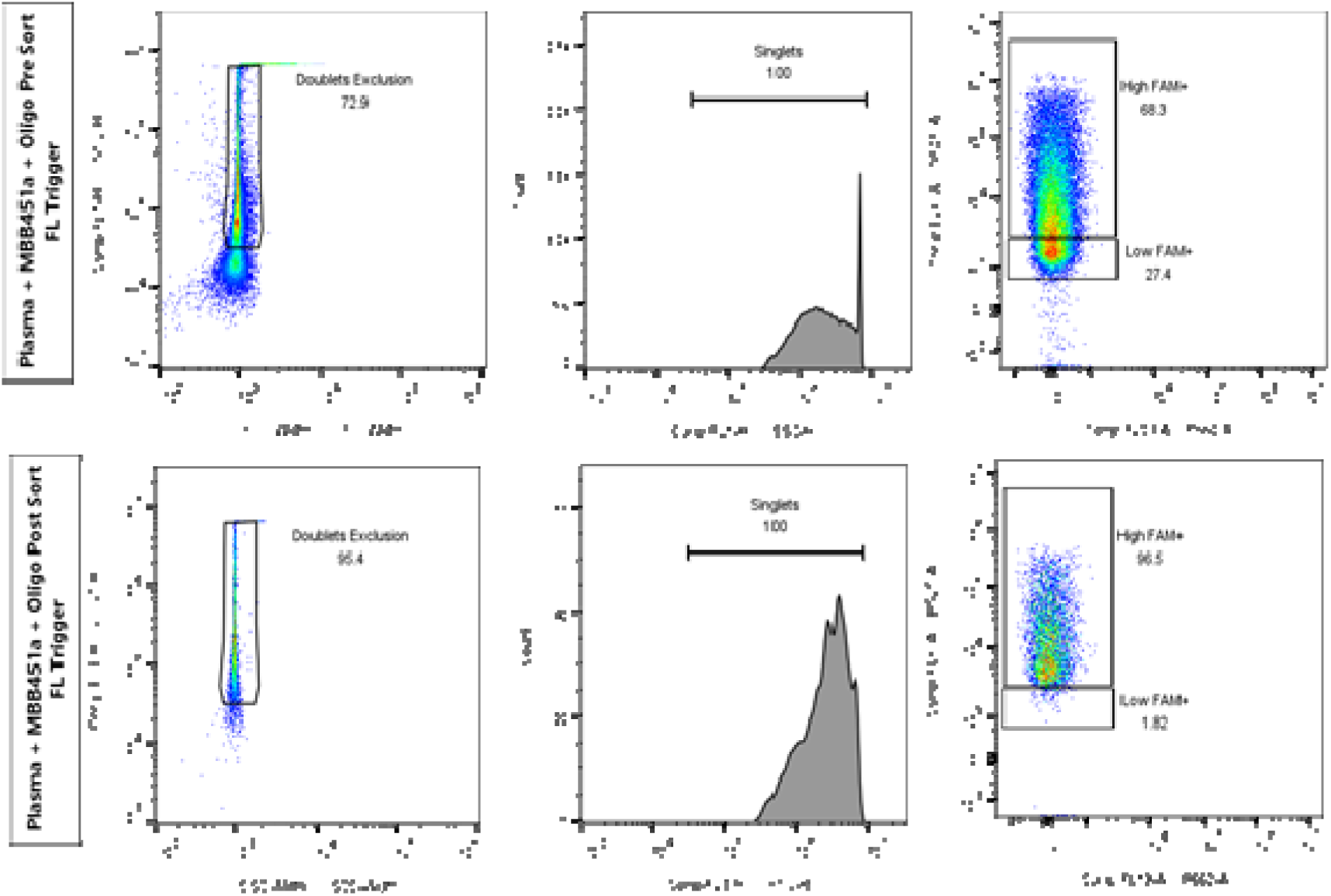
nanoFACS of miR-451a in plasma using MBB-451a. nanoFACS analysis of plasma samples labeled with MB-451a and sorted on a CytoFLEX SRT. (Top row) Pre-sort analysis showing doublet exclusion, singlet discrimination, and separation of Low and High FAM⁺ populations following incubation with MB-451a and trigger oligonucleotide. (Bottom row) Post-sort analysis demonstrating enrichment of the High FAM⁺ population after sorting. Gating strategy includes exclusion of aggregates, selection of singlets, and identification of FAM⁺ events corresponding to miR-451a–positive vesicles.

## 4. Discussion

In recent years, nano-Flow Cytometry has become an essential technology for the analysis of EV populations. While early studies demonstrated that nanoparticles could be sorted using conventional flow cytometers (<u>Danielson et al., 2016</u>; <u>Groot et al., 2016</u>; <u>Higginbotham et al., 2016)</u>, they also underscored the technical challenges involved—especially the extensive hardware modifications required to adapt traditional cell sorters for nanoscale resolution (<u>Morales-Kastresana et al., 2019</u>). Later developments simplified aspects of the workflow (<u>Morales-Kastresana et al., 2020</u>) and even enabled downstream analyses such as EV miRNA profiling (<u>Oliveira et al., 2020</u>), yet standardization and accessibility remain as significant limitations (<u>Welsh et al., 2023</u>). In response to these challenges, the CytoFLEX SRT was proposed as a commercially available platform capable of overcoming prior technical hurdles associated with nanoscale particle sorting. With hardware inherently compatible with its analytical counterpart, the CytoFLEX S, this sorter offers a streamlined interface for nanoparticle detection and isolation. Accordingly, the present study aimed to develop a robust nanoFACS workflow using the CytoFLEX SRT, to evaluate its performance in detecting, characterizing, and sorting nanoparticles, and to assess its effectiveness on increasingly complex biological samples. To this end, the study was structured in three phases: phases 1 and 2 focused on establishing the feasibility of using the CytoFLEX SRT for the detection and isolation of nanosized standard particles based on size and fluorescence, while evaluating the instrument’s sensitivity and detection limits. In the final phase, the optimized methodology was applied to the characterization and sorting of EVs derived from complex biological samples. Overall, the results support the main hypothesis and demonstrate the potential of nanoFACS for robust nanoparticle and EV sorting, particularly through the use of protein-specific fluorescent tags.

Previous work has established that the CytoFLEX S is capable of detecting and characterizing nanosized particles based on size and/or fluorescence (<u>Brittain et al., 2019</u>; <u>Burnie et al., 2025</u>). Similar to the CytoFLEX S, the CytoFLEX SRT can detect particles based on size or fluorescence but also incorporates a fully integrated sorting mechanism via the CytExpert software. Sorting is achieved through the application of electrical charges to droplets containing target particles. As these droplets pass through the sorting chamber, they are deflected by high-voltage plates into designated collection tubes, based on user-defined gating regions. The system’s minimal hardware modifications and automated interface streamline the process, making nanoscale sorting more accessible and reproducible across experiments. This streamlined workflow not only facilitates the isolation of EV subpopulations, but also preserves their integrity for downstream analyses. For example, sorted EVs can be subjected to molecular assays such as Western Blotting or Gel Electrophoresis to probe their protein and nucleic acid content (<u>Arraud et al., 2017</u>; <u>Marchisio et al., 2023</u>; <u>Choi et al., 2020</u>). These approaches enable validation of disease-associated biomarkers and functional cargo within specific EV subsets thereby strengthening biological interpretation and translational relevance.

Alongside the sorting capabilities of the CytoFLEX SRT, FCM_PASS_ software was implemented to standardize both light scatter and fluorescence measurements, ensuring accurate particle discrimination and reliable data comparability across instruments. NIST-traceable beads were used for scatter calibration, and MESF standards for fluorescence, minimizing instrument-dependent variability and supporting consistent analysis of EV subpopulations. This calibration framework ensures that observed differences in size and fluorescence reflect biological heterogeneity rather than technical artifacts. Furthermore, by incorporating calibration metadata in accordance with community reporting standards such as MIFlowCyt-EV, FCM_PASS_ enhances transparency, reproducibility, and cross-laboratory comparability of nanoscale Flow Cytometry datasets (<u>Welsh et. al, 2020</u>). In line with these principles, all calibrated data and associated metadata have been made publicly available through the NanoFlow Repository, enabling independent validation and reuse of the datasets (<u>Arce et al., 2023</u>).

Based on manufacturer product descriptions and post-FCM_PASS_ analysis of NIST bead sets, the CytoFLEX SRT has limitations in terms of ability to detect particles of sizes below 100 nm. However, this limitation is resolved with the use of fluorescence-based sorting. While VSSC is an effective method for separation, it results in a skewed % positive upon post sort analysis as described in the sorting of NIST beads previously (Figure 2). This is due to instrument noise, more commonly referred to as "dark" or "shot" noise (<u>Jia et al., 2024</u>). Dark noise is inherent to nanoFACS as thresholds are decreased to enhance the limit of detection for nanoparticles. As the signal to noise ratio becomes dependent on concentration of the sorted sample, nanoparticles are diluted by the droplets formed and their concentration is decreased. The noise component, which is a consistent value, has a higher event rate than the sorted nanoparticles. This leads to the misrepresentation of percent purity. In order to have a better representation of purity using VSSC, the sorted nanoparticles should be concentrated. In this study, all events are represented to highlight the separation and purity of populations based on multiparametric flow cytometry and to show the effect of dark noise on S/N. An example of how fluorescence-based detection can improve population representation is illustrated in Figure 3, where Fluoresbrite YG microspheres are used to demonstrate accurate separation without the need for post-sort concentration.

To evaluate the performance of fluorescence-based sorting for small biological particles, we utilized virus-like particles (VLPs), mouse leukemia virus (MLVs) and recombinant EVs (rEVs) as model systems. VLPs, tagged with either eGFP or tdTomato, served as a well-characterized model for fluorescence nanosorting experiments. Additionally, MLVs (ViroFlow) were included due to their defined structural composition and predictable behaviour in flow cytometry.

Similarly, GFP-labeled rEVs (Sigma-Aldrich), which are commercially available fluorescent standards, were employed for their known properties and consistency. rEVs share important biochemical and physical characteristics with native EVs, such as size (50-180 nm), morphology, and protein content (including Tetraspanins CD9, CD63, and CD81). These features make them ideal for quality control, instrument calibration, and method development in EV research, ensuring reproducibility and accuracy.

Figure 4 summarizes two distinct gating approaches for sorting based on 525 nm fluorescence intensity levels. In the case of MLV-derived VLPs (Figure 4A), sorting was performed by dividing the population into two groups—low and high sfGFP intensity—without predefined MFI differences, allowing us to assess the resolution of fluorescence-based separation. This approach serves as a model for cases where particles exhibit a clear fluorescence signal. Conversely, for HIV gag-eGFP VLPs (Figure 4B), sorting was refined into three subpopulations (“GFP Low”, “GFP Mid” and “GFP High”), better reflecting scenarios where fluorescence intensity varies more subtly, akin to EVs with different epitope densities. These strategies highlight the adaptability of fluorescence-based sorting for distinguishing small particle subpopulations beyond size-based discrimination. Expanding on this approach, we introduced Aco490 labeling to further refine the separation of EV subpopulations (Figure 5). While fluorescence-based sorting previously allowed differentiation based on a single fluorescent signal, the addition of Aco490 provided an additional selection parameter. This enabled a more precise distinction between EVs, not only by their intrinsic fluorescence but also by their membrane composition. Even with the commercially available rEVs, we observed two distinct populations: one double-positive for both GFP and Aco490, and another Aco490-positive but GFP-negative. In this context, the rEVs—produced via transfection of HEK293T cells (Sigma-Aldrich)—correspond to the GFP/Aco490 double-positive EVs, while the Aco490-only population could represent EVs naturally released by the same cells. This revealed the complexity within EV populations and demonstrated that combining intrinsic fluorescence and membrane markers allows for the identification and isolation of previously indistinguishable subpopulations, showcasing the enhanced capabilities of Single Vesicle nanoFACS. These findings are akin to nanoparticle methodologies where multiple fluorescent tags are used to identify subpopulations associated with a specific protein.

The hardware and software improvements implemented in the CytoFLEX platform enable true dual thresholding using both VSSC and a fluorescence channel of interest, providing a substantial advantage for nanoparticle characterization. This capability enables the simultaneous discrimination of particles based on size and epitope density without compromising S/N. As a result, distinct subpopulations can be resolved more accurately, reducing the contribution of noise-related false-negative events. Importantly, in this study the interpretation and comparison of our sorting results rely on FCM_PASS_-based calibration, which converts instrument-specific detector outputs into standardized units. Furthermore, because the CytoFLEX S and SRT share identical optical architectures, this calibration framework allows analytical and sorting measurements to be directly compared, ensuring that values such as Equivocal GFP Molecules and Core–Shell–derived diameter estimates reflect particle-intrinsic properties rather than instrument-dependent variability. This standardization is essential for nanoscale work, where small changes in gain, thresholding, or fluidics can substantially shift particle statistics. Using this framework, we compared sorting outcomes obtained using VSSC-versus fluorescence-based triggering for previously analyzed rEVs (Figure 6). Both approaches yielded highly consistent results, with VSSC-based sorting resulting in a median FITC intensity of 614 Equivocal GFP Molecules and a nominal diameter of 153 nm, while fluorescence-based triggering yielded a median FITC intensity of 636 Equivocal GFP Molecules and a nominal diameter of 142 nm (RI 1.40–1.48, Core–Shell modeling). This analysis was subsequently extended to another well-characterized biological reference material, HIV gag-eGFP/tdTomato–tagged VLPs (Figure 7). FCMPASS-calibrated analysis of the sorted populations demonstrates that both VSSC- and fluorescence-based triggering yield highly comparable fluorescence intensities, with MESF values of 556 and 563, respectively, indicating preserved epitope density. In contrast, a modest shift in nominal diameter estimates (153 nm for VSSC versus 136 nm for fluorescence-based sorting) reflects the improved sensitivity of fluorescence triggering toward smaller particles rather than a change in particle composition, as refractive index does not influence MESF values. Consistent with the rEV sorting results (Figure 6), these data indicate that rEV and HIV gag-eGFP/tdTomato tagged VLPs statistics remain stable before and after sorting regardless of the threshold strategy employed, highlighting the combined impact of the CytoFLEX platform capabilities and FCM_PASS_ standardization. Furthermore, this framework enables direct cross-instrument comparisons, as illustrated by the FCM_PASS_-calibrated sorting of HIV gag-eGFP/tdTomato–tagged VLPs conducted independently at UCD and BIDMC (Figure S11, Supplemental). Together, these data demonstrate that fluorescence-based thresholding, when properly calibrated, faithfully reproduces VSSC-derived measurements and provides a robust, standardized basis for evaluating nanoparticle sorting performance at the nanoscale.

Based on these findings, the techniques and parameters established in the sorting of small biological standard samples were successfully applied to SKBR3 cell EVs. SKBR3 is a well-established HER2-overexpressing Breast Cancer cell line widely used as a model to study the HER2-enriched molecular subtype of Breast Cancer. This subtype is characterized by aggressive tumor behavior and poor prognosis in the absence of targeted treatment. Despite this, HER2-targeted therapies such as Herceptin (Trastuzumab) have significantly improved patient outcomes, highlighting the clinical relevance of HER2 as a therapeutic and biomarker target. By fluorescently labeling the HER2 surface protein, nanoFACS enabled the isolation of HER2-positive EV subpopulations, providing a strategy to further classify EV subsets based on their surface markers and cargo. The successful sorting of single HER2-positive EVs, followed by post-sort confirmation of their membranous nature, supports the specificity of the approach and minimizes the contribution of antibody aggregates or non-vesicular particles (Figure 8A). In addition, the detection and sorting of EVs based on the simultaneous presence of HER2, membrane labeling, and CD9 EV marker demonstrates the capability of nanoFACS to resolve EV heterogeneity at the single-vesicle level (Figure 8B). Together, these results support nanoFACS as a robust platform for the selective isolation and characterization of biologically relevant EV subpopulations derived from breast cancer cells. While this study focused on HER2-positive EVs, other EV-associated markers such as CD44 and tetraspanins including CD81 have been reported to play key roles in breast cancer recurrence and metastasis (<u>Ramos et al., 2021</u>; <u>Ramos et al., 2022)</u>. Therefore, the ability to selectively isolate EV subpopulations based on protein markers may enable more comprehensive analyses of EV cargo and support the development of EV-based strategies for disease monitoring and mechanistic studies.

Next we evaluated the sorting methodology under physiologically relevant conditions, using plasma as a complex matrix to challenge EV detection and sorting. Direct-from-plasma sorting is a primary goal in liquid biopsy research, as it shields EVs from the mechanical stress and buffer-induced artifacts inherent in traditional isolation methods such as ultracentrifugation and size-exclusion chromatography. We demonstrated that 490RL and 600RL liposomes (Acoerela), as well as HIV gag-eGFP VLPs, could be successfully sorted from both PBS and plasma without significantly altering their size statistics. Subsequently, rEVs and RBC-derived EVs were spiked into diluted plasma followed by in situ labelling with the fluorogenic dye Aco490. This demonstrated the dye’s capacity to discriminate distinct EV populations within a complex biological environment; Aco490-positive rEVs were isolated and their identity was confirmed via microscopy. Finally, we assessed the feasibility of sorting HeLa-derived EVs spiked into plasma. Notably, antibody labeling was performed after spiking to allow for the formation of a plasma protein corona. This approach more accurately mimics patient samples, as the corona has been shown to potentially interfere with antibody accessibility and labeling efficiency (<u>Försönits et al., 2025)</u>. CD147 and EGFR were selected as target markers based on their expression profiles in HeLa cell EVs, where CD147 served as a high-frequency marker (22.5%) and EGFR as a low-frequency marker (0.27%). We observed that this dual-staining approach facilitated the detection of weak EGFR+ EVs, which would otherwise be obscured by background noise in a single-stain control, while effectively discriminating EV events from non-specific antibody aggregates. Post sort analysis of the CD147+ EGFR-, CD147- EGFR+ and CD147+ EGFR+ sorted populations revealed three distinct populations with different size characteristics and marker expression. These characteristics remained stable across various spike ratios (1:1 to 1:19), validating the robustness of in-plasma EV sorting for liquid biopsy applications. However, the increased acquisition time required for samples with low marker frequency remains a significant bottleneck for the clinical translation of assays targeting extremely rare EV subpopulations.

The application of molecular beacons (MBs) combined with nano-Flow Cytometry represents a significant methodological advance for the detection and isolation of microRNA-containing Extracellular Vesicles. One notable application of this approach was described by <u>Oliveira et al. (2020)</u>, who used molecular beacons (MBs) in combination with nanoscale Flow Cytometry to investigate the miRNA content of red blood cell-derived EVs (RBC.EVs). Each MB is customized for specific applications becoming an advantageous tool, capable of distinguishing DNA sequences with a single nucleotide difference and enabling studies in areas evolving protein-DNA and RNA-DNA interactions, living systems and biosensors (<u>Tan et al., 2004</u>).

In this study, MB-based detection enabled highly specific identification of miR-451a at the single-particle level, overcoming several limitations traditionally associated with miRNA analysis in complex biological matrices. Most notably, a critical challenge in miRNA detection is the fact that extracellular and intracellular RNAs are frequently complexed with RNA-binding proteins, which can hinder probe accessibility. Nevertheless, MBs exhibit substantially higher binding affinities to their complementary sequences than RNA-binding proteins, enabling effective displacement of protein–RNA interactions and reliable hybridization even within biologically complex environments. The hairpin-loop structure of MBs ensures minimal background fluorescence in the absence of target nucleic acids, while target hybridization induces a conformational change that generates a robust and quantifiable fluorescent signal (<u>Tyagi & Kramer, 1996</u>; <u>Giesendorf et al., 1998</u>). This target-dependent activation is particularly advantageous in nanoscale applications, where signal-to-noise ratios are often compromised by instrument noise and low analyte abundance.

These properties have been previously demonstrated in live-cell and EV-based systems and is further supported here by the successful detection of miR-451a within red blood cell–derived extracellular vesicles (RBC.EVs). The validation of MB-451a specificity using both nano-flow cytometry and dark-field fluorescence microscopy underscores the robustness of this approach and confirms that fluorescence activation is strictly dependent on target presence (Figures 12-14). Moreover, the lysing of RBC.EVs, within a plasma substrate, and subsequent detection via MB-451a, exemplifies the ability of nanoFACS to isolate circulating miRNAs from biofluids. These findings lead to the possibility of multiplexing miRNA detection using multiple MB sequences with varying fluorophores.

In totality, the results have shown the capability of nanoFACS to detect, characterize, and isolate nanoparticles. Results have demonstrated that previously isolated nanoparticles, and those from biofluids can be recovered with high efficiency and minimal manipulation. This methodology allows for more standardized, less time consuming, and more robust isolation from contaminants of nanoparticles, especially EVs and their cargo.

## 1. Conclusions

Collectively, these results demonstrate that the CytoFLEX SRT is a robust and versatile platform for the detection and isolation of nanoscale particles from complex biofluids. By integrating fluorescence-based detection, multiparametric gating, and reproducible sorting workflows, this nanoFACS methodology addresses some key limitations of previous approaches and positions the CytoFLEX SRT as a promising candidate for standardized EV analysis in both research and translational settings, including future applications in patient sample profiling. While limitations still exist in the form of epitope densities, labeling efficiencies, limit of detection, and sample dilution; nanoFACS via the CytoFLEX SRT has been shown to be a standardizable and reproducible method.

## 2. Acknowledgements

We gratefully acknowledge Acoerela for kindly providing reagents and expertise. We would like to thank Dr. Jennifer C Jones (NIH/NCI) for her mentorship and expertise in nanoFACS. In addition, we would like to thank Anietie Uko and Mamadou Diallo (BC) for their field service expertise. To Kenneth Gray (BC) for providing reagents and expertise with the CytoFLEX products. The work was funded by the Horizon Europe Twinflag consortium (HORIZON-WIDERA-2021-ACCESS-03-01/101079489) and also supported by SFI Infrastructure Programme Award (21/RI/9718) and Research Ireland RD&I Fellowship (23/IRDIFB/12112).

## Author Contributions

Conceptualization – MMG, JT, AB; Methodology – IC, JT, VC, BP, KB, AB; Investigation – IC, JT, VC, BP, IG, KB, AB, FD, CR; Resources – MMG, JT, AB; Supervision – MMG, JT; Writing (original draft) – IC, JT; Writing (review & editing) – All authors

## References

Arce, J. E., Welsh, J. A., Cook, S., Tigges, J., Ghiran, I., Jones, J. C., Jackson, A., Roth, M., & Milosavljevic, A. (2023). The NanoFlow Repository. Bioinformatics (Oxford, England), 39(6), btad368. 10.1093/bioinformatics/btad368

Arraud, N., Linares, R., Tan, S., Gounou, C., Pasquet, J., Mornet, S., & Brisson, A. R. (2014). Extracellular vesicles from blood plasma: determination of their morphology, size, phenotype and concentration. Journal Of Thrombosis And Haemostasis, 12(5), 614–627. 10.1111/jth.12554

Bratu, D. P., Cha, B. J., Mhlanga, M. M., Kramer, F. R., & Tyagi, S. (2003). Visualizing the distribution and transport of mRNAs in living cells. Proceedings of the National Academy of Sciences of the United States of America, 100(23), 13308–13313. 10.1073/pnas.2233244100

Brennan, K., Iversen, K. F., Blanco-Fernández, A., Lund, T., Plesner, T., & Mc Gee, M. M. (2022). Extracellular Vesicles Isolated from Plasma of Multiple Myeloma Patients Treated with Daratumumab Express CD38, PD-L1, and the Complement Inhibitory Proteins CD55 and CD59. Cells, 11(21), 3365. 10.3390/cells11213365

Brittain, G.C., Chen, Y.Q., Martinez, E. et al. A Novel Semiconductor-Based Flow Cytometer with Enhanced Light-Scatter Sensitivity for the Analysis of Biological Nanoparticles. Science Reports 9, 16039 (2019). 10.1038/s41598-019-52366-4

Burnie, J., Ouano, C., Costa, V., Castrosín, I., Hammond, C., Matthews, H., Tigges, J., & Corbett-Helaire, K. S. (2026). Evaluating the utility of a nanoscale flow cytometer for detection of surface proteins on HIV and extracellular vesicles. Virology journal, 10.1186/s12985-026-03169-3. Advance online publication. https://doi.org/10.1186/s12985-026-03169-3

Cecchin, R., Troyer, Z., Witwer, K., & Morris, K. V. (2023). Extracellular vesicles: The next generation in gene therapy delivery. Molecular therapy: the journal of the American Society of Gene Therapy, 31(5), 1225–1230. 10.1016/j.ymthe.2023.01.021

Chen, M., Ma, Z., Wu, X., Mao, S., Yang, Y., Tan, J., Krueger, C. J., & Chen, A. K. (2017). A molecular beacon-based approach for live-cell imaging of RNA transcripts with minimal target engineering at the single-molecule level. Scientific reports, 7(1), 1550. 10.1038/s41598-017-01740-1

Choi, D., Montermini, L., Jeong, H., Sharma, S., Meehan, B., & Rak, J. (2019). Mapping Subpopulations of Cancer Cell-Derived Extracellular Vesicles and Particles by Nano-Flow Cytometry. ACS Nano, 13(9), 10499–10511. 10.1021/acsnano.9b04480

Danielson, K. M., Estanislau, J., Tigges, J., Toxavidis, V., Camacho, V., Felton, E. J., Khoory, J., Kreimer, S., Ivanov, A. R., Mantel, P. Y., Jones, J., Akuthota, P., Das, S., & Ghiran, I. (2016). Diurnal Variations of Circulating Extracellular Vesicles Measured by Nano Flow Cytometry. PloS one, 11(1), e0144678. 10.1371/journal.pone.0144678

Försönits, A. I., Tóth, E. Á., Jezsoviczky, S., Bárkai, T., Khamari, D., Galinsoga, A., Királyhidi, P., Kittel, Á., Fazakas, J., Lenzinger, D., Hegyesi, H., Osteikoetxea, X., Visnovitz, T., Pálóczi, K., Bősze, S., & Buzás, E. I. (2025). Improved Accessibility of Extracellular Vesicle Surface Molecules Upon Partial Removal of the Protein Corona by High Ionic Strength. Journal of Extracellular Vesicles, 14(7), e70124. 10.1002/jev2.70124

Geeurickx, E., Tulkens, J., Dhondt, B., Van Deun, J., Lippens, L., De Wever, O., & Hendrix, A. (2019). The generation and use of recombinant extracellular vesicles as biological reference material. Nature Communications 10, 3288 (2019). 10.1038/s41467-019-11182-0

Giesendorf, B. A. J., Vet, J. A. M., Tyagi, S., Mensink, E. J. B. M., Trijbels, F. J. M., & Blom, H. J. (1998). Molecular beacons: A new approach for semiautomated mutation analysis. Clinical Chemistry, 44(3), 482–486. 10.1093/clinchem/44.3.482

Groot Kormelink, T., Arkesteijn, G. J. A., Nauwelaers, F. A., van den Engh, G., Nolte-’t Hoen, E. N. M., & Wauben, M. H. M. (2016). Prerequisites for the analysis and sorting of extracellular vesicle subpopulations by high-resolution flow cytometry. Cytometry Part A, 89(2), 135–147. 10.1002/cyto.a.22644

Higginbotham, J. N., Zhang, Q., Jeppesen, D. K., Scott, A. M., Manning, H. C., Ochieng, J., Franklin, J. L., & Coffey, R. J. (2016). Identification and characterization of EGF receptor in individual exosomes by fluorescence-activated vesicle sorting. Journal of Extracellular Vesicles, 5(1), 29254. 10.3402/jev.v5.29254

Jia, X., Wei, Q., Zhu, Y., & Zhang, W. (2024). Analysis of noise and its characteristics in avalanche photodiode. AIP Advances, 15, Article 095212. 10.1063/5.0229293

Livshits, M. A., Khomyakova, E., Evtushenko, E. G., Lazarev, V. N., Kulemin, N. A., Semina, S. E., Generozov, E. V., & Govorun, V. M. (2015). Isolation of exosomes by differential centrifugation: Theoretical analysis of a commonly used protocol. Scientific Reports 5, 17319 (2015). 10.1038/srep17319

Maqsood, Q., Sumrin, A., Saleem, Y., Wajid, A., & Mahnoor, M. (2024). Exosomes in cancer: Diagnostic and therapeutic applications. Clinical Medicine Insights: Oncology, 18. 10.1177/11795549231215966

Marchisio, M., Simeone, P., Bologna, G., Ercolino, E., Pierdomenico, L., Pieragostino, D., Ventrella, A., Antonini, F., Del Zotto, G., Vergara, D., Celia, C., Di Marzio, L., Del Boccio, P., Fontana, A., Bosco, D., Miscia, S., & Lanuti, P. (2020). Flow Cytometry Analysis of Circulating Extracellular Vesicle Subtypes from Fresh Peripheral Blood Samples. International Journal Of Molecular Sciences, 22(1), 48. 10.3390/ijms22010048

Morales-Kastresana, A., Telford, B., Musich, T.A. et al. Labeling Extracellular Vesicles for Nanoscale Flow Cytometry. Scientific Reports 7, 1878 (2017). 10.1038/s41598-017-01731-2

Morales-Kastresana, A., Musich, T. A., Welsh, J. A., Telford, W., Demberg, T., Wood, J. C. S., Bigos, M., Ross, C. D., Kachynski, A., Dean, A., Felton, E. J., Van Dyke, J., Tigges, J., Toxavidis, V., Parks, D. R., Overton, W. R., Kesarwala, A. H., Freeman, G. J., Rosner, A., Perfetto, S. P.,… Jones, J. C. (2019). High-fidelity detection and sorting of nanoscale vesicles in viral disease and cancer. Journal of Extracellular Vesicles, 8(1), 1597603. 10.1080/20013078.2019.1597603

Morales-Kastresana, A., Welsh, J. A., & Jones, J. C. (2020). Detection and Sorting of Extracellular Vesicles and Viruses Using nanoFACS. Current protocols in cytometry, 95(1), e81. 10.1002/cpcy.81

Oliveira, G. P., Jr, Zigon, E., Rogers, G., Davodian, D., Lu, S., Jovanovic-Talisman, T., Jones, J., Tigges, J., Tyagi, S., & Ghiran, I. C. (2020). Detection of Extracellular Vesicle RNA Using Molecular Beacons. iScience, 23(1), 100782. 10.1016/j.isci.2019.100782

Oliveira Junior, G. P. de, Welsh, J. A., Pinckney, B., Palu, C. C., Lu, S., Zimmerman, A., Barbosa, R. H., Sahu, P., Noshin, M., Gummuluru, S., Tigges, J., Jones, J. C., Ivanov, A. R., & Ghiran, I. C. (2023). Human red blood cells release microvesicles with distinct sizes and protein composition that alter neutrophil phagocytosis. Journal of Extracellular Biology, 2(11), e107. 10.1002/jex2.107

Puryear, W. B., Yu, X., Ramirez, N. P., Reinhard, B. M., & Gummuluru, S. (2012). HIV-1 incorporation of host-cell-derived glycosphingolipid GM3 allows for capture by mature dendritic cells. Proceedings of the National Academy of Sciences of the United States of America, 109(19), 7475–7480. 10.1073/pnas.1201104109

Puryear, W. B., Akiyama, H., Geer, S. D., Ramirez, N. P., Yu, X., Reinhard, B. M., & Gummuluru, S. (2013). Interferon-inducible mechanism of dendritic cell-mediated HIV-1 dissemination is dependent on Siglec-1/CD169. PLoS pathogens, 9(4), e1003291. 10.1371/journal.ppat.1003291

Ramos, E., Dashzeveg, N., Jia, Y., Manu, M., Taftaf, R., Cao, Y., Hoffmann, A., Tsai, C., El-Shennawy, L., Adorno-Cruz, V., Schuster, E., Scholten, D., Patel, D., Liu, X., Patel, P., Wray, B., Zhang, Y., Zhang, S., Mathews, J. V.,… Liu, H.. (2021). CD81 Facilitates Tumor Cell Clustering and Metastasis in Triple Negative Breast Cancer. Social Science Research Network. 10.2139/ssrn.3871398

Ramos, E. K., Tsai, C., Jia, Y., Cao, Y., Manu, M., Taftaf, R., Hoffmann, A. D., El-Shennawy, L., Gritsenko, M. A., Adorno-Cruz, V., Schuster, E. J., Scholten, D., Patel, D., Liu, X., Patel, P., Wray, B., Zhang, Y., Zhang, S., Moore, R. J.,… Liu, H.. (2022). Machine learning-assisted elucidation of CD81–CD44 interactions in promoting cancer stemness and extracellular vesicle integrity. eLife, 11. 10.7554/elife.82669

Renner, T.M., Tang, V.A., Burger, D., Langlois, M. (2020). Intact Viral Particle Counts Measured by Flow Virometry Provide Insight into the Infectivity and Genome Packaging Efficiency of Moloney Murine Leukemia Virus. Journal of Virology 94:10.1128/jvi.01600-19. 10.1128/jvi.01600-19

Ryazantsev, D. Y., Kvach, M. V., Tsybulsky, D. A., Prokhorenko, I. A., Stepanova, I. A., Martynenko, Y. V., Gontarev, S. V., Shmanai, V. V., Zavriev, S. K., & Korshun, V. A. (2014). Design of molecular beacons: 3IZ couple quenchers improve fluorogenic properties of a probe in real-time PCR assay. Analyst, 139, 2867–2872. 10.1039/C4AN00081A

Simeoli, R., Montague, K., Jones, H. R., Castaldi, L., Chambers, D., Kelleher, J. H., Vacca, V., Pitcher, T., Grist, J., Al-Ahdal, H., Wong, L. F., Perretti, M., Lai, J., Mouritzen, P., Heppenstall, P., & Malcangio, M. (2017). Exosomal cargo including microRNA regulates sensory neuron to macrophage communication after nerve trauma. Nature communications, 8(1), 1778. 10.1038/s41467-017-01841-5

Tan, W., Wang, K., & Drake, T. J. (2004). Molecular beacons. Current Opinion in Chemical Biology, 8(5), 547–553. 10.1016/j.cbpa.2004.08.010

Tian, W. J., Huang, M. L., Qin, Q. F., Chen, Q., Fang, K., & Wang, P. L. (2016). Prognostic Impact of Epidermal Growth Factor Receptor Overexpression in Patients with Cervical Cancer: A Meta-Analysis. PloS one, 11(7), e0158787. 10.1371/journal.pone.0158787

Tyagi, S., & Kramer, F. R. (1996). Molecular beacons: Probes that fluoresce upon hybridization. Nature Biotechnology, 14(3), 303–308. 10.1038/nbt0396-303

Vargas, D. Y., Raj, A., Marras, S. A., Kramer, F. R., & Tyagi, S. (2005). Mechanism of mRNA transport in the nucleus. Proceedings of the National Academy of Sciences of the United States of America, 102(47), 17008–17013. 10.1073/pnas.0505580102

Wang, X., Tian, L., Lu, J., & Ng, I. O.-L. (2022). Exosomes and cancer: Diagnostic and prognostic biomarkers and therapeutic vehicle. Oncogenesis, 11, Article 54. 10.1038/s41389-022-00431-5

Welsh, J. A., & Jones, J. C. (2020). Small Particle Fluorescence and Light Scatter Calibration Using FCM PASS Software. Current Protocols In Cytometry, 94(1). 10.1002/cpcy.79

Welsh, J. A., Jones, J. C., & Tang, V. A. (2020). Fluorescence and Light Scatter Calibration Allow Comparisons of Small Particle Data in Standard Units across Different Flow Cytometry Platforms and Detector Settings. Cytometry. Part A : the journal of the International Society for Analytical Cytology, 97(6), 592–601. 10.1002/cyto.a.24029

Welsh, J. A., Arkesteijn, G. J. A., Bremer, M., Cimorelli, M., Dignat-George, F., Giebel, B., Görgens, A., Hendrix, A., Kuiper, M., Lacroix, R., Lannigan, J., Van Leeuwen, T. G., Lozano-Andrés, E., Rao, S., Robert, S., De Rond, L., Tang, V. A., Tertel, T., Yan, X.,… Van Der Pol, E. (2023). A compendium of single extracellular vesicle flow cytometry. Journal Of Extracellular Vesicles, 12(2). 10.1002/jev2.12299

Xin, X., Zeng, X., Gu, H., Li, M., Tan, H., Jin, Z., Hua, T., Shi, R., & Wang, H. (2016). CD147/EMMPRIN overexpression and prognosis in cancer: A systematic review and meta-analysis. Scientific reports, 6, 32804. 10.1038/srep32804

Zhang, J., Wu, J., Wang, G., He, L., Zheng, Z., Wu, M., & Zhang, Y. (2023). Extracellular Vesicles: Techniques and Biomedical Applications Related to Single Vesicle Analysis. ACS nano, 17(18), 17668–17698. 10.1021/acsnano.3c03172

