## Supplemental for "Advancing Nano-Flow Cytometry: High-Precision Sorting and Analysis of Extracellular Vesicles": Castrosín et al- Supplemental.docx

1. **Supplementary Materials**
   1.1. **NIST Traceable beads**

| **Name** | **Catalog Number** | **Certified Mean Diameter** | **Lot Number** | **Standard Deviation** | **Coefficient of Variation** |
| --- | --- | --- | --- | --- | --- |
| 70 nm | 3070A | 70 nm ± 3 nm | 264344 | 7.3 nm | 10.4 % |
| 80 nm | 3080A | 81 nm ± 3 nm | 238296 | 9.5 nm | 11.7 % |
| 90 nm | 3090A | 92 nm ± 3 nm | 238628 | 7 nm | 7.6 % |
| 100 nm | 3100A | 101 ± 3 nm | 261562 | 6.2 nm | 6.1 % |
| 125 nm | 3125A | 122 ± 3 nm | 235037 | 6.8 nm | 5.6 % |
| 150 nm | 3150A | 151 ± 3 nm | 264883 | 5.3 nm | 3.5 % |
| 200 nm | 3200A | 202 ± 4 nm | 260104 | 5.1 nm | 2.5 % |
| 220 nm | 3220A | 221 ± 6 nm | 264288 | 4.7 nm | 2.1 % |
| 240 nm | 3240A | 237 ± 5 nm | 265408 | 4.7 nm | 2.0 % |
| 300 nm | 3300A | 300 ± 5 nm | 267148 | 6.6 nm | 2.2 % |
| 350 nm | 3350A | 345 ± 7 nm | 264514 | 6.5 nm | 1.9 % |
| 400 nm | 3400A | 401 ± 6 nm | 264624 | 5.0 nm | 1.3 % |
| 495 nm | 3495A | 496 ± 8 nm | 263540 | 8.6 nm | 1.7 % |

**Table S1.** Specifications of the NIST Traceable beads used for calibration with FCM_PASS_ of the different cytometers.

1.2. **Flow Cytometry: Instrument Configurations**

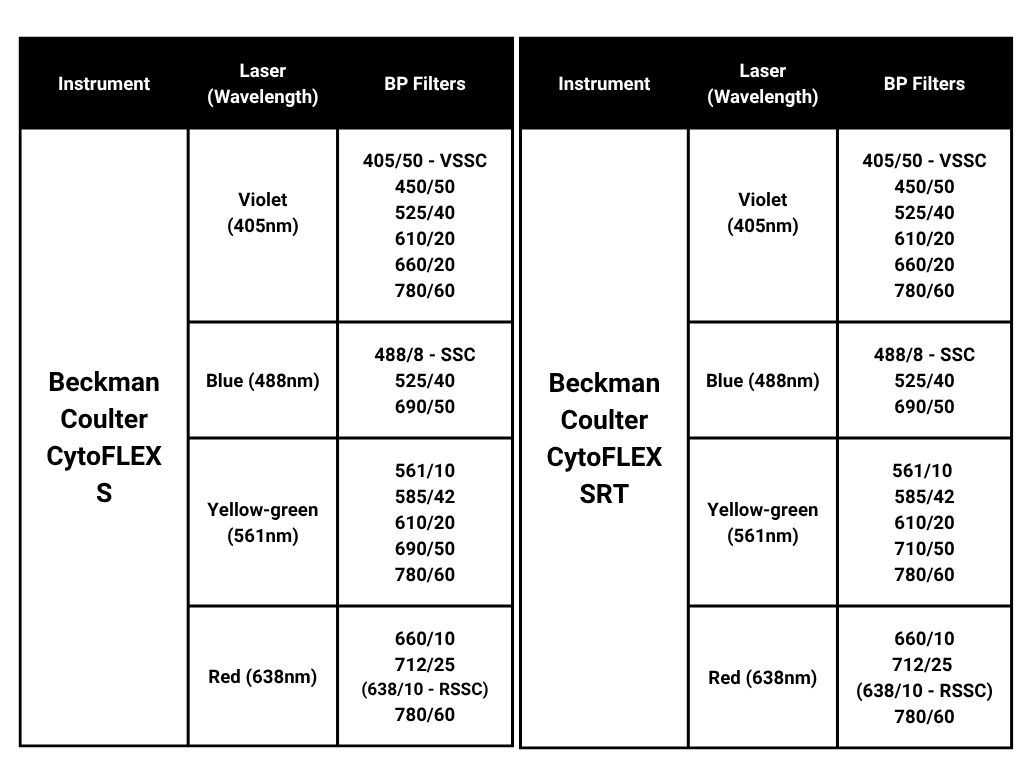

**Table S2**. Specifications of CytoFLEX S and CytoFLEX SRT instruments used for acquisition of the experiments.

1.3. **Supplementary Figures**

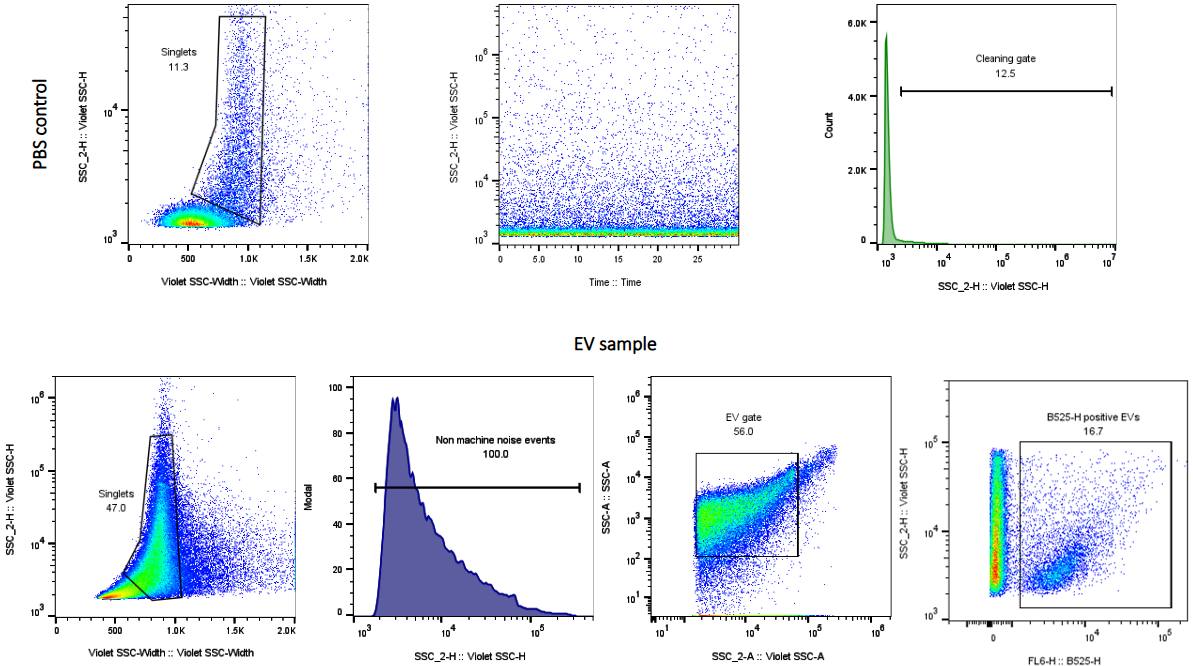

**Figure S1 - Example of gating strategy for EV analysis**

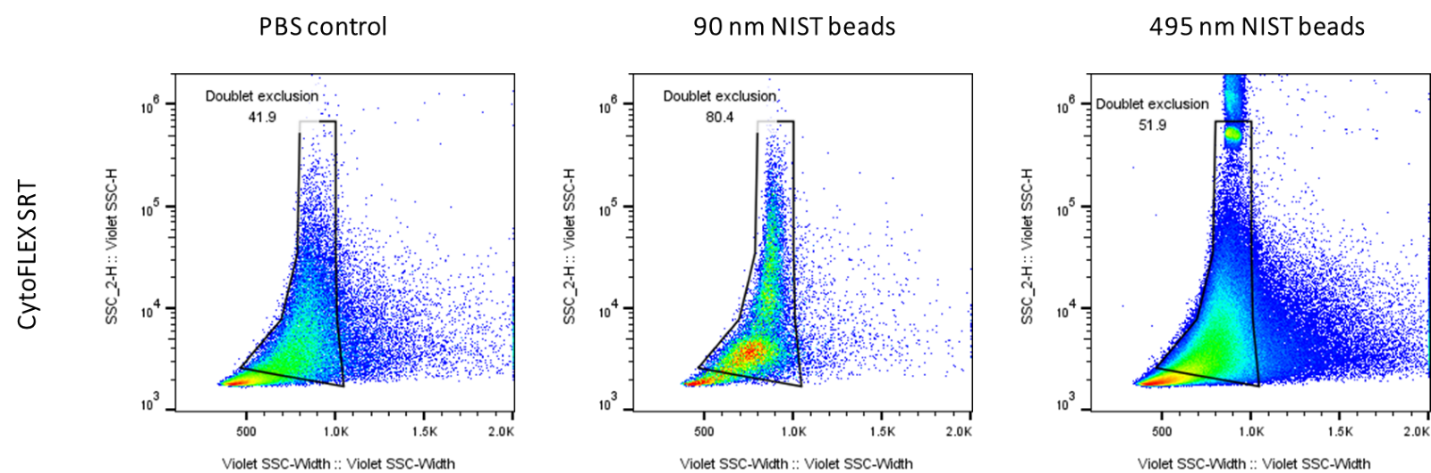

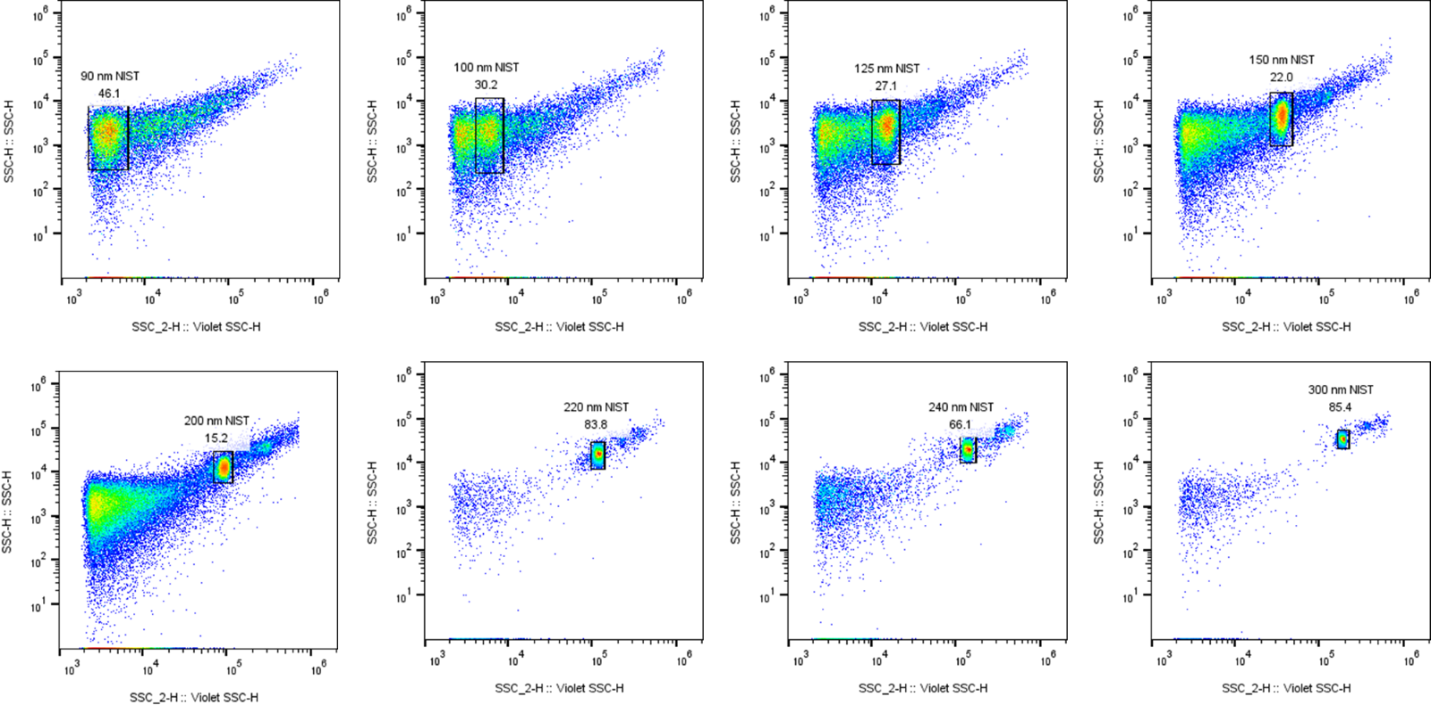

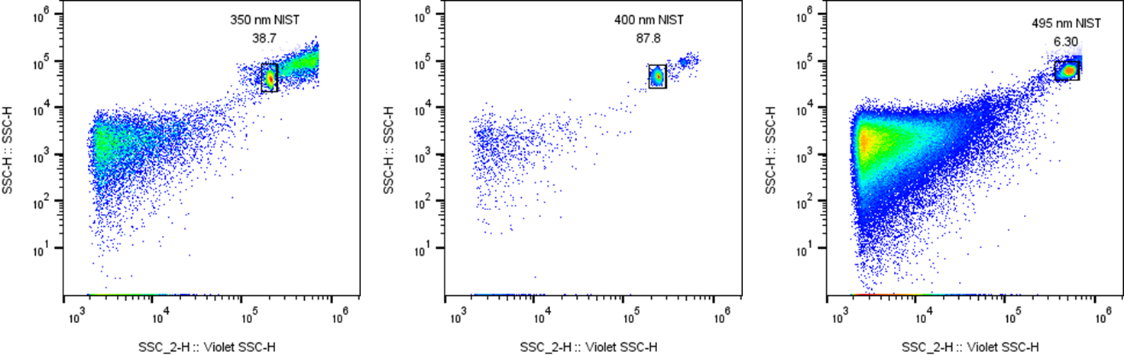

**Figure S2 – Analysis of NIST beads (90, 100, 125, 150, 200, 220, 240, 300, 350, 400 and 495 nm) on the CytoFLEX SRT at UCD**.

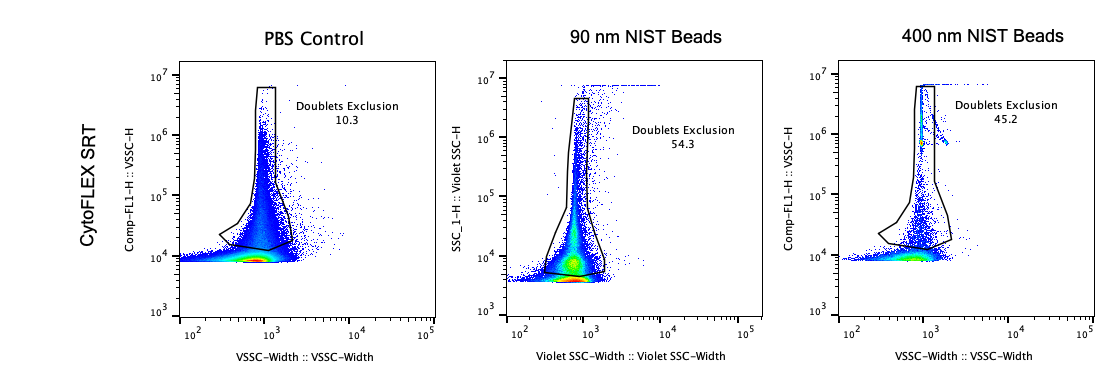

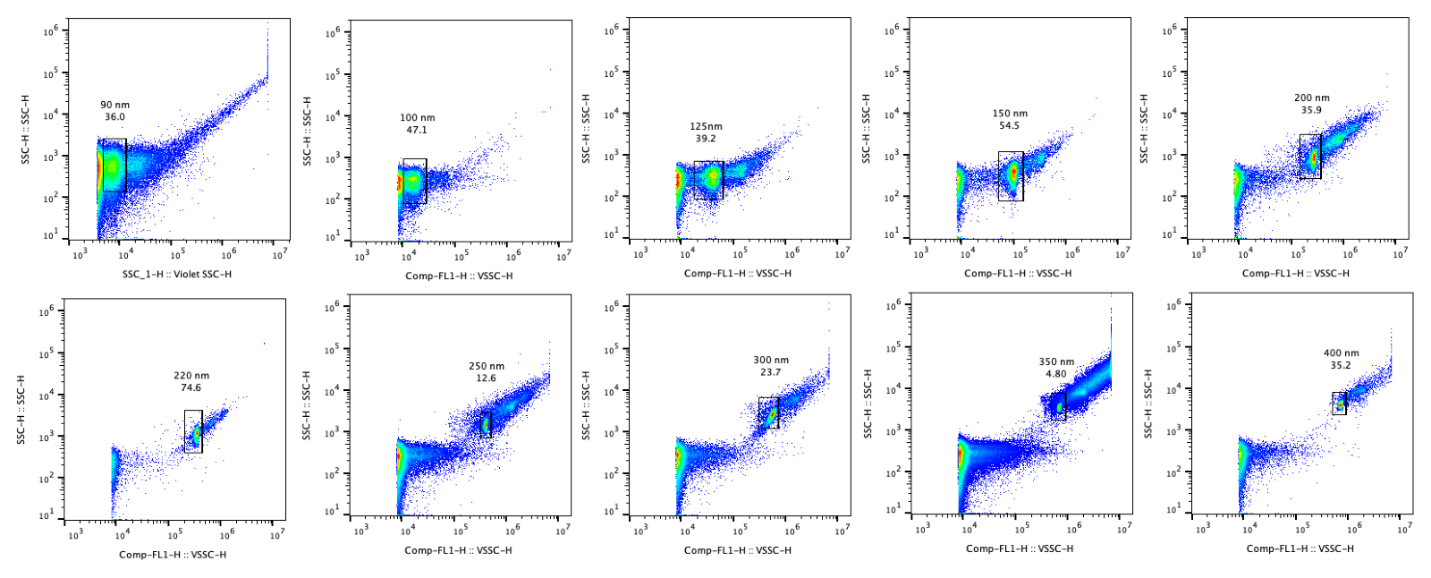

**Figure S3 – Analysis of NIST beads (790, 100, 125, 150, 200, 220, 240, 300, 350, 400 nm) on the CytoFLEX SRT at BIDMC.**

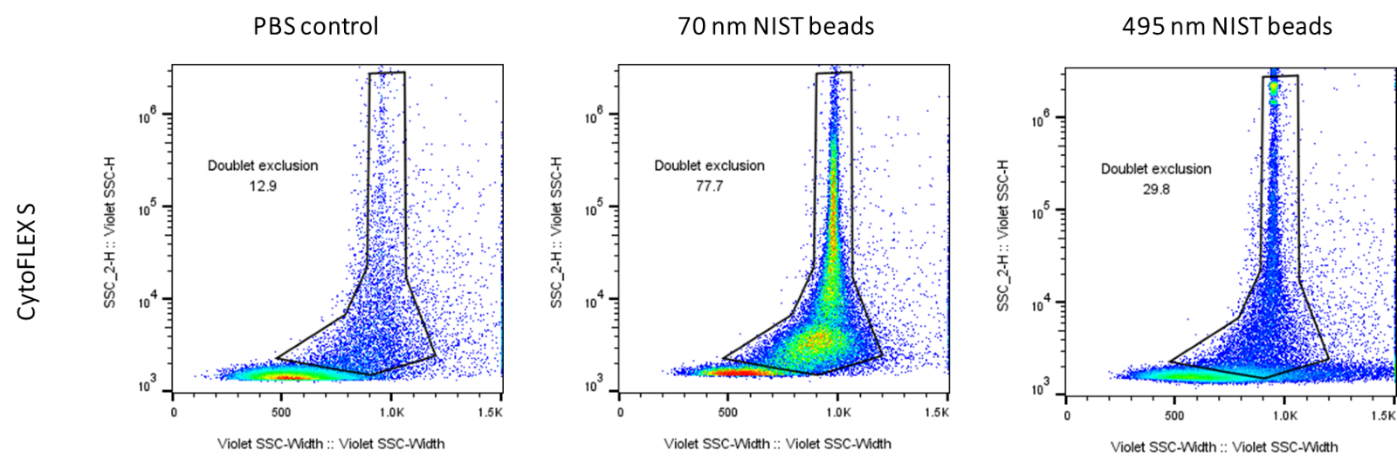

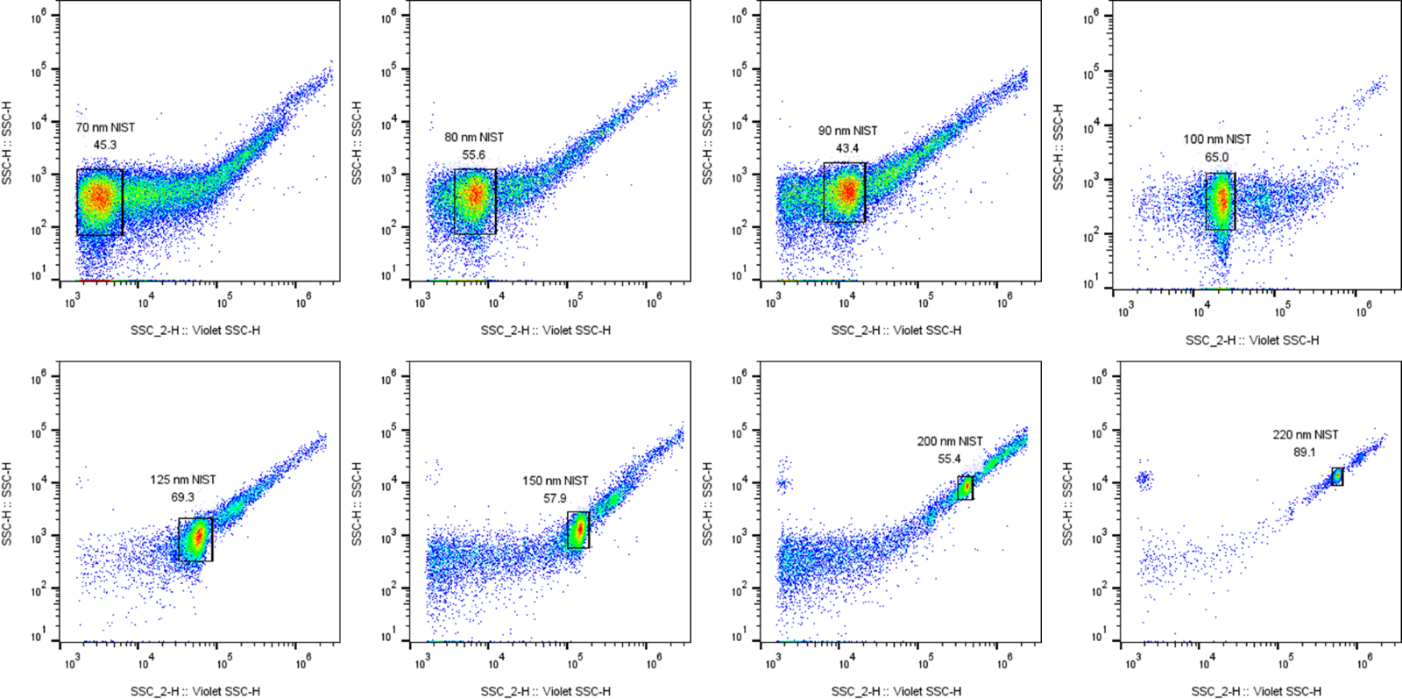

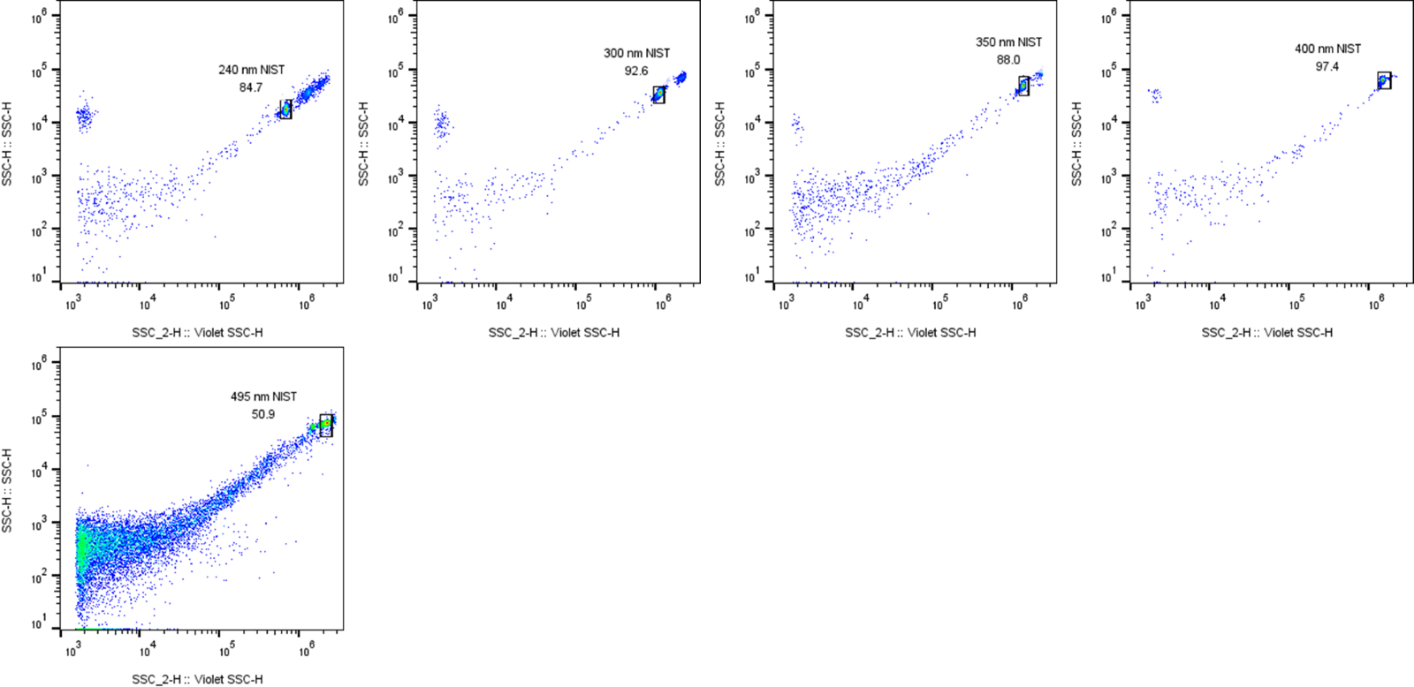

**Figure S4 – Representative analysis of NIST beads (70, 80, 90, 100, 125, 150, 200, 220, 240, 300, 350, 400 and 495 nm) on the CytoFLEX S for both BIDMC and UCD.**

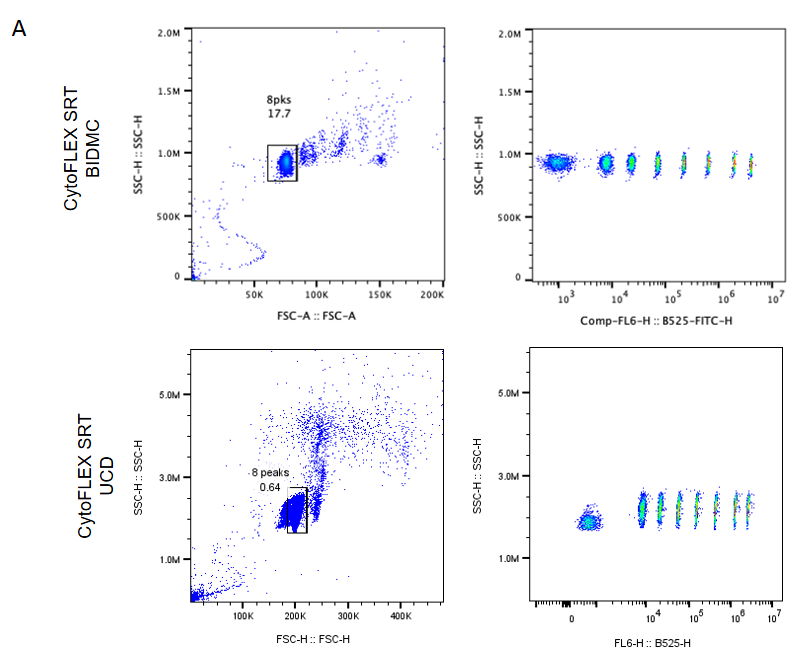

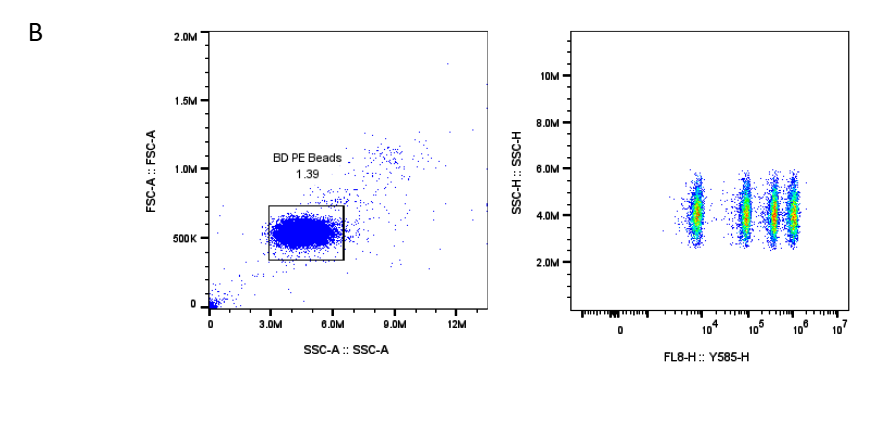

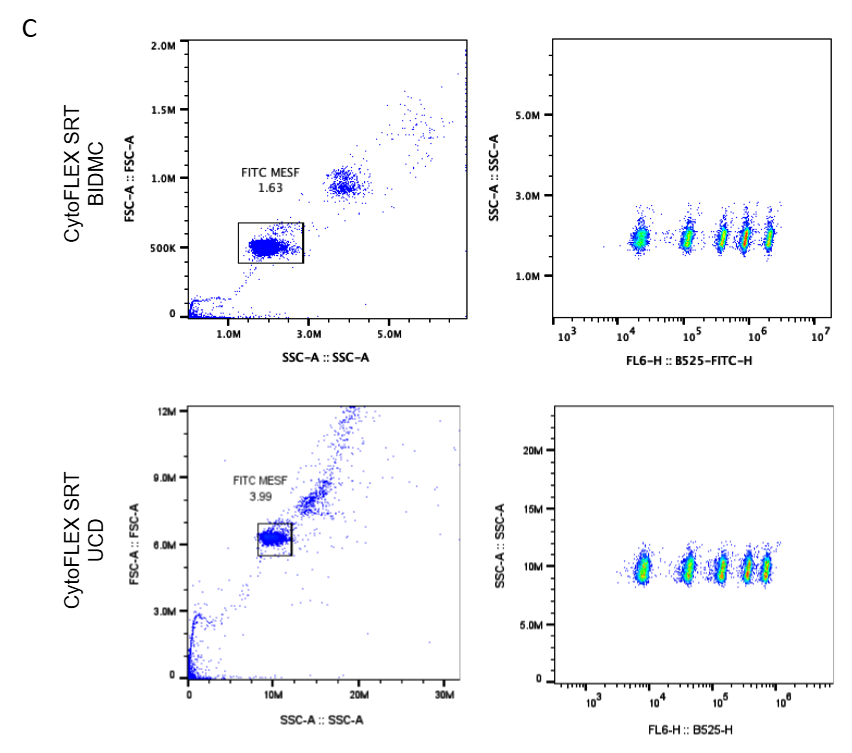

**Figure S5 – Fluorescent beads for FCM_PASS_ calibration**

A) SpheroTech Rainbow Beads analysis. B) Representative analysis of BD Biosciences PE MESF Beads for both BIDMC and UCD. C) FITC MESF Beads analysis.

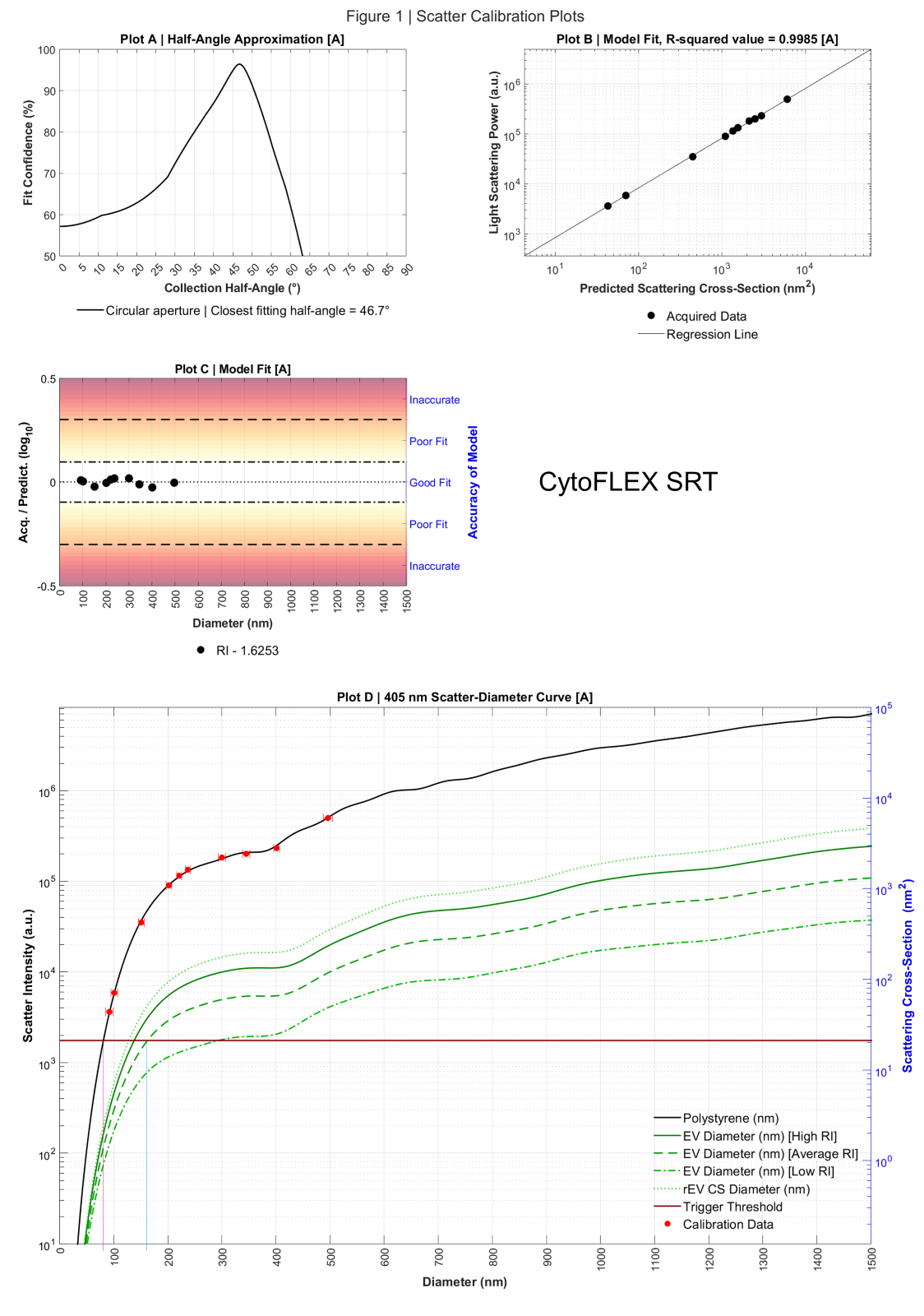

**Figure S6 -** FCM_PASS_ scatter calibration report from UCD’s CytoFLEX SRT.

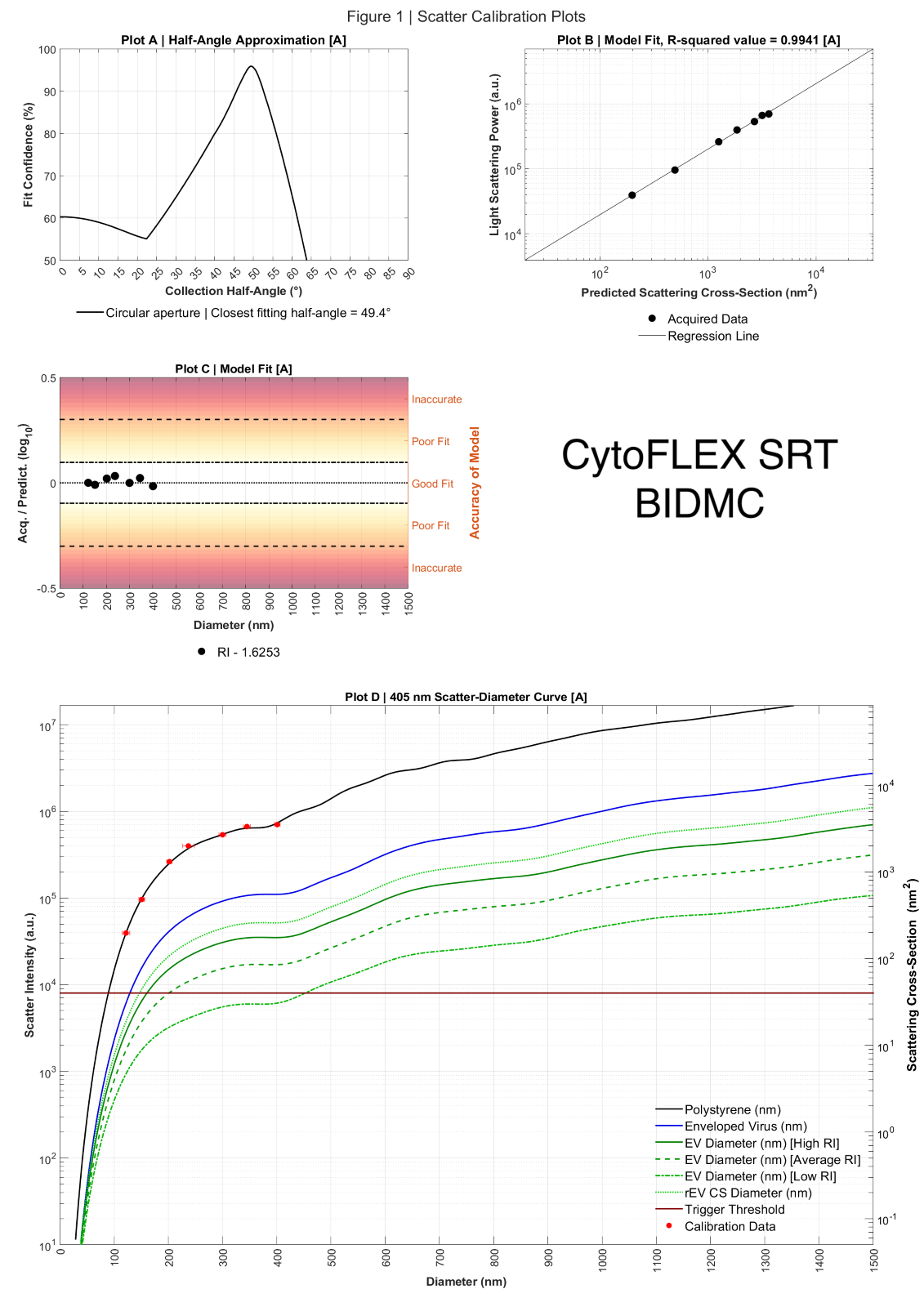

**Figure S7 -** FCM_PASS_ scatter calibration report from BIDMC’s CytoFLEX SRT.

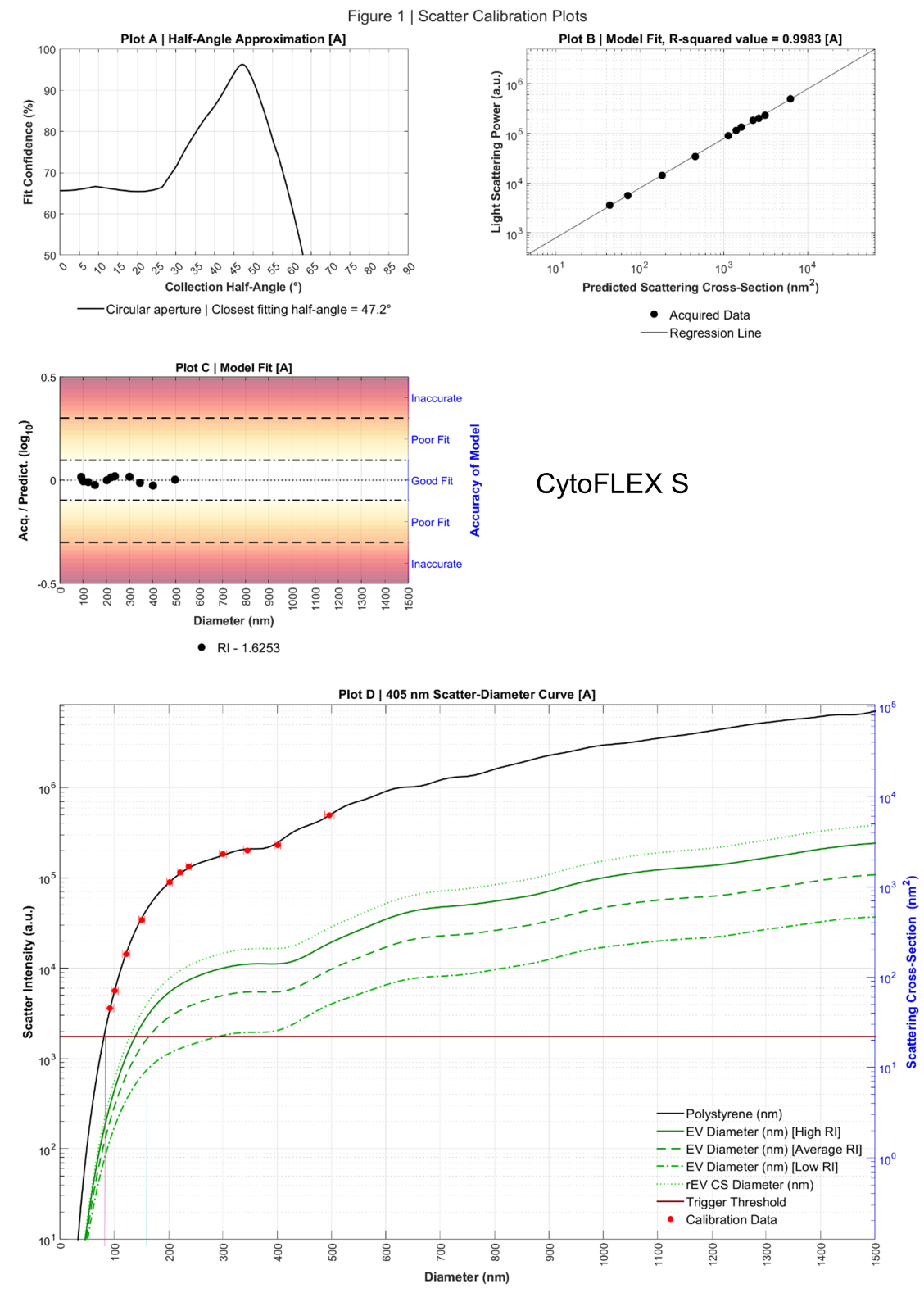

**Figure S8 -** Representative FCM_PASS_ scatter calibration report from a CytoFLEX S.

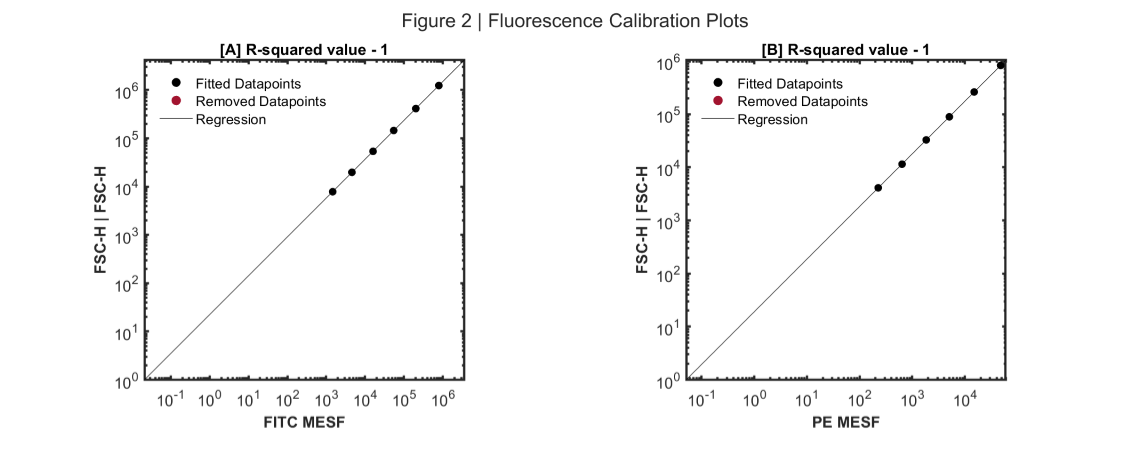

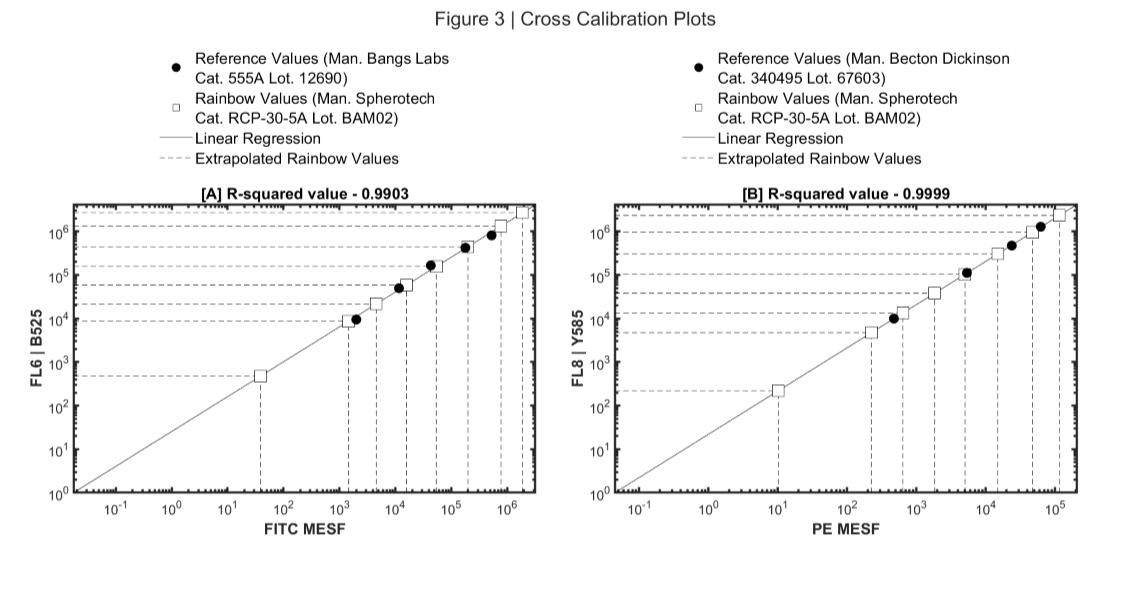

**Figure S9 -** Representative FCM_PASS_ fluorescent calibration and Cross Calibration reports from a CytoFLEX SRT.

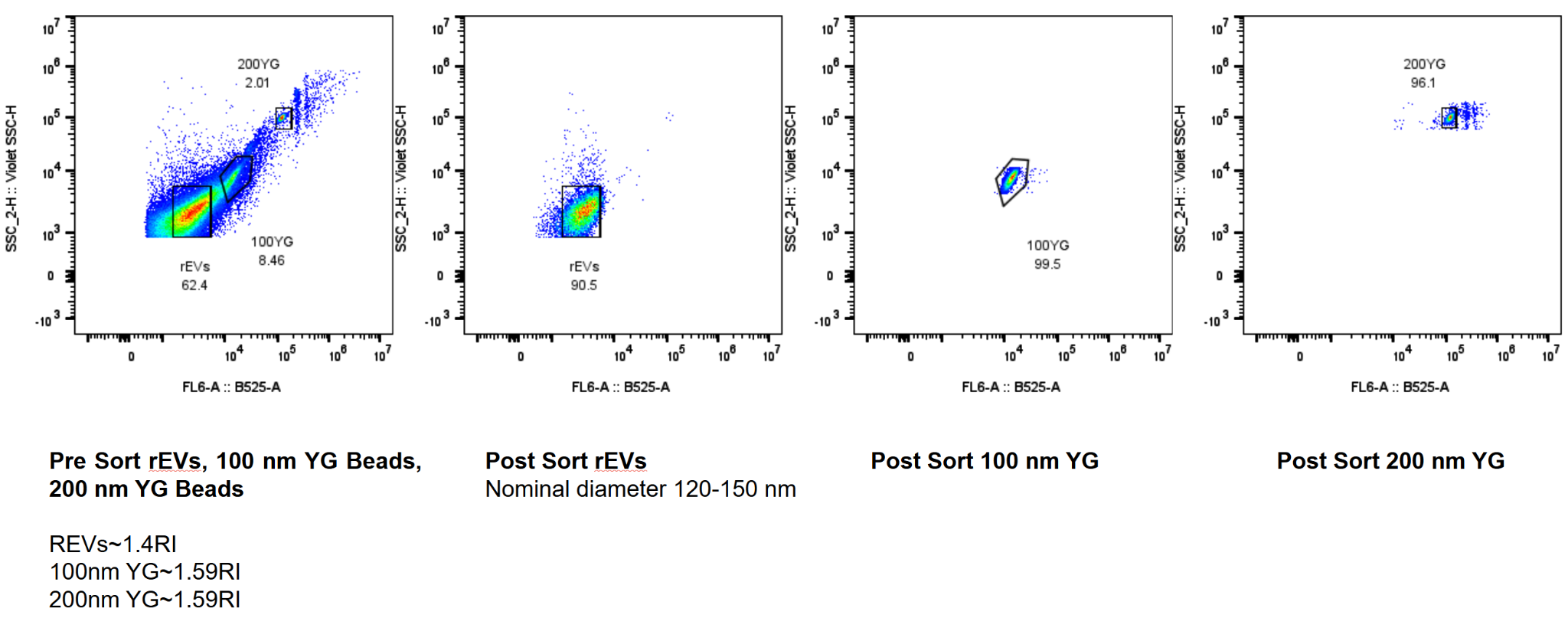

**Figure S10 - Separation based on 525 nm fluorescence: rEVs, 100 YG and 200 YG beads.**

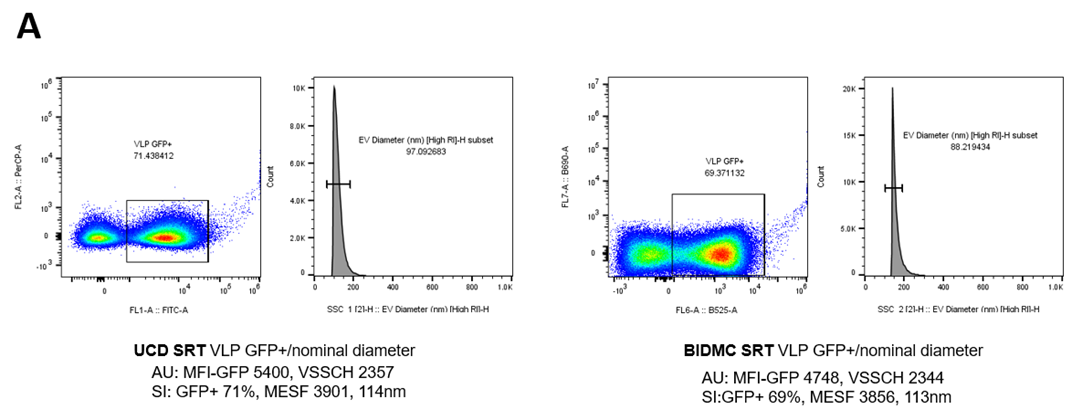

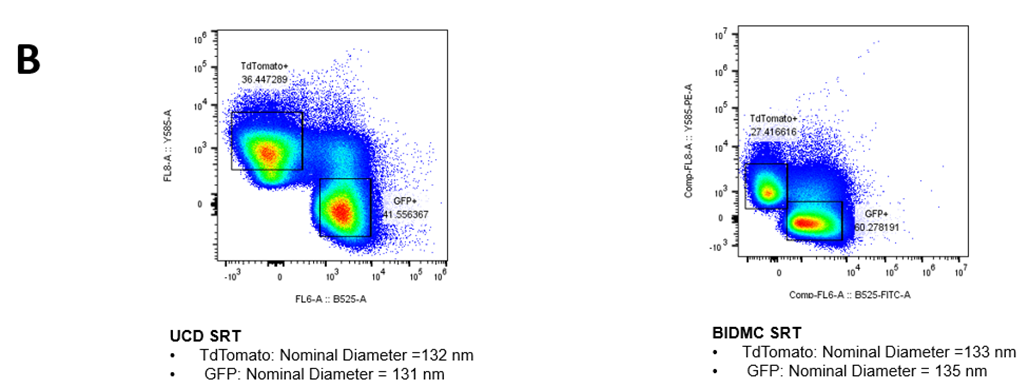

**Figure S11 - Sorting of HIV gag-eGFP/tdTomato VLPs, calibration and standardization via FCM_PASS_ between two institutions: University College of Dublin (UCD) and Beth Israel Deaconess Medical Center (BIDMC).**

A) HIV gag eGFP+ VLPs calibration and analysis via FCM_PASS_ between UCD and BIDMC. FCM_PASS_ software allows calibrating both light scattering and fluorescence signals. Thus, FCM_PASS_ allows the conversion of scattering intensity in arbitrary units (AU) to diameter distribution (in nm), and SI allows for 1:1 comparison of data. B) Comparison of the sortings performed in each institution. Note that the first plot is the sort done at UCD (detailed in Figure 8-B). HIV gag-eGFP VLPs were successfully separated from HIV gag-tdTomato VLPs by using dual thresholding. The calibration of the scattering signal though FCM_PASS_ resulted in a consistent measurement of the nominal diameter for each population. Therefore, FCM_PASS_ facilitates direct comparisons between institutions by standardizing data analysis, leading to improved accuracy and reliability in multi-institutional studies.

**A**

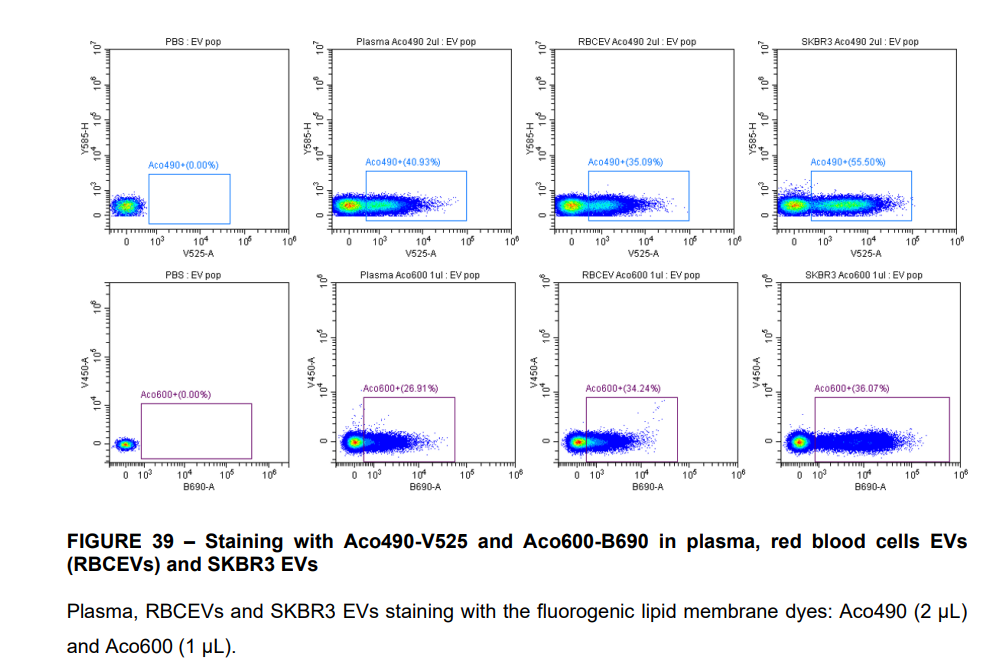

**B**

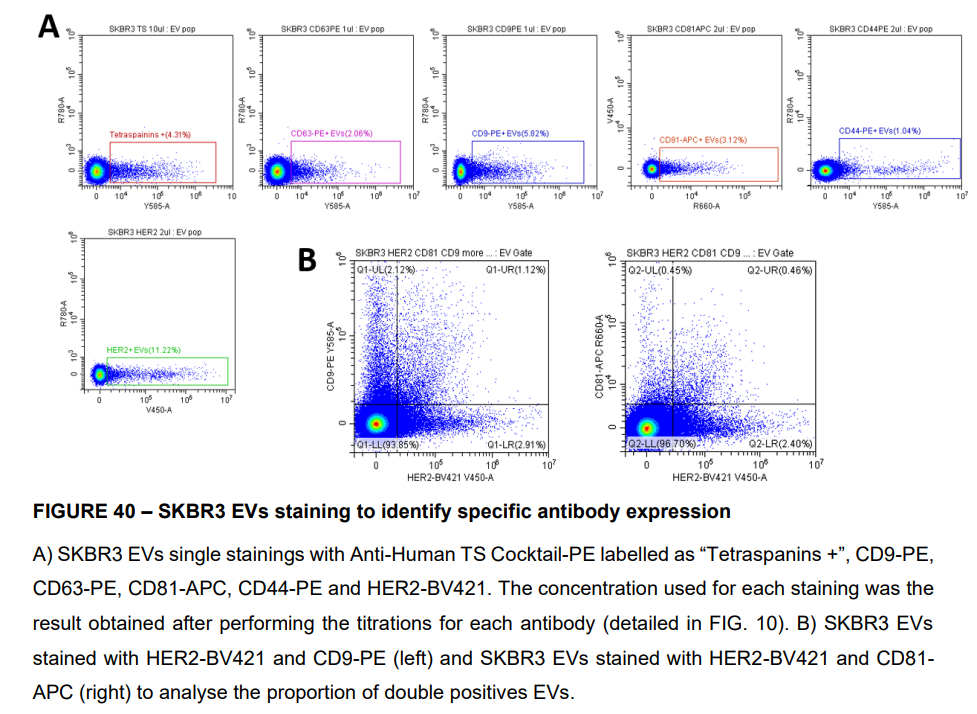

**C**

**Figure S12 - SKBR3 EVs single and double staining analysis**

A) Plasma, RBC.EVs and SKBR3 EVs staining with the fluorogenic lipid membrane dyes: Aco490 (V525+, 2 µL) and Aco600 (B690+,1 µL). B) SKBR3 EVs single staining with Anti-Human TS Cocktail-PE labelled as ‘Tetraspanins +’, CD9-PE, CD63-PE, CD81-APC, CD44-PE and HER2-BV421. The concentration used for each antibody was the result obtained after performing the titrations for each antibody. C) SKBR3 EVs stained with HER2-BV421 and CD9-PE (left) and SKBR3 EVs stained with HER2-BV421 and CD81-APC (right) to determine the proportion of double positives EVs.

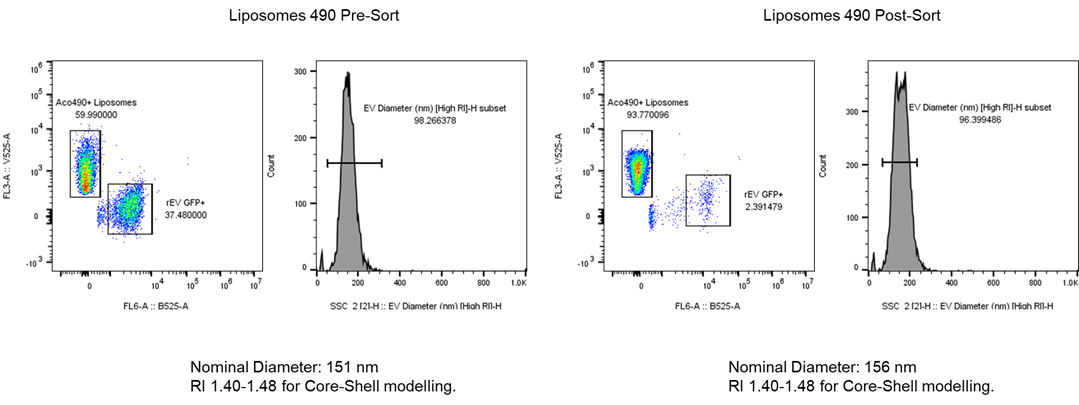
**Figure S13 - Sorting of Aco490+ liposomes based on fluorescence intensity and excitation wavelengths.**

The sample Pre-Sort (left) included Aco490+ liposomes (Ex. 405nm/Em. 510nm) and HIV gag-eGFP VLPs (Ex. 488nm/Em. 525nm). Aco490+ liposomes were sorted using a double threshold (V525/B525) and remained consistent in fluorescence (right). Core-Shell modelling with FCM_PASS_ analysis for 490 Liposomes showed nominal diameters of 151 nm before sorting and 156 nm after sorting, demonstrating that the sorting process has no significant impact on EV size statistics.

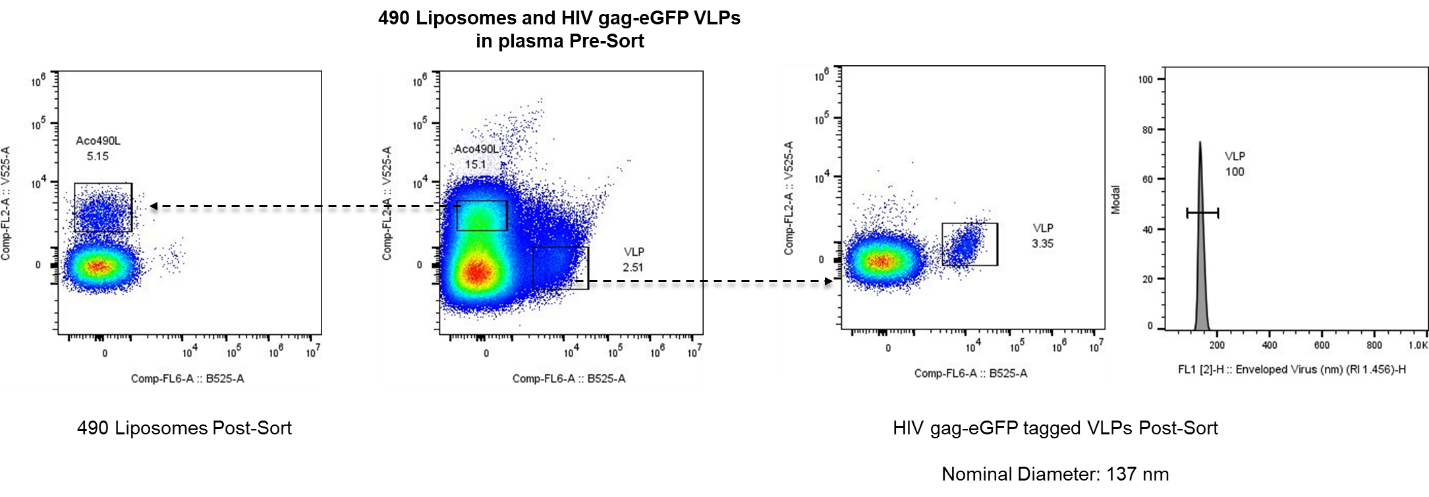
**Figure S14 – 490 Liposomes and HIV gag-eGFP VLPs spiked into plasma**

The spiked 490RL and HIV gag-eGFP VLPs were isolated from the plasma sample using a VSSC threshold. Reanalysis showed an enrichment of each nanoparticle population, with the HIV gag-eGFP VLPs exhibiting a nominal diameter similar to that calculated in previous analyses using the Core-Shell modeling in FCM_PASS_.

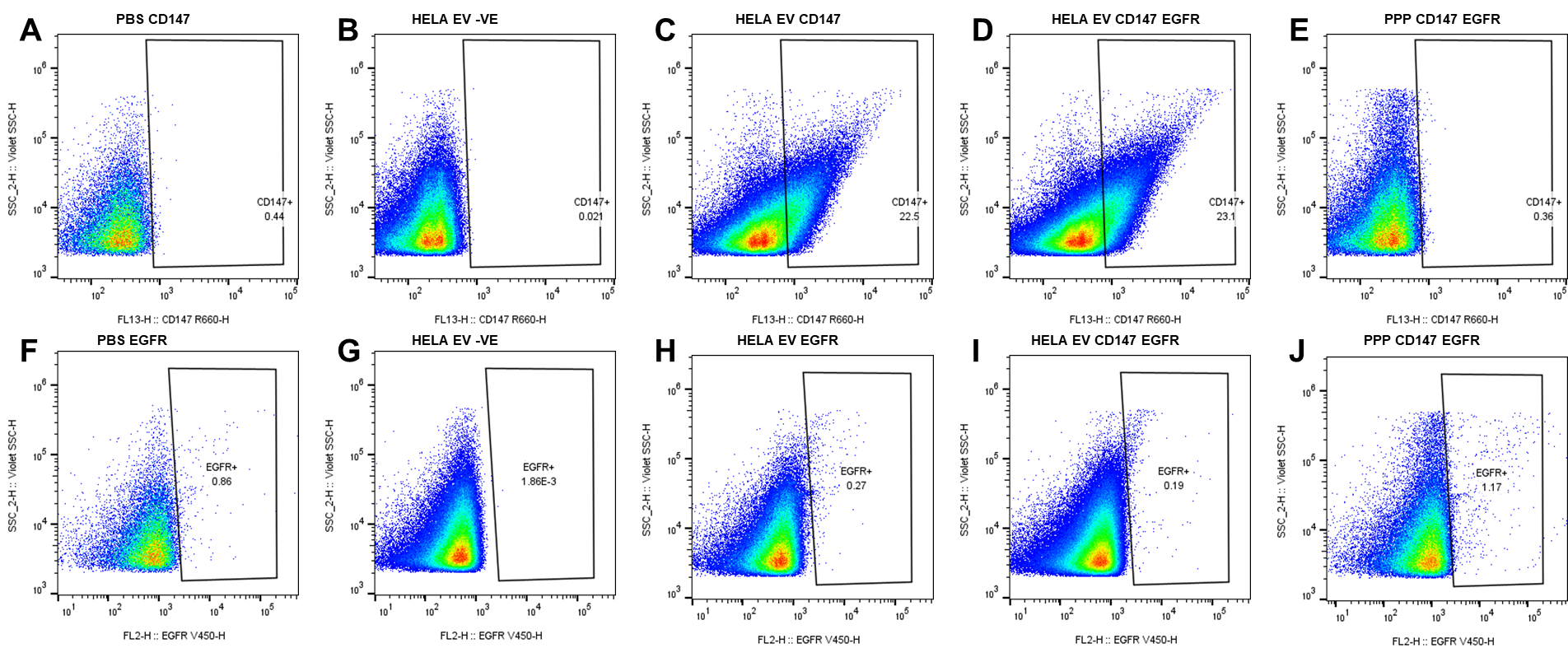

**Figure S15 - HELA EVs single and double staining**

Surface CD147 (A-E) and EGFR (F-J) expression was measured in HELA EVs (B-D, and G-I) and plasma (E and J) incubated with 0.1μl anti-CD147-APC or 0.2 µl anti-EGFR-BV450 for 1h and diluted 1-20 with PBS prior to recording. PBS (A and F) was used as a control to determine the antibody background. CD147+ and EGFR+ gates were set up to identify CD147+ and EGFR+ events. The percentages in the dot plots indicate the number of CD147+ and EGFR+ EVs amongst the total EVs.

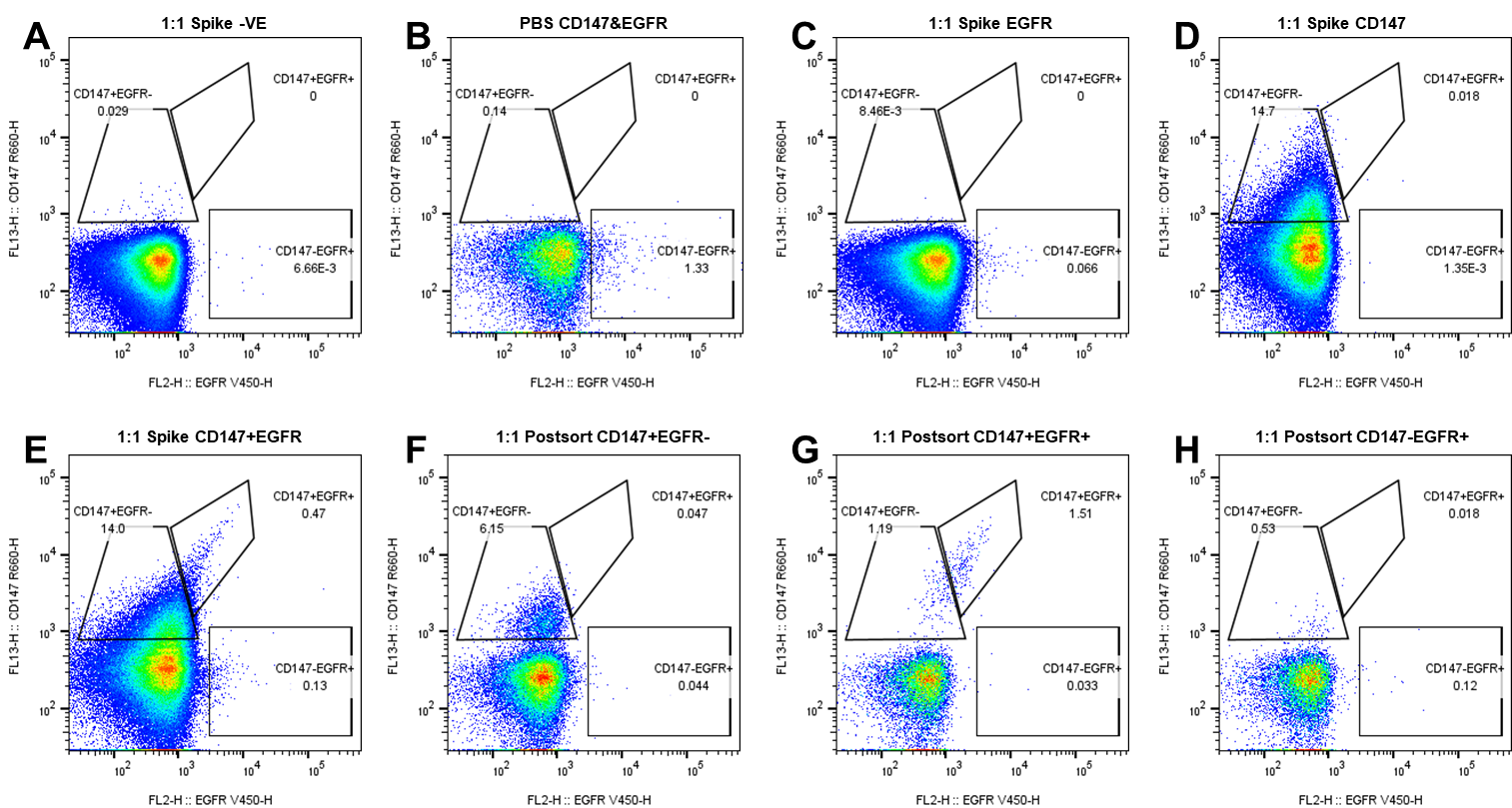

**Figure S16- Sorting of HELA EV spike in from plasma**

CD147+EGFR-, CD147-EGFR+ and CD147+EGFR+ EVs were sorted from plasma spiked with HELA EVs at a 1:1 ratio. Samples were incubated with 0.1μl anti-CD147-APC and 0.2 µl anti-EGFR-BV450 for 1h and diluted 1-20 with PBS prior to recording. Unstained spiked plasma (A) and PBS (B) were used as controls to determine the antibody background. Single and double stained samples (C-E) were used to set up the CD147+EGFR-, CD147-EGFR+ and CD147+EGFR+ gates. Post sort analysis of the 3 sorted populations on R660-H vs V450-H dot plots are shown (F-H), with CD147+EGFR+ double positive EVs (G) losing some of their EGFR-BV450 and CD147-APC fluorescence, so that 44% of CD147+ events are being detected in the CD147+EGFR- gate. The CD147+EGFR- have a lower CD147 intensity than the double positive population (F), meanwhile the CD147-EGFR+ events show a very low abundance and appear heterogeneous (H). The percentages in the dot plots indicate the number of CD147+ and EGFR+ EVs amongst the total EVs.
